# GpsB acts as an adapter for MacP-mediated activation of class A penicillin-binding protein aPBP2a in *Streptococcus pneumoniae*, independently of MacP phosphorylation

**DOI:** 10.64898/2026.06.26.734906

**Authors:** Merrin Joseph, Bohumil Kubeša, Ho-Ching T. Tsui, Mattia Benedet, Orietta Massida, Pavel Branny, Linda Doubravová, Malcolm E. Winkler

**Affiliations:** Department of Biology, Indiana University Bloomington, Bloomington, IN, USA; Institute of Microbiology, Czech Academy of Sciences, Prague, Czech Republic; Center for Medical Sciences (CISMed), Department of Cellular, Computational, and Integrative Biology (CIBIO), University of Trento, Italy

**Keywords:** peptidoglycan (PG) biosynthesis, class A PBP regulation, protein phosphorylation, GpsB binding and function, RocS regulator

## Abstract

Regulation of class A penicillin-binding proteins (aPBPs) in peptidoglycan biosynthesis is incompletely understood in Gram-positive bacteria. One example is activation of aPBP2a by GpsB and phosphorylated MacP in the ovoid-shaped pathogen, *Streptococcus pneumoniae*. We set out to examine whether phosphorylation of Thr residues other than Thr32 contributed to MacP activation of aPBP2a. We also wanted to determine whether GpsB and MacP activation of aPBP2a were related. Here we report that MacP was phosphorylated about equally at Thr32 and Thr56 in physiological and biochemical assays. However, based on transformation and growth assays, phosphorylation of MacP was not required for aPBP2a activation. A structure-function analysis confirmed that most of the MacP cytoplasmic domain, which was predicted by AlphaFold3 to be disordered, was not required for aPBP2a activation. These analyses further identified amino acids in the MacP transmembrane domain and the aPBP2a juxtamembrane region, as well as a variant of the GpsB-binding motif in the membrane-proximal cytoplasmic region of MacP, required for aPBP2a activation. Together, these results support a tripartite model in which GpsB acts as an adapter for activation of aPBP2a by MacP. Finally, additional interaction, Tn-seq, and growth assays suggested other modes of direct or indirect regulation of aPBP2a activity.

## 1. INTRODUCTION

The biosynthesis of the bacterial peptidoglycan (PG) cell wall mesh is carried out by two different kinds of PG synthases. One type of PG synthase contains a polytopic-membrane SEDS (shape, elongation, division, sporulation) protein (Meeske *et al*., 2016). This subunit has glycosyl transferase (GT) activity that synthesizes glycan chains from Lipid II substrate (Rohs & Bernhardt, 2021, Meeske *et al*., 2016). The FtsW or RodA SEDS proteins are essential in most bacteria and carry out glycan synthesis in the divisome and elongasome, respectively (Taguchi *et al*., 2019, Sjodt *et al*., 2018, Emami *et al*., 2017). FtsW and RodA form complexes with different class B penicillin-binding proteins (bPBPs) that have transpeptidase (TP) activity (Cho *et al*., 2016, Egan *et al*., 2020, Sjodt *et al*., 2020). The class B PBP transpeptidase activities crosslink muropeptides to complete the PG mesh (Cho *et al*., 2016). The activities of divisome or elongasome PG synthases are highly regulated, as exemplified by interactions with FtsN or FzlA (Lyu *et al*., 2022, Mahone *et al*., 2024) or MreC and RodZ (Rohs & Bernhardt, 2021, Zhan *et al*., 2026), respectively, to provide spatiotemporal control of cell division and elongation throughout the division cycle.

The second kind of PG synthase is class A PBPs (aPBPs), which contain both GT and TP domains in a single protein (Sauvage *et al*., 2008, Typas *et al*., 2011). There are multiple homologs of aPBPs in most bacteria, often including a pair of redundant aPBPs with a shared essential function that results in synthetic lethality (Straume *et al*., 2021). Class A PBPs play roles in PG fortification and repair of damaged PG (Straume *et al*., 2020, Vigouroux *et al*., 2020, Daitch & Goley, 2020, Dion *et al*., 2019). In addition, apparently redundant aPBPs show different sensitivities to stress conditions, such as low or high pH (Mueller *et al*., 2019), exposure to cell-wall antibiotics (Zapun *et al*., 2008), and osmolarity (Navarro *et al*., 2025). The regulation of *Escherichia coli* (*Eco*) aPBP1A or aPBP1B activity by cognate outer-membrane lipoprotein LpoA or LpoB, respectively, has been well characterized (Pazos and Vollmer, 2021, Straume *et al*., 2021). Strikingly, it was recently reported that LpoB activation of aPBP1B activity occurs by a transient, allosteric coupling mechanism in regions of low PG density (Shlosman *et al*., 2025). In addition, an LpoB-independent activity of one isoform of *Eco* aPBP1B was linked to osmotic protection of the divisome (Navarro *et al*., 2025).

By contrast, examples of regulation of aPBPs in Gram-positive bacteria are only starting to emerge. The first example was discovered for aPBP2a in *Streptococcus pneumoniae* (*Spn*; pneumococcus). In pneumococcus, aPBP2a and aPBP1a are synthetically lethal in that both genes cannot be knocked out simultaneously (Hoskins *et al*., 1999, Paik *et al*., 1999). Therefore, putative positive regulators of aPBP2a activity are identified as by non-essential genes that become synthetically lethal in a Δ*pbp1a* mutant. Two positive regulators, GpsB and MacP, were identified for aPBP2a by using synthetic lethality with Δ*pbp1a* (Rued *et al*., 2017, Cleverley *et al*., 2019, Fenton *et al*., 2018, Midonet *et al*., 2023). GpsB is a hexameric (trimer of dimers), cytoplasmic adapter protein that can link together different proteins, and thereby regulate PG biosynthesis and cell division (Rismondo *et al*., 2016, Cleverley *et al*., 2019, Millat *et al*., 2026). In progenitor strains of *Spn*, GpsB is essential, but its absence can be suppressed by several mutations (Rued *et al*., 2017, Tsui *et al*., 2023). Suppressed Δ*gpsB* was shown to be synthetically lethal with Δ*pbp1a* (Rued *et al*., 2017). Tn-seq screening of a Δ*pbp1a* mutant further revealed MacP as a positive regulator required for aPBP2a activity (Fenton *et al*., 2018), but not of aPBP1a or aPBP1b (**Fig. S2**). Other instances of aPBP regulation have recently been reported in Gram-positive bacteria. In *S. pneumoniae*, aPBP1a activity was shown to be activated by the Ess S-protein (Millat *et al*., 2026, Burnier *et al*., 2026). In *Bacillus subtilis* (*Bsu*), the C-terminal intrinsically disordered region (IDR) of PBP1 was found to sense PG damage under some conditions (Brunet *et al*., 2022). In addition, the accessory *B. subtilis* protein RpdA, which lacks a homolog in *S. pneumoniae*, was reported to be required for PBP4 localization and activity (Huang *et al*., 2025).

The *macP* gene appears to be in an operon between two genes encoding metabolic enzymes, one of which is upstream of *rocS*, which encodes a chromosome partition regulator (**Fig. S1A**) (Demuysere *et al*., 2024, Mercy *et al*., 2019). MacP is a bitopic membrane protein with a predicted structure consisting of a very short ≈4-amino-acid (M101-L104) extracellular domain (Midonet *et al*., 2023), a 23-amino-acid (V78-A100) alpha-helical transmembrane (TM) domain (based on TMHMM (Sonnhammer *et al*., 1998)), a short 13-amino-acid (I65-N77) cytoplasmic alpha-helical domain (based on AlphaFold3 (Abramson *et al*., 2024)), and a remaining 76-amino acid (M1-R76) cytoplasmic domain that is predicted by AlphaFold3 to be remarkably disordered, except for a short 8-amino-acid (D8-N15) stretch of alpha helix. (**Fig. 1A** and **S1B**-**S1D**). Consistent with Tn-seq results suggesting that MacP is a positive regulator of aPBP2a activity, MacP was shown to interact with aPBP2a (Fenton *et al*., 2018). aPBP2a is also a bitopic membrane protein with an overall structure similar to other class A PBPs (**Fig. S1C**). aPBP2a contains a short, largely disordered N-terminal domain containing a binding site (<u>RRS</u>RSD<u>RK</u>) for the N-terminal domains of GpsB (Cleverley *et al*., 2019) (**Fig. S1D**). The extracellular GT domain occurs immediately after the TM domain (57-76 based on TMHMM (Sonnhammer *et al*., 1998)) and is followed by a putative regulatory linker domain and then the TP domain (**Fig. S1B**-**S1D**) (Midonet *et al*., 2023). AlphaFold3 predicts that the carboxy-terminal domain of aPBP2a may fold into an alpha-helical domain **(Fig. S1C**), unlike the intrinsically disordered regions (IDRs) at the C-termini of many Gram-positive aPBPs, including *Bsu* PBP1 (Brunet *et al*., 2022) and *Spn* aPBP1a and aPBP1b (Lamanna *et al*., 2022, Rahman *et al*., 2025).

**Fig. 1.**
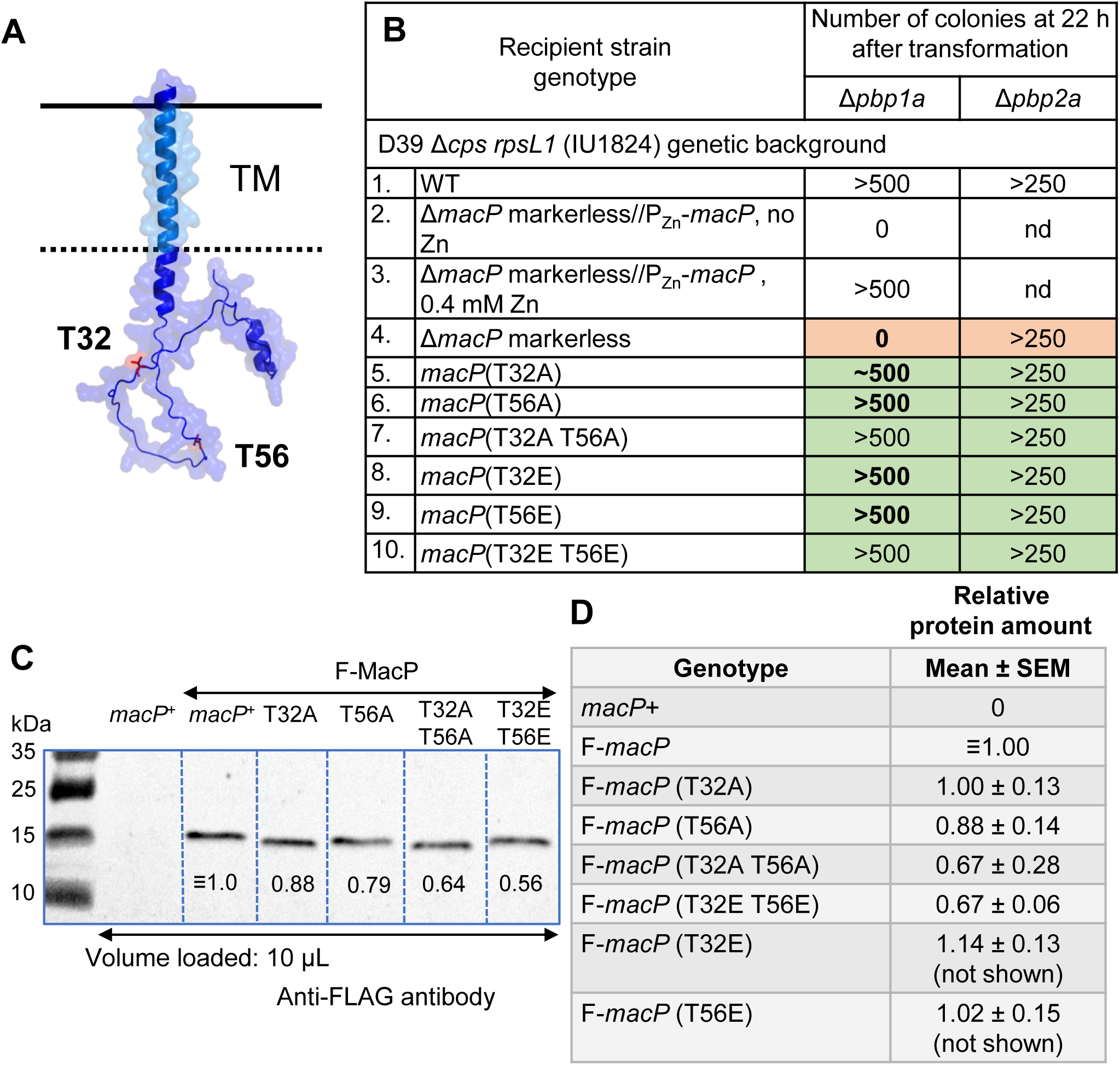
MacP phosphoablative mutations are not synthetically lethal with Δ*pbp1a*. **(A)** AlphaFold3 predicted structure of MacP(*Spn*) with T32 and T56 residues indicated by red. The N-terminal domain of MacP was predicted to be largely unstructured, resulting in a low (0.29) pTM score. **(B)** Transformation assays of the indicated strains with a Δ*pbp1a*::P_c_-*erm* test or a Δ*pbp2a*::P_c_-*erm* control amplicon as described in *Experimental procedures.* Number of colonies and their morphologies on TSAII-BA transformation plates containing erythromycin were recorded after 20h of incubation at 37°C in an atmosphere of 5% CO_2_. Zn indicates (0.4 mM Zn^2+^ + 0.04 mM Mn^2+^) and was present (or absent) from the transformation plates where indicated. All colonies displayed WT morphology. Transformations with a second Δ*pbp1b*::P_c_-*erm* control amplicon gave >500 colonies with WT morphology (data not shown). Similar results were obtained in 2 or more biological replicates. Strains used: WT (IU1824); Δ*macP* markerless//P_Zn_-*macP* (IU15084); Δ*macP* markerless (IU14699); *macP*(T32A) (IU14823); *macP*(T56A) (IU16618); *macP*(T32A T56A) (IU16788); *macP*(T32E) (IU14821); *macP*(T56E) (IU16616); and *macP*(T32E T56E) (IU16787). **(C)** Representative western blot showing relative cellular amounts of F-*macP* and F-*macP* mutant proteins in the strains indicated below were performed as described in *Experimental procedures*. 10 µL of protein lysate was loaded into each lane, and protein amount in each lane was normalized by staining. The predicted molecular mass of F-MacP was 13 kDa. Numbers below bands indicate relative F-MacP amounts in this blot. **(D)** Summary table (means ± SEM) of relative protein amounts from quantitative western blots for three biological replicates performed as described in *Experimental procedures* and illustrated in **Fig. S3E**. Strains used: *macP^+^* (IU1824); F-*macP* (IU17032); F-*macP* (T32A) (IU16978); F-*macP* (T56A) (IU16980); F-*macP* (T32A T56A) (IU17035); F-*macP* (T32E T56E) (IU18342); F-*macP* (T32E) (IU18354); and F-*macP* (T56E) (IU18356). The blot is not shown for F-*macP*(T32E) (IU18354) and F-*macP*(T56E) (IU18356).

Previously, it was shown that the TM domain of MacP is required for midcell localization of MacP, independently of aPBP2a (Fenton *et al*., 2018). Furthermore, MacP was shown to be phosphorylated at Thr32 by the StkP Ser/Thr protein kinase of *Spn* (Fenton *et al*., 2018). A *macP*(T32A) phosphoablative mutation was synthetically lethal upon depletion of aPBP1a, suggesting that phosphorylation of MacP(T32) was required for activation of aPBP2a activity under the conditions tested (Fenton *et al*., 2018). However, a *macP*(T32E) phosphomimetic mutation was also synthetically lethal upon depletion of aPBP1a, and a suppressed Δ*stkP* deletion was not synthetically lethal with aPBP1a depletion, as predicted if phosphorylation of MacP were obligatorily required for aPBP2a activation (Fenton *et al*., 2018). These unexpected results suggested that aPBP2a may be subjected to additional forms of regulation. A recent paper identified suppressor mutations in *pbp2a* that bypass the requirement for MacP activation (Midonet *et al*., 2023). These suppressors included amino-acid changes in the conformationally flexible “jaw” domain thought to activate aPBP2a GT activity and in the putative regulatory linker domain between the GT and TP domains (**Fig. S1B**-**S1D**). The locations of these suppressor amino-acid changes and AlphaFold3 modeling suggested that MacP stimulates aPBP2a GT activity (Midonet *et al*., 2023).

In this study, we set out to further determine the roles of GpsB and MacP phosphorylation in the activation of aPBP2a. Recently, GpsB was shown biochemically to positively activate StkP phosphorylation of target proteins (Stauberová *et al*., 2024); thereby, confirming earlier physiological studies (Fleurie *et al*., 2014, Rued *et al*., 2017). This finding suggested the simple model that GpsB activation of aPBP2a was due to GpsB stimulation of StkP phosphorylation of MacP. However, it was also reported that GpsB interacted directly with MacP (Stauberová *et al*., 2024). Moreover, recent phosphoproteomic studies demonstrated that MacP is phosphorylated on Thr residues besides Thr32 (Hirschfeld *et al*., 2019, Ulrych *et al*., 2021), suggesting other possible levels of aPBP2a regulation.

Here we report that MacP was phosphorylated at Thr32 and Thr56 by the StkP Ser/Thr protein kinase. However, standard transformation and growth assays did not support a requirement for MacP phosphorylation for aPBP2a activation in pneumococcal cells. This unexpected finding was corroborated by structure-function analyses of MacP showing that most of the unstructured cytoplasmic domain of MacP, including Thr32 and Thr56, was not required for aPBP2a activation. Structure-function analyses further identified amino acids that potentially interact in the TM domain of MacP and the juxtamembrane region of aPBP2a. These studies also revealed a variant of the Arg-rich GpsB-binding motif in MacP that is required for aPBP2a activation. Together, these results support a model in which GpsB acts as an adapter that binds separately to both aPBP2a and MacP and promotes optimal activation of aPBP2a by MacP. They also suggest variability in the GpsB-binding motif that may account for binding of GpsB to other PG biosynthesis and cell-division proteins. Finally, MacP interaction assays, additional Tn-seq experiments, and growth assays suggested other modes of direct or indirect regulation of aPBP2a activity, including a link to RocS and the phosphorylation state of proteins other than MacP.

## 2. RESULTS

### 2.1 MacP is phosphorylated at Thr32 and Thr56 by the StkP Ser/Thr kinase, but MacP phosphorylation is not required for aPBP2a activation

An early *Spn* phosphoproteomic study indicated that Thr32 is phosphorylated in MacP (Sun *et al*., 2010) (**Fig. 1A**). Fenton et al. confirmed that MacP(Thr32) is phosphorylated by the StkP Ser/Thr kinase, and their results suggested that MacP(Thr32) phosphorylation was required for positive regulation of aPBP2a activity in the assays used (Fenton *et al*., 2018). A later phosphoproteomic study reported MacP(Thr56) phosphorylation by the StkP Ser/Thr kinase (Hirschfeld *et al*., 2019). Phosphorylation of MacP at both Thr32 and Thr56 in MacP was further confirmed by a recent phosphoproteomic study of exponentially growing *Spn* cells (Ulrych *et al*., 2021). In addition, MacP was found to be phosphorylated at additional residues Thr7 and Thr37 in laboratory strain Rx1 grown stressed with ampicillin (Ulrych *et al*., 2021).

Given these diverse findings, we revisited the topic of MacP phosphorylation in exponentially growing cultures of a derivative of *Spn* type-2 progenitor strain D39W (Slager *et al*., 2018, Lanie *et al*., 2007). We constructed chromosomal *macP* mutations that changed Thr32, or Thr56, or both to phosphoablative Ala (A) or phosphomimetic Glu (E) residues (**Fig. 1**; **Table S1** and **S2**). The single and double phosphoablative and phosphomimetic Thr32/Thr56 *macP* mutants grew in BHI broth with a similar growth rate and yield (**Fig S3A**) and cell morphology (**Fig. S3B**; summarized in **Fig. S4A**) to those of the WT *macP^+^* parent or Δ*macP* mutant. Antibiotic-resistance cassettes inserted into *macP* grew slower and to a lower growth yield than markerless Δ*macP* mutants and were not used in most of this study (**Fig. S3C**). This growth defect was likely due to polarity, consistent with *macP* being in a multigene operon (**Fig. S1A**).

Transformation assays confirmed that Δ*macP* was synthetically lethal with Δ*pbp1a* (lines 2 and 4, **Fig. 1B** and **S2**) (Fenton *et al*., 2018) and that this synthetic lethality was complemented by ectopic expression of *macP*^+^ (line 3, **Fig. 1B**). Unexpectedly, the *macP*(T32A), *macP*(T56A), and *macP*(T32A T56A) phosphoablative mutations were not synthetically lethal with Δ*pbp1a* in transformation assays (lines 5-7, **Fig. 1B**), where the number and morphology of colonies obtained were similar to those of the WT strain. These Δ*pbp1a* transformants grew similarly to the *pbp1a^+^*strains (**Fig. S3A** and **S3D**). These results did not support the previous conclusion that phosphorylation of MacP(T32) (and now MacP(T56)) is required for activation of aPBP2a (Fenton *et al*., 2018).

To determine the phosphorylation state of MacP and relative amounts of MacP mutant variants, we constructed FLAG (F)-tagged fusions (**Table S1** and **S2**). Transformation of the F-MacP^+^, F-MacP(T32A), F-MacP(T56A), and F-MacP(T32A T56A) strains with Δ*pbp1a* resulted in approximately the same number of colonies as the untagged strains (lines 5-10, **Fig. 1B**; data not shown), and these transformants grew similarly to the WT and untagged strains (**Fig. S3D**; data not shown). Together, these results indicated that the F-MacP fusion constructs were functional. The cellular amount of F-MacP(T32A or E) or F-MacP(T56A or E) was approximately the same as F-MacP^+^ in exponentially growing cells, while the amount of F-MacP(T32A T56A) or F-MacP(T32E T56E) was about 70% of the WT amount (**Fig. 1C**, **1D**, **S3E**, and **S4A**). Finally, co-IP and B2H assays demonstrated that phosphoablative Thr to Ala or phosphomimetic Thr to Glu changes did not change the apparent strength of the MacP interaction with aPBP2a or GpsB (**Fig. S3F, S3G, S4A,** and **S5A**).

F-MacP^+^, F-MacP(T32A), F-MacP(T56A), and F-MacP(T32A T56A) were concentrated from cell lysates by binding to anti-FLAG magnetic beads, and degree of phosphorylation was determined by Phos-tag western blotting. In WT F-MacP^+^, we observed two prominent bands and two faint bands (upper panel, **Fig. 2A**). Comparison to the MacP phosphoablative mutants indicated that the lower prominent band in the MacP^+^ sample was unphosphorylated MacP (≈36%) (lower panel, **Fig. 2A**), the upper prominent band was doubly phosphorylated at Thr32 and Thr56 (≈41%), the lower faint band was singly phosphorylated at Thr32 or Thr56 (≈10% each), and the upper faint band contained additional phosphorylation, likely at Thr7 or Thr37 (≈13%) as previously detected in stressed cells (Ulrych *et al*., 2021). Consistent with the Phos-tag analysis, nearly equal amounts of T32 and T56 phosphorylation were detected in immunoblots of F-MacP variants probed with-anti-pThr antibody (**Fig. S3H**).

**Fig. 2.**
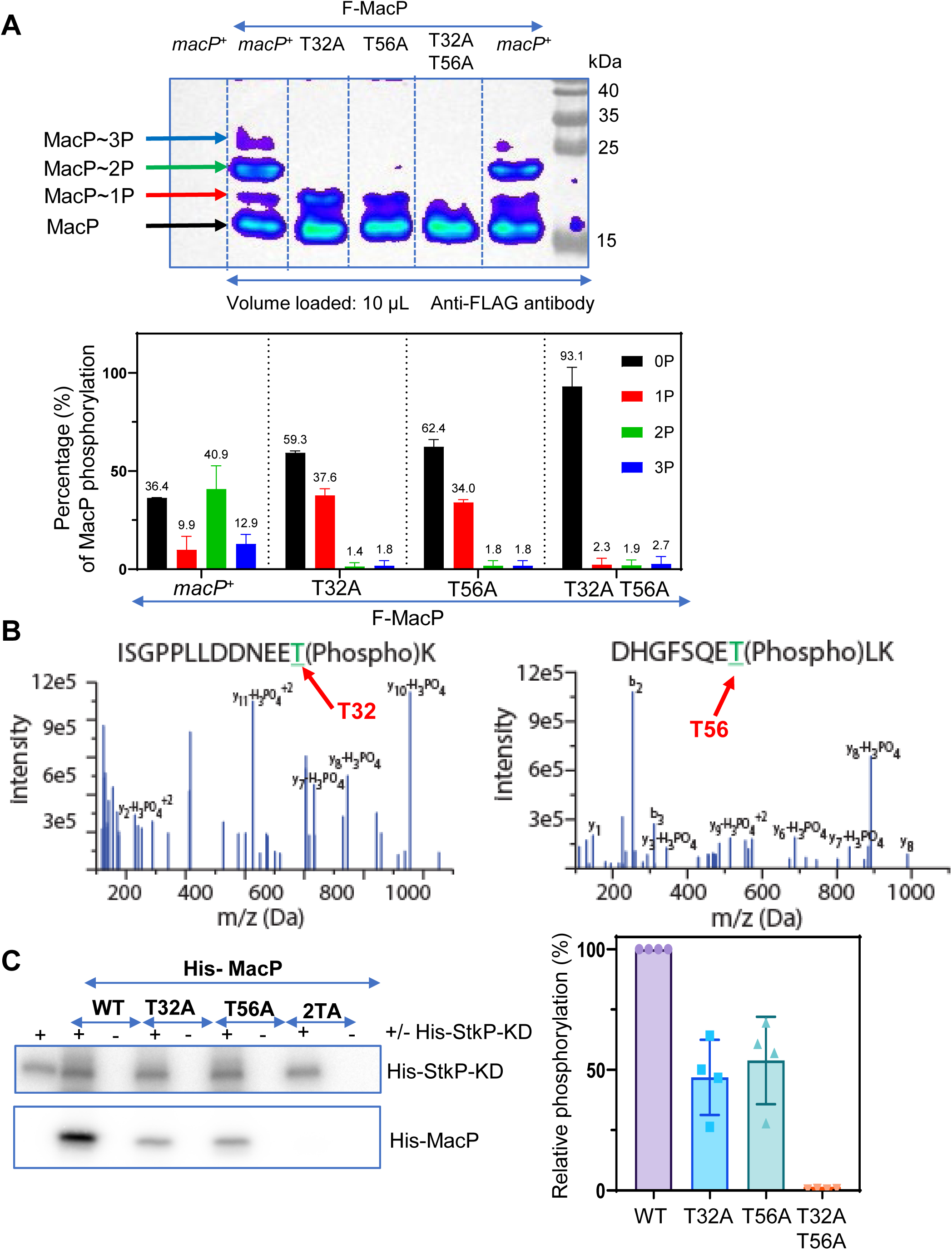
MacP(*Spn*) is phosphorylated at Thr32 and Thr56. **(A)** Top panel, MacP phosphorylation sites in *S. pneumoniae* assessed by 75 µM Phos-tag SDS-PAGE as described in *Experimental procedures* for the following strains: WT F-MacP (lane 2 (IU17032) and lane 6 (IU17033; second isolate) and MacP phosphoablative mutants (lane 3, F-MacP(T32A) (IU16978); lane 4, F-MacP(T56A) (IU16980); and lane 5, F-MacP (T32A T56A) (IU17035)). A representative western blot is shown of a Phos-tag gel of F-MacP and mutant MacP variants enriched by pull downs of lysates of cells grown exponentially in BHI broth. Positions of unphosphorylated MacP, F-MacP∼P, F-MacP∼P2, and F-MacP∼P3 bands are indicated in by colored arrows keyed to the graph in the bottom panel, which shows means ± SEM of the percentage unphosphorylated (0) and singly, doubly, and triply phosphorylated F-MacP from 2 biological replicates. **(B)** Fragmentation spectra from tandem MS of the peptides ISGPPLLDDNEET(phospho)K, ISGPPLLDDNEET(phospho)KILPTSSSR and DHGFSQET(phospho)LK from MacP isolated by co-IP with IreB-L-F3 bait (**Fig. S6**) as described in *Experimental procedures*. The precursor peptide had a charge of +2, which is consistent with a singly phosphorylated peptide. Fragmentation of this precursor peptide yielded the spectra shown. **(C)** Left panel, representative radiograph of the *in vitro* kinase assay with purified His-MacP variants and the StkP kinase domain (His-StkP-KD) described in *Experimental procedures*. Proteins were separated by 16% SDS-PAGE. Lane 1, StkP-KD; lane 2, MacP-WT + StkP-KD; lane 3, MacP-WT; lane 4, MacP-T32A + StkP-KD; lane 5, MacP-T32A; lane 6, MacP T56A + StkP-K D; lane 7, MacP-T56A; lane 8, MacP-2TA + StkP-KD; and lane 9, MacP-T32A T56A(2TA). Right panel, quantitation of the kinase assays of MacP phosphorylation from at least four independent experiments.

An MS analysis further confirmed primary phosphorylation of Thr32 and Thr56 in non-tagged MacP in strain D39 (**Fig. 2B**), which is the progenitor strain of Rx1 used before (Ulrych *et al*., 2021). IreB^+^-L-F3, which is a negative regulator of the MurZ and MurA enzymes (Tsui *et al*., 2023), was used as bait in a co-IP experiment (**Fig. S6**). IreB-L-F^3^ pulled down MacP^+^, suggesting a possible complex containing IreB and MacP. MS/MS fragmentation spectra of MacP^+^ peptides indicated phosphorylation at both Thr32 and Thr56 (**Fig. 2B**). Finally, approximately phosphorylation of Thr32 and Thr56 in MacP was detected in biochemical reactions containing purified His-tagged His-MacP^+^ or His-MacP phosphoablative variants and His-tagged StkP Ser/Thr kinase catalytic domain (His-StkP-KD) (**Fig. 2C**). About the same level of His-StkP-KD autophosphorylation was seen in reactions containing WT His-MacP and three phosphoablative His-MacP variants. In contrast, phosphorylation of His-MacP(T32A) or His-MacP(T56A) was half that of WT His-MacP^+^, and no phosphorylation of His-MacP(T32A T56A) (2TA) was detected (**Fig. 2C**). Altogether, we conclude that about 40% of MacP was doubly phosphorylated at Thr32 and Thr56 in non-stressed, exponentially growing *Spn* cells (**Fig. 2A**). Yet, phosphorylation of Thr32 and Thr56 in MacP did not play a detectable role in positive activation of aPBP2a in transformation assays or during growth of *S. pneumoniae* in culture (**Fig. 1B, S3A,** and **S3D**).

### 2.2. The cytoplasmic domain of MacP from amino acids 1-58 is not required for activation of aPBP2a activity

The topology of MacP consists of an N-terminal cytoplasmic domain, a transmembrane (TM) domain, and 4-amino-acid C-terminal extracellular domain (**Fig. 1A** and **S1B**-**S1D**). The MacP(TM) domain was shown previously to be important for midcell localization (Fenton *et al*., 2018). Since phosphorylation was not necessary, we determined which MacP domains and interactions are required for aPBP2a activation. To investigate the functions of the N- and C-terminal domains, truncated MacP derivatives were generated and tested for function in transformation assays with Δ*pbp1a*. Consurf conservation analysis (Yariv *et al*., 2023) guided the choice of amino acids to truncate in the N-terminal and transmembrane domains of MacP (**Fig. S1E**).

AlphaFold3 predicted that the N-terminal domain of MacP is largely unstructured to Arg69, with the exception of a short alpha-helical domain from Asp8 to Asn15 (**Fig. 1A** and **3A**). This lack of structure resulted in a non-significant overall structure prediction for MacP (pTM = 0.29). Arg69 begins a predicted alpha-helical domain, in which Val78 is the first predicted amino acid of the MacP(TM) domain (see below). Based on amino acid conservation, we designed and constructed MacP cytoplasmic-domain deletions and TM-truncation mutants: *macP*(Δ5-8), *macP*(Δ21-58), *macP*(Δ30-33), and *macP*(Δ46-53) (**Fig. 3A** and **S4A**). Growth curves and microscopy showed that these *macP* truncation mutants had no growth or morphological phenotypes compared to WT (**Fig. S5B**; data not shown). All cytoplasmic-domain truncation mutants were transformable with the Δ*pbp1a* amplicon, resulting in >500 normal-sized colonies similar to the WT strain (lines 1, 2-7, **Fig. 3B**). Growth curves and microscopy showed that Δ*pbp1a macP*(Δ5-8), Δ*pbp1a macP*(Δ21-58), Δ*pbp1a macP*(Δ30-33), and Δ*pbp1a macP*(Δ46-53) mutants grew slower and formed narrow, smaller cells than WT, similar to the Δ*pbp1a* single mutant (**Fig. S5C** and **S5D**)(Land & Winkler, 2011). Notably, the *macP*(Δ21-58) deletion lacked both the Thr32 and Thr56 phosphorylation sites, confirming that MacP(Thr32) and MacP(Thr56) phosphorylation and the surrounding amino acids in the cytoplasmic domain were not required for activation of aPBP2a activity (line 5, **Fig. 3B**). We determined whether MacP cytoplasmic-domain truncations were stably expressed by western blotting F-tagged MacP variants (**Fig. 3C** and **3D**). All F-tagged cytoplasmic deletion mutants, similar to the untagged mutants, were not synthetically lethal with Δ*pbp1a*. While most variants were expressed at over 70% of the WT level, MacP(Δ21-58) was expressed at only ≈11% of the WT amount; nevertheless, *macP*(Δ21-58) was not synthetically lethal with Δ*pbp1a* (line 5, **Fig. 3B** and **S5C**). This result suggested that cytoplasmic amino acids (21–58) impact MacP stability and that MacP is present in excess, since ≈11% MacP(Δ21-58) expression was sufficient for aPBP2a activation in transformation assays. Co-IP experiments further showed that MacP cytoplasmic truncation mutants still interacted with aPBP2a or GpsB, indicating these cytoplasmic regions did not play major roles in the interaction with these two proteins (**Fig. S4A** and **S5A**).

**Fig. 3.**
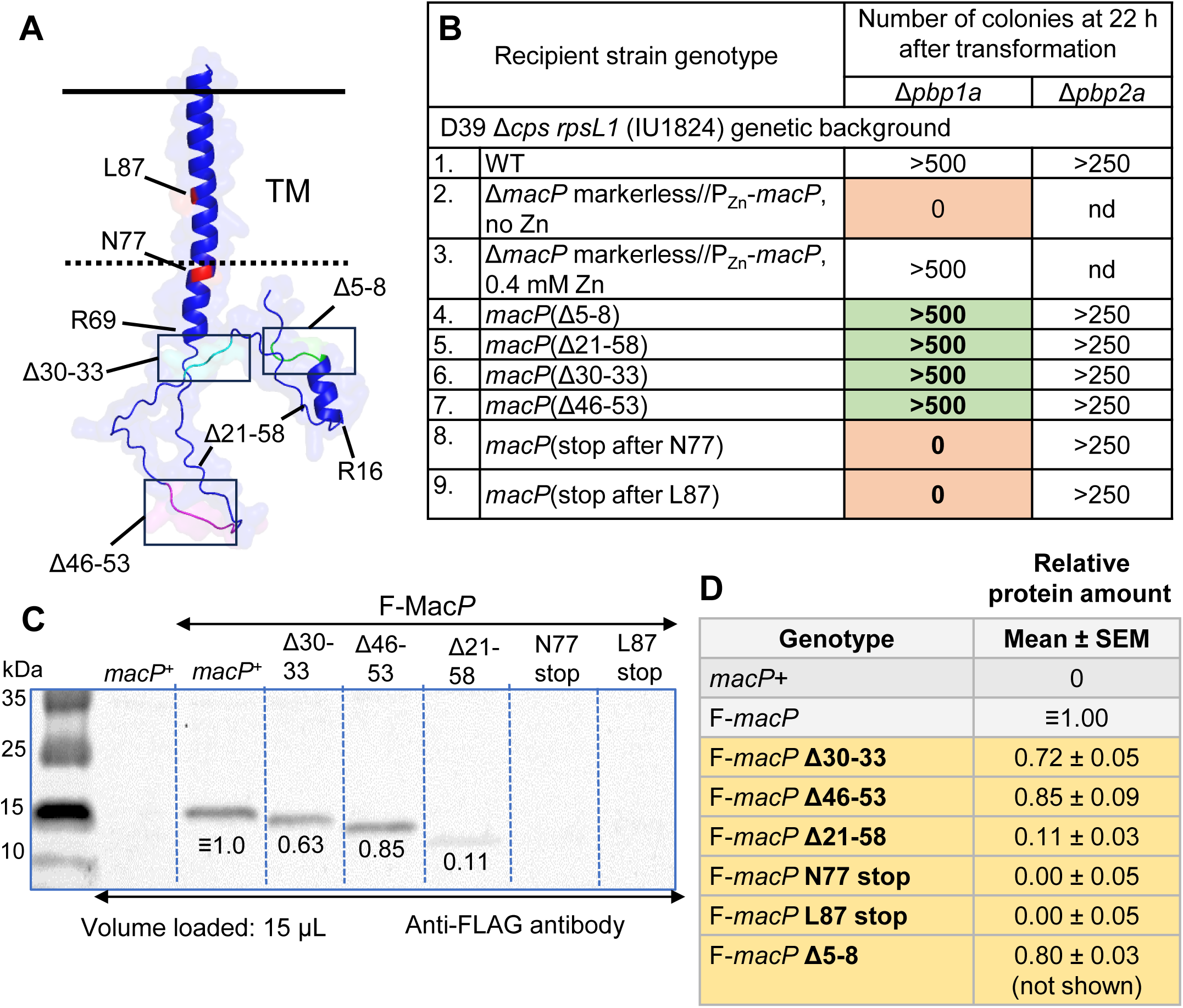
MacP cytoplasmic deletion mutants are not synthetically lethal with Δ*pbp1a*. **(A)** AlphaFold3-predicted structure of MacP(*Spn*). The Δ5-8, Δ21-58, Δ30-33, and Δ46- 53 deleted regions are boxed, and positions of stop codons in the TM domain are indicated by red. **(B)** Transformation assay with the Δ*pbp1a*::P_c_-*erm* test or Δ*pbp2a*::P_c_-*erm* control amplicon was performed as described in the *Experimental procedures* and the legend to **Fig.1B** for the following strains: WT (IU1824); Δ*macP* markerless//P_Zn_-*macP* (IU15084) ± Zn; *macP*(Δ5-8) (IU17228); *macP*(Δ21-58) (IU17386); *macP*(Δ30-33) (IU17384); *macP*(Δ46-53) (IU17232); *macP*(stop after N77) (IU17236); and *macP*(stop after L87) (IU17240). All colonies showed WT morphology on TSAII-BA plates containing erythromycin. Similar results were obtained for two or more biological replicates. **(C)** Representative western blot showing relative cellular amounts of F-*macP* and F-*macP* mutant proteins in the strains indicated below. 15 µL of protein lysate was loaded into each lane, and protein amount in each lane was normalized by staining. The predicted molecular mass of F-MacP was 13 kDa. Numbers below bands indicate relative F-MacP amounts in this blot. **(D)** Summary table (means ± SEM) of relative protein amounts from quantitative western blots for three biological replicates performed as described in *Experimental procedures* and illustrated in **Fig. S3E**. Strains used: *macP*^+^ (IU1824); F-*macP* (IU17032); F-*macP* Δ5-8 (IU19530); F-*macP* Δ21-58 (IU18348); F-*macP* Δ30-33 (IU18346); F-*macP* Δ46-53 (IU18345); F-*macP* N77 stop (IU18351); F-*macP* L87 stop (IU18352).The blot is not shown for the F-*macP*(Δ5-8) (IU19530) mutant listed in the table.

### 2.3 Identification of amino acids in MacP(TM) and the juxtamembrane region of aPBP2a(TM) important for MacP activation of aPBP2a

We examined how changes in the MacP(TM) domain affected its function. We constructed mutations that removed MacP(TM) by introducing a stop codon after N77 or L87 (**Fig. S1B-D**). Both MacP(TM) stop mutations were synthetically lethal with Δ*pbp1a* (lines 8 and 9, **Fig. 3B** and **S4A**). F-MacP(N77(stop)) or F-MacP(L87(stop)) cytoplasmic protein fragments were not detected in western blots (**Fig. 3C**, **3D**, and **S4**), indicating that the absence of the TM domain destabilized MacP, resulting in synthetic lethality with Δ*pbp1a*. In contrast, MacP variants truncated by stop codons were stabilized in fusions used for B2H assays (upper panel, **Fig. S7A**). Nevertheless, the truncated MacP variants lacking the TM domain did not bind to aPBP2a or GpsB in B2H assays (lower panel, **Fig. S7A**), consistent with the requirement of MacP(TM) for interactions leading to midcell localization in pneumococcal cells (Fenton *et al*., 2018). The self-interaction of MacP in B2H assays, which is a control for functional intactness for interaction, was also abolished by the absence of MacP(TM) (lower panel, **Fig. S7A**). Whether MacP dimerizes in *S. pneumoniae* cells was not addressed in this study.

We next identified amino acids in MacP(TM) and in the predicted juxtamembrane region of aPBP2a important for MacP activation of aPBP2a. We used AlphaFold3 (Abramson *et al*., 2024) to model a moderately accurate structure (pTM = 0.68) of a putative MacP-aPBP2a complex, where the predicted aPBP2a structure alone was highly accurate (pTM = 0.74) (**Fig. 4A**). We then used PyMOL to identify amino acids in predicted interfaces between MacP(TM) (V78-A100) and aPBP2a (**Fig. S7B**). Based on these predictions, we changed amino acids in potential interfaces of the MacP(TM) domain (variants *macP*(R76A N80A) (**Fig. 4B**) and *macP*(L83A I86A L87A) (**Fig. 4C**)), in the C-terminal region of MacP (variant *macP*(M101A L103A) (**Fig. 4D**)), and in the juxtamembrane region of PBP2a (variant *pbp2a*(R51A H53A K56A) (**Fig. 4B**)). The boundaries of the aPBP2a(TM) domain have not been determined experimentally, and other predictions suggest that the juxtamembrane region is part of the aPBP2a(TM) domain (Guilmi *et al*., 1999). We assessed aPBP2a function in these variants by transformation assays with a Δ*pbp1a* amplicon (**Fig. 5A** and **S4A**). The *macP*(L83A I86A L87A) or *pbp2a*(R51A H53A K56A) mutant could not be transformed with Δ*pbp1a* (lines 3 or 5, **Fig. 5A, S4A,** and **S4B**), suggesting that these amino acids are important for function. In contrast, *macP*(R76A N80A) could be transformed and was functional (line 2, **Fig. 5A**). The F-MacP(L83A I86A L87A) or F-PBP2a(R51A H53A K56A) mutant variant was expressed at 46% or 140% of the WT level, respectively, in quantitative western blots (**Fig. 5B**-**5D**).

**Fig. 4.**
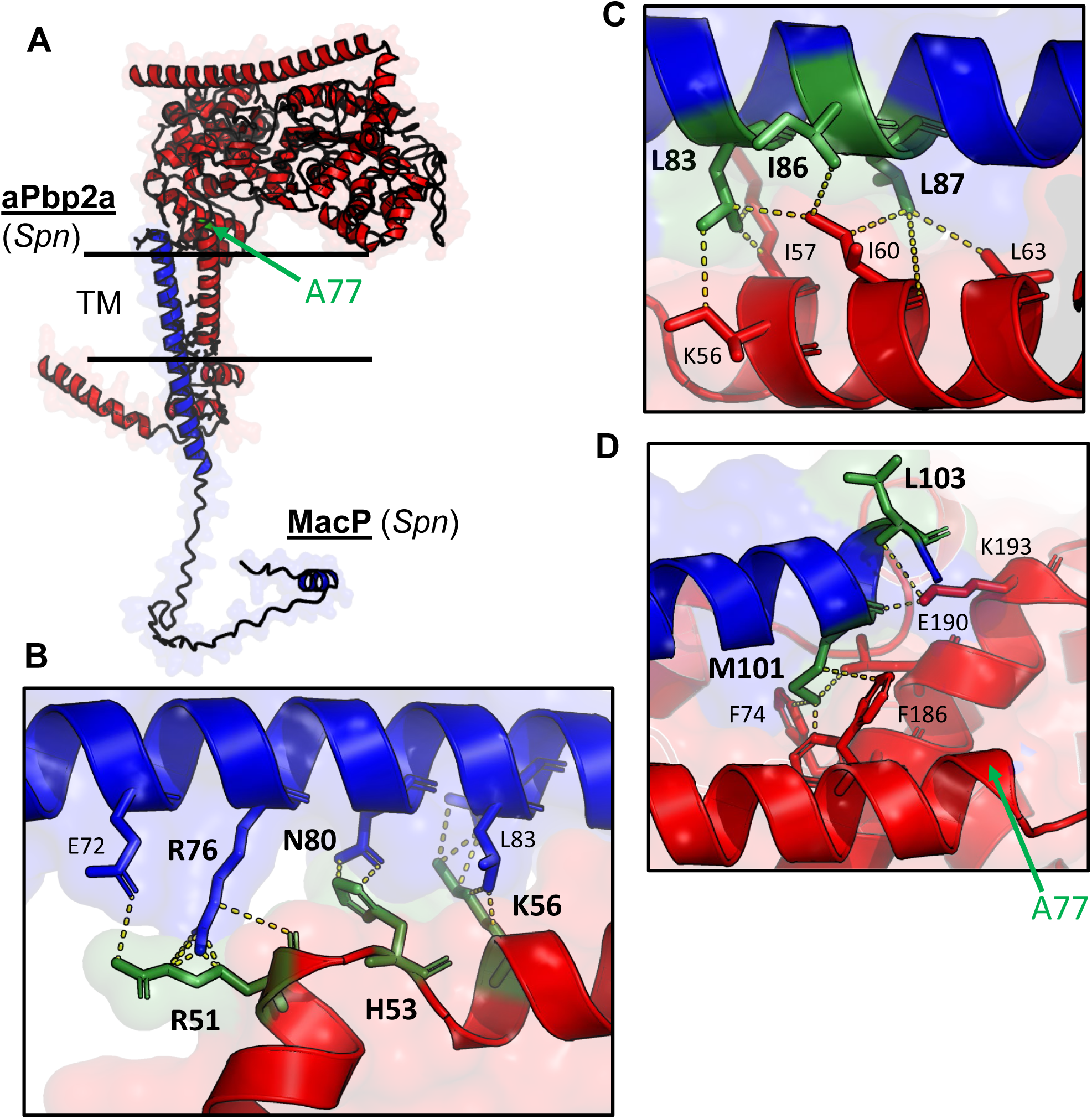
Putative interactions of MacP(TM) with the juxtamembrane region and TM domain of PBP2a. **(A)** Structure of the MacP:aPBP2a complex was predicted by AlphaFold 3 and presented as described in *Experimental procedures*. MacP (1-104 amino acids) and aPBP2a (1-731 amino acids) are shown in blue and red, respectively. aPBP2a(A77) implicated in activation (Midonet *et al*., 2023) is indicated in green. The pTM for aPBP2a was 0.74, and the pTM and ipTM for the MacP:aPBP2a complex were 0.44 and 0.68, respectively, where the cytoplasmic domain of MacP was predicted to be largely unstructured. The TM domain of MacP (V78-A100) or aPBP2a (I57-V76) were predicted by TMHMM (Sonnhammer *et al*., 1998, Krogh *et al*., 2001), A different prediction suggested that aPBP2a(TM) starts at Y52 (**Fig. S1D)**, which is denoted as the juxtamembrane region here (Guilmi *et al*., 1999). **(B-D)** Enlargement of transmembrane regions of MacP (blue) and aPBP2a (red) predicted to interact, where interacting amino acids drawn as sticks. Dashed yellow lines indicate polar interactions between amino acids. **(B)** Putative aPBP2a(R51 H53 K56) (juxtamembrane region) interaction with MacP(E72 R76 N80 L83) (TM domain). **(C)** Putative MacP(L83 I86 L87) (TM domain) interaction with aPBP2a(K56 I57 I60 L63) (TM domain). **(D)** MacP(M101 L103) (extracellular domain) interaction with aPBP2a(F74 F186 E190 K193) (extracellular domain). Amino acids in green were changed to Ala to test for phenotypes consistent with interactions (see **Fig. 5**). See text for additional details.

**Fig. 5.**
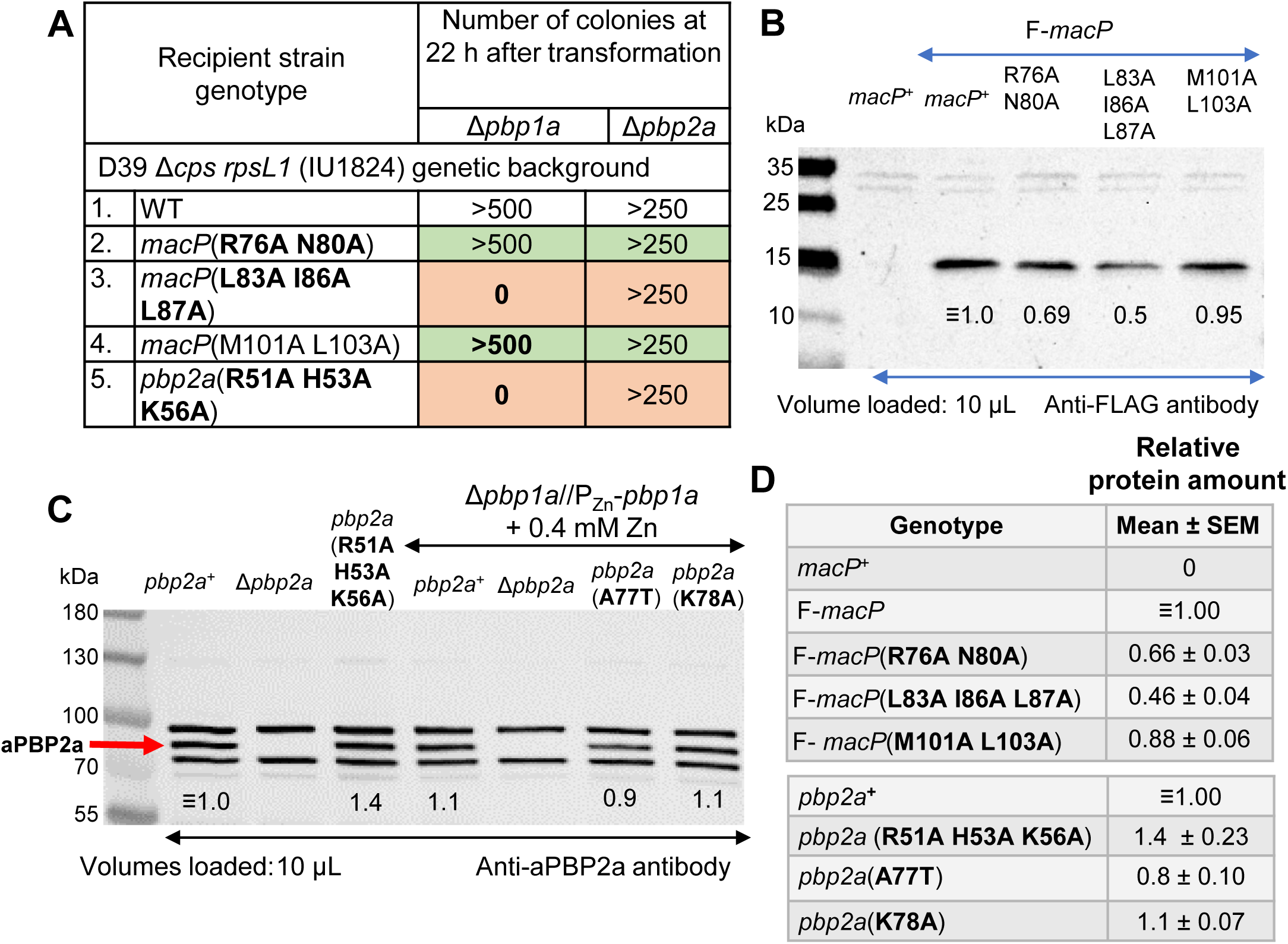
*macP*(L83A I86A L87A) and *pbp2a*(R51A H53A K56A) are synthetically lethal with Δ*pbp1a*, but the proteins are still expressed. **(A)** Transformation assay with the Δ*pbp1a*::P_c_-*erm* test or Δ*pbp2a*::P_c_-*erm* control amplicon was performed as described in the *Experimental procedures* and the legend to **Fig.1B** for the following strains: WT (IU1824); *macP*(R76A N80A) (IU19522); *macP*(L83A I86A L87A) (IU19526); *macP*(M101A L103A) (IU19529); *pbp2a*(R51A H53A K56A) (IU19533). All colonies showed WT morphology on TSAII-BA plates containing erythromycin. Similar results were obtained for two or more biological replicates. **(B)** Representative western blot showing relative cellular amounts of F-MacP and F-MacP mutant proteins in the strains indicated below. 10 µL of protein lysate was loaded into each lane, and protein amount in each lane was normalized by staining. The predicted molecular mass of F-MacP was 13 kDa. Numbers below bands indicate relative F-MacP amounts in this blot. **(C)** Representative western blot showing relative cellular amount of WT aPBP2a and aPBP2a mutant variants in the strains indicated below. Western blotting was performed and quantitated as described in *Experimental procedures* using anti-aPBP2a primary antibody and Licor IR Dye800 CW secondary antibody. 10 µL of protein lysate was loaded into each lane, and protein amount in each lane was normalized by staining. The position of aPBP2a is indicated by the red arrow. Numbers below bands indicate relative aPBP2a amounts in this blot. **(D)** Summary table (means ± SEM) of relative protein amounts from quantitative western blots for three biological replicates of the F-MacP and variants (top panel) and two biological replicates of the aPBP2a and variants (left panel). Quantitation was performed as described in *Experimental procedures* and illustrated in **Fig. S3E**. F-MacP strains used in **(B)** and **(D)**: m*acP*^+^(IU1824); *F-macP* (IU17032); F-*macP*(R76A N80A) (IU19524); F-*macP*(L83A I86A L87A) (IU19528); and F-*macP*(M101A L103A) (IU19529). aPBP2a stains used in **(C)** and (D): *pbp2a*^+^(IU1824); Δ*pbp2a* (IU13256); *pbp2a*(R51A H53A K56A) (IU19533); *pbp2a*^+^ Δ*pbp1a*//P_Zn_-*pbp1a* (IU14357); Δ*pbp2a* Δ*pbp1a* //P_Zn_-*pbp1a* (IU19216); *pbp2a* (A77T) Δ*pbp1a* //P_Zn_-*pbp1a* (IU19240); and *pbp2a* (K78A) Δ*pbp1a* //P_Zn_-*pbp1a* (IU19242).

Qualitative B2H assays in *E. coli* indicated some level of interaction between MacP(L83A I86A L87A) or PBP2a(R51A H53A K56A) and aPBP2a or MacP, respectively (**Fig. S7C**). However, quantitative co-IP assays demonstrated a large (≈95%) drop in the interaction between F-MacP(L83A I86A L87A) and aPBP2a in *Spn* cells (**Fig. S7D** and **S7E**), which would account for the lack of MacP(L83A I86A L87A) activation of aPBP2a seen in transformation assays (line 3, **Fig. 5A** and **Fig. S4A**). There was also an ≈80% drop in the interaction between F-Mac-(R76A N80A) and aPBP2a (**Fig. S7D** and **S7E**), but this drop was not sufficient to prevent aPBP2a activation in transformation assays (line 3, **Fig. 5A**). Finally, F-MacP(M101A L103A) showed an ≈30% reduction in its interaction with aPBP2a (**Fig. S7D** and **S7E**), but was functional in stimulating aPBP2a activity (line 4, **Fig. 5A**). Notably, GFP-MacP(L83A I86A L87A) localized normally to midcell, like WT GFP-MacP (rows 2 and 5, **Fig. S7F**). Together, these results demonstrated that certain amino acids in the MacP(TM) interface and aPBP2a juxtamembrane region interface, such as MacP(L83 I86 L87) and aPBP2a(R51 H53 K56), are required for MacP activation of aPBP2a. The changes in MacP(L83A I86A L87A) abolished a required interaction with aPBP2a.

### 2.4. aPBP2a(A77T) bypasses the requirement for MacP activation, but does not necessarily cause severe defects in cell morphology

Midonet and coworkers recently reported that aPBP2a(A77T) bypasses the requirement of MacP function (Midonet *et al*., 2023). aPBP2a(A77) is located at the end of the TM helix at the start of a flexible domain leading to the glycosyltransferase (GT) domain (**Fig. 4A**, **4D**, and **S1B-S1D**). A current model is that MacP binding alters this interface domain to activate aPBP2a (Midonet *et al*., 2023). We confirmed that *pbp2a*(A77T) bypasses the requirement for MacP when aPBP1a is depleted in a Δ*macP* mutant, but not to the WT level under the conditions tested (orange triangles, **Fig. S8A**). In contrast, a change in the adjacent amino acid (K78A) did not bypass the requirement for MacP (**Fig. S8A**). aPBP2a(A77T) and aPBP2a(K78A) were expressed at WT level in cells growing in BHI broth (bottom, **Fig. 5D**), and both mutants grew similarly to WT and a Δ*pbp2a* mutant in BHI or THY broth (**Fig. S8B** and **S8D**). As reported previously (Midonet *et al*., 2023), the *pbp2a*(A77T) mutant had a small (≈1.1X) increase in cell width compared to WT (**Fig. S8B** and **S8C**). However, we did not observe heterogenous, defective cell morphologies of *pbp2a*(A77T) mutant cells, which appeared similar in shape and size to WT cells in BHI or THY broth (**Fig. S8C** and data not shown). Therefore, our results did not support the conclusion that the moderate constitutive activation of aPBP2a(A77T) significantly altered pneumococcal cell morphology under these growth conditions.

### 2.5. MacP interacts with several proteins involved in septal PG (sPG) and elongasome PG (ePG) synthesis

A previous study concluded that MacP interacts with aPBP1a and aPBP2a, but not with itself, MreC, MreD, or CozE in B2H assays (Fenton *et al*., 2018). In addition, MacP was shown to form complexes with GpsB by co-IP and B2H assays (Stauberová *et al*., 2024). To further confirm these interactions and determine if other PG synthesis proteins interact with MacP, we performed additional bidirectional B2H assays wih both vector combinations (**Fig. 6A**). The B2H assays confirmed interactions of MacP with aPBP1a, aPBP2a, and GpsB and detected bidirectional interactions with several other proteins involved in PG synthesis or cell division, including the MpgA muramidase, components of the ePG elongasome (RodZ, MreC, PBP2b, and RodA), and components of the sPG divisome (EzrA, FtsL, DivIB(FtsQ)) (**Fig. 6A**). MacP self-interaction was also detected. Unidirectional and/or weaker interactions were observed between MacP and bPBP2x or FtsW (sPG synthase), MreD (ePG elongasome), FtsA (FtsZ anchor/regulator), or DivIVA (septal regulator). By comparison, B2H assays indicated interactions of GpsB or MacP with many of the same proteins, with the exceptions of StkP (Ser/Thr kinase), aPBP2b, and RodA (red boxes, **Fig. 6A**). In addition, B2H assays did not detect an interaction between MacP and IreB (data not shown).

**Fig. 6.**
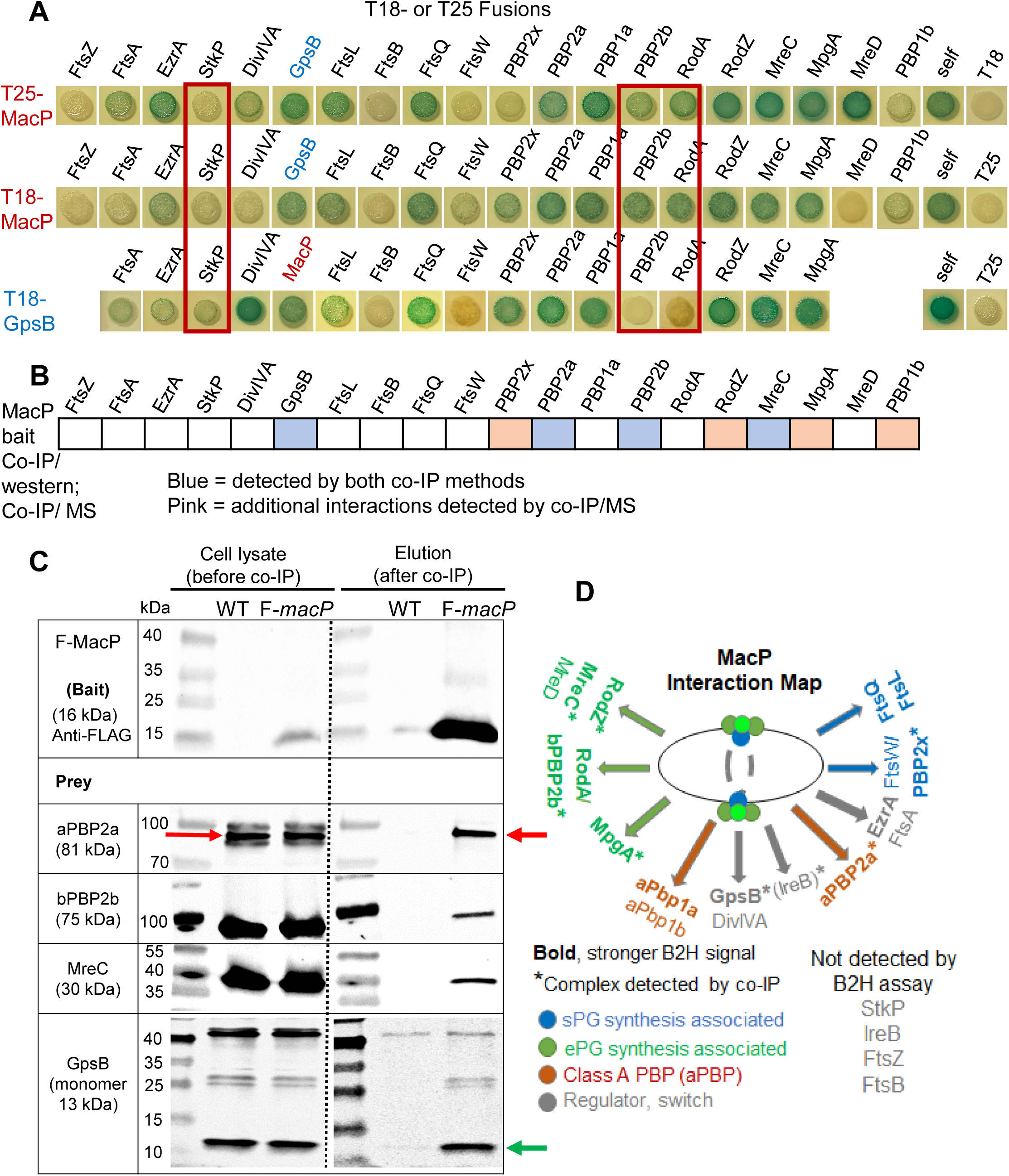
MacP interacts with several proteins involved in septal PG (sPG) and elongasome PG (ePG) synthesis. (A) Interactions of MacP detected by B2H assays. MacP interacted with EzrA, GpsB, FtsL, FtsQ, aPBP2a, aPBP1a, bPBP2b, RodA, RodZ, MreC, MpgA, or itself in both directions in in B2H assays performed as described in *Experimental procedures*. MacP also interacted with FtsA, DivIVA, FtsW, bPBP2x, MreD, or aPBP1b unidirectionally and/or with a lower signal. No interactions were detected with StkP and IreB (data not shown). T25 or T18 fusions were expressed from low- or high-copy plasmids, respectively. Plasmid pairs pKNT25/pUT18 and pKT25-*zip*/pUT18C-*zip* were used as negative (-ve) and positive (+ve) controls, respectively (data not shown). Agar plates were photographed after 40 h at 30°C. The results are representative of at least two biological replicates. B2H assay results for GpsB from (Rued *et al*., 2017) are shown for comparison. MacP showed a similar interaction pattern to that of GpsB, except for the interactions highlighted in red boxes, indicating that GpsB interacted with StkP, but not with bPBP2b or RodA. **(B)** Interactions of MacP determined by co-IP using F-macP as bait with MS detection (**Table S4**) and/or in the pairwise co-IP experiments below. These complexes containing MacP contained many proteins shown to interact with MacP in the B2H assays shown above. See text for additional details. (**C**) Representative pairwise co-IP with western-blot detection showing complexes containing F-MacP (bait) with aPBP2a (red arrow), bPBP2b, MreC, or GpsB (green arrow). Prey were detected with polyclonal antibodies to the indicated native proteins. Strains used: *macP^+^* WT (IU1824) for control and F-*macP* (IU17032) for bait. Similar results were obtained for two or more biological replicates. Note that of the three bands detected by anti-aPBP2a antibody (see **Fig. 5C**), only the central band corresponding to aPBP2a (red arrow) was detected after co-IP. **(D)** Summary of MacP interactions determined by B2H assays and co-IP experiments. Qualitative interaction strength in B2H assays, complex detection by co-IP, and protein functions are indicated. A complex containing MacP and IreB was detected by unbiased co-IP/MS with IreB-F^3^ (**Fig. S6**) or F-MacP (**Table S4**) as bait, but a direct interaction was not detected by the B2H assay (data not shown).

Co-IP assays showed that MacP was in complexes at some stage of pneumococcal cell cycle with many of the proteins interrogated in the B2H assay (**Fig. 6B**). The criterion for likely complex formation was detection of peptides in unbiased co-IP/MS from samples pulled down by the F-MacP bait, but absent or barely detectable in the untagged-MacP control (**Fig. 6B**; **Table S4**). By this criterion, MacP was in complexes with GpsB, bPBP2x, aPBP2a, aPBP2b, RodZ, MreC, or MpgA (**Fig. 6B**) and additionally with IreB (negative regulator of MurZ/MurA), PgdA (PG deacetylase), FtsX (PG hydrolase subunit), or aPBP1b (**Table S4**). Last, we used co-IP with western-blot detection of specific proteins to confirm complexes containing MacP and aPBP2a, GpsB, bPBP2b, or MreC at some stage of the cell cycle (**Fig. 6C**). Co-IP experiments also indicated complex formation between these four proteins and the phosphomimetic F-MacP(T32E T56E) mutant or phosphoablative F-MacP(T32A T56A) mutant as bait instead of WT F-MacP (data not shown). Together, these studies corroborated interactions between GpsB and aPBP2a (Cleverley *et al*., 2019), MacP and aPBP2a (Fenton *et al*., 2018), and MacP and GpsB (Stauberová *et al*., 2024). In addition, they showed that MacP interacts with numerous PG-synthesis and cell-division proteins in B2H assays and is in complexes with some of these proteins at stages of the cell cycle during exponential growth (**Fig. 6D**).

### 2.6. MacP and GpsB bind to different sites in aPBP2a, but the same amino acids in GpsB are required for interaction with aPBP2a or MacP

The above results show that GpsB and MacP, which are both required for activation of aPBP2a (Fenton *et al*., 2018, Rued *et al*., 2017), bind to each other and bind to aPBP2a. We next asked whether GpsB binding to aPBP2a or MacP was related. GpsB binds to a (S/R)RS(R/G)(K/S)xR motif at amino acids 31-37 in the N-terminal cytoplasmic domain of aPBP2a (**Fig. 7A** and **Fig. S1D**) (Cleverley *et al*., 2019). Deletion of segments of this binding motif in aPBP2a reduced binding to GpsB in B2H assays at earlier times of incubation, while minimally changing binding between aPBP2a and MacP (**Fig. 7A**). From this result, we conclude that MacP binds to aPBP2a in a region different from the GpsB-binding motif.

**Fig. 7.**
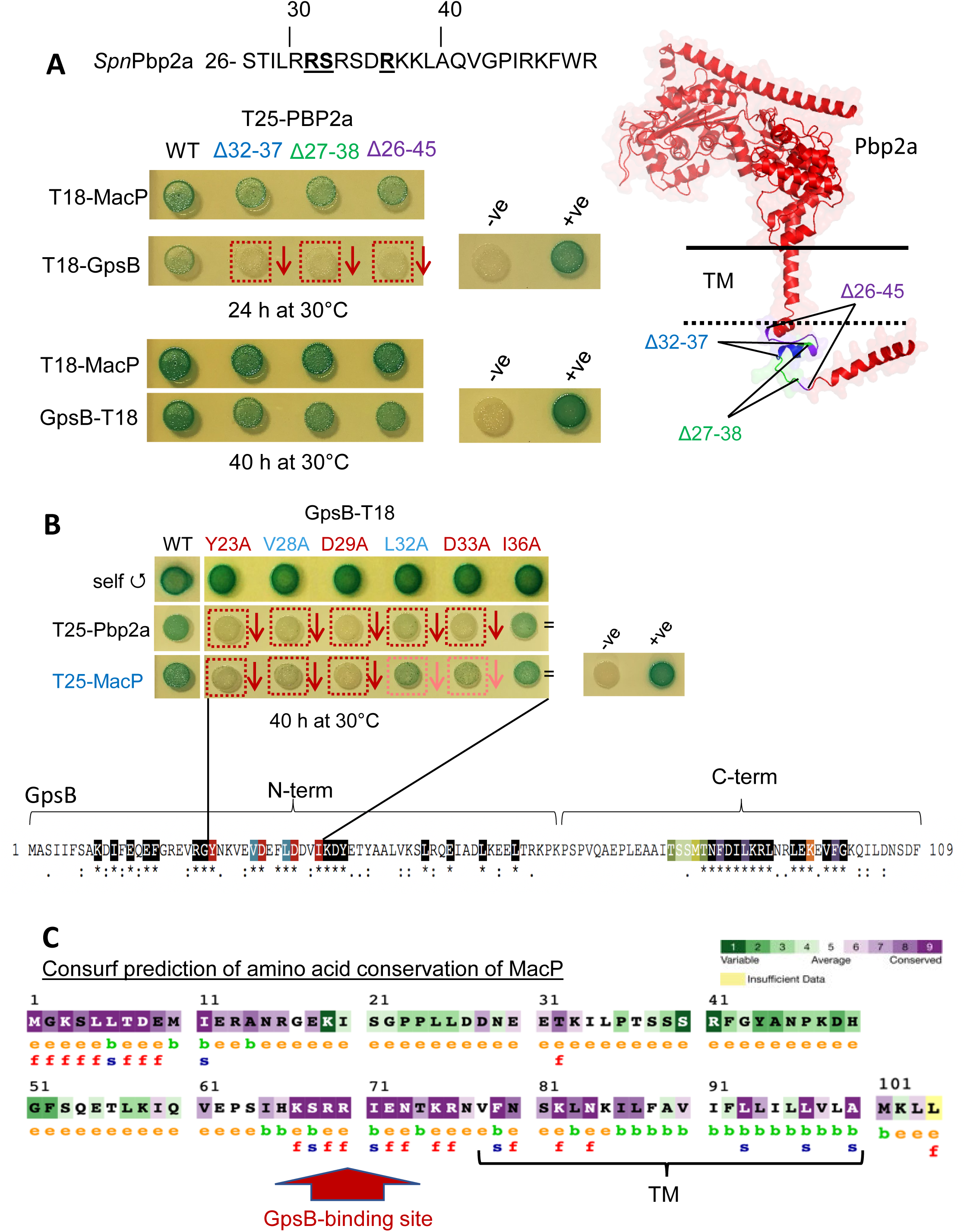
MacP and GpsB bind to different sites in aPBP2a, but the same amino acids in GpsB are required for interaction with PBP2a or MacP. **(A)** Top, sequence of GpsB-binding site in the amino terminus of PBP2a composed of conserved Arg31, Ser32, and Arg36 in an RSxxxR motif (Cleverley *et al*., 2019). Bottom, small deletions in the RSxxxR GpsB-binding motif of PBP2a eliminated GpsB binding, but did not change interaction with MacP. Interactions were assayed by B2H assays as described in *Experimental procedures.* Plasmid pairs pKNT25/pUT18 and pKT25-*zip*/pUT18C-*zip* were used as negative (-ve) and positive (+ve) controls, respectively. Agar plates were photographed after 36 h of incubation at 30°C. Similar results were obtained in two biological replicates. Right, predicted AlphaFold3 structure of aPBP2a (pTM = 0.74) showing the regions in the GpsB-binding site that were deleted: Δ32-37 (blue); Δ27-38 (green); and Δ26-45 (purple). **(B)** The same Ala substitutions in the conserved N-terminal domain of GpsB (indicated below) reduced interactions with aPBP2a (second row) or MacP (third row) in B2H assays. B2H assays were performed as described above, except that agar plates were photographed after 40h of incubation at 30 °C. Similar results were obtained for two biological replicates. **(C)** Conserved amino acids in *Spn* MacP determined by Consurf prediction as described in *Experimental procedures.* Deletion analysis (see **Fig. 3** and text) showed that amino acids before Lys58 can be deleted without abolishing MacP activation of aPBP2a activity. The conserved Arg-rich region in MacP adjacent to the TM domain resembles a GpsB-binding motif and was shown to mediate GpsB binding (see **Fig. 8** and **9**).

We next inquired whether the same region of GpsB was required for binding aPBP2a and MacP. Structural studies showed that amino acids 23-33 in the N-terminal domain of GpsB were required for binding the (S/R)RS(R/G)(K/S)xR motif of aPBP2a (**Fig. 7B**) (Cleverley *et al*., 2019). Because GpsB is a hexameric trimer of dimers, there are six of these motif-binding sites in each GpsB molecule, allowing binding of one GpsB molecule to different motif-containing proteins (Cleverley *et al*., 2019). Amino-acid changes in the GpsB-binding site decreased binding between GpsB and aPBP2a in endpoint B2H assays, as expected from previous studies (**Fig. 7B**) (Cleverley *et al*., 2019). Notably, the same amino-acid changes in GpsB reduced binding to MacP, except that changes in GpsB(L32) and GpsB(D33) reduced binding to aPBP2a more than to MacP. Together, these results show that the same region of GpsB binds to different motifs in aPBP2a and MacP.

### 2.7. MacP contains a variant GpsB-binding motif required for activation of aPBP2a activity

Cleverley et al. showed that GpsB interacts with aPBP2a through two Arg residues spaced 2-4 residues apart in the GpsB-binding motif (<u>RR</u>S<u>R</u>SD<u>R</u>K) (**Fig. 7A**) (Cleverley *et al*., 2019). Moreover, binding of aPBP2a molecules to two GpsB monomers in a dimer used different pairs of Arg residues (Arg31/Arg36 or Arg33/Arg36) (Cleverley *et al*., 2019). Therefore, we inspected the amino acid sequence of MacP for similarly spaced Arg residues that are conserved in MacP homologs. We identified a single putative cytoplasmic GpsB-binding motif (68-KS<u>RR</u>IENTK<u>R</u>-76) in MacP immediately adjacent to the TM domain (**Fig. 7C** and **S1D**).

To test whether MacP(68-KS<u>RR</u>IENTK<u>R</u>-76) plays a role in aPBP2a activation, we constructed a *macP*(S68A R69A R70A K75A R76A) pentuple mutant in the bacterial chromosome (**Fig. 8A**). The pentuple mutant grew at a rate and to a yield similar to those of WT (**Fig. 8B**). F-MacP(S68A R69A R70A K75A R76A) was expressed at about 34% of WT F-MacP (**Fig. S9A** and **S9B**). This level of expression was greater than the low (≈11%) relative expression of F-MacP(Δ21-58) sufficient to activate aPBP2a and prevent synthetic lethality with Δ*pbp1a* (**Fig. 3C**, **3D**, and **S5C**). Yet, transformation of *macP*(S68A R69A R70A K75A R76A) with Δ*pbp1a* resulted in severely impaired growth (**Fig. 8B** and **S9C**) and defective cell morphologies (**Fig. 8C**). AlphaFold3 modeling further predicted that MacP(68-KS<u>RR</u>IENTK<u>R</u>-76) interacts with the N-terminal domain of GpsB that binds to aPBP2a and other proteins (**Fig. 9D**). Together, these results suggest that the conserved MacP(68-KS<u>RR</u>IENTK<u>R</u>-76) region is important for MacP stability and binding to GpsB during activation of aPBP2a.

**Fig. 8.**
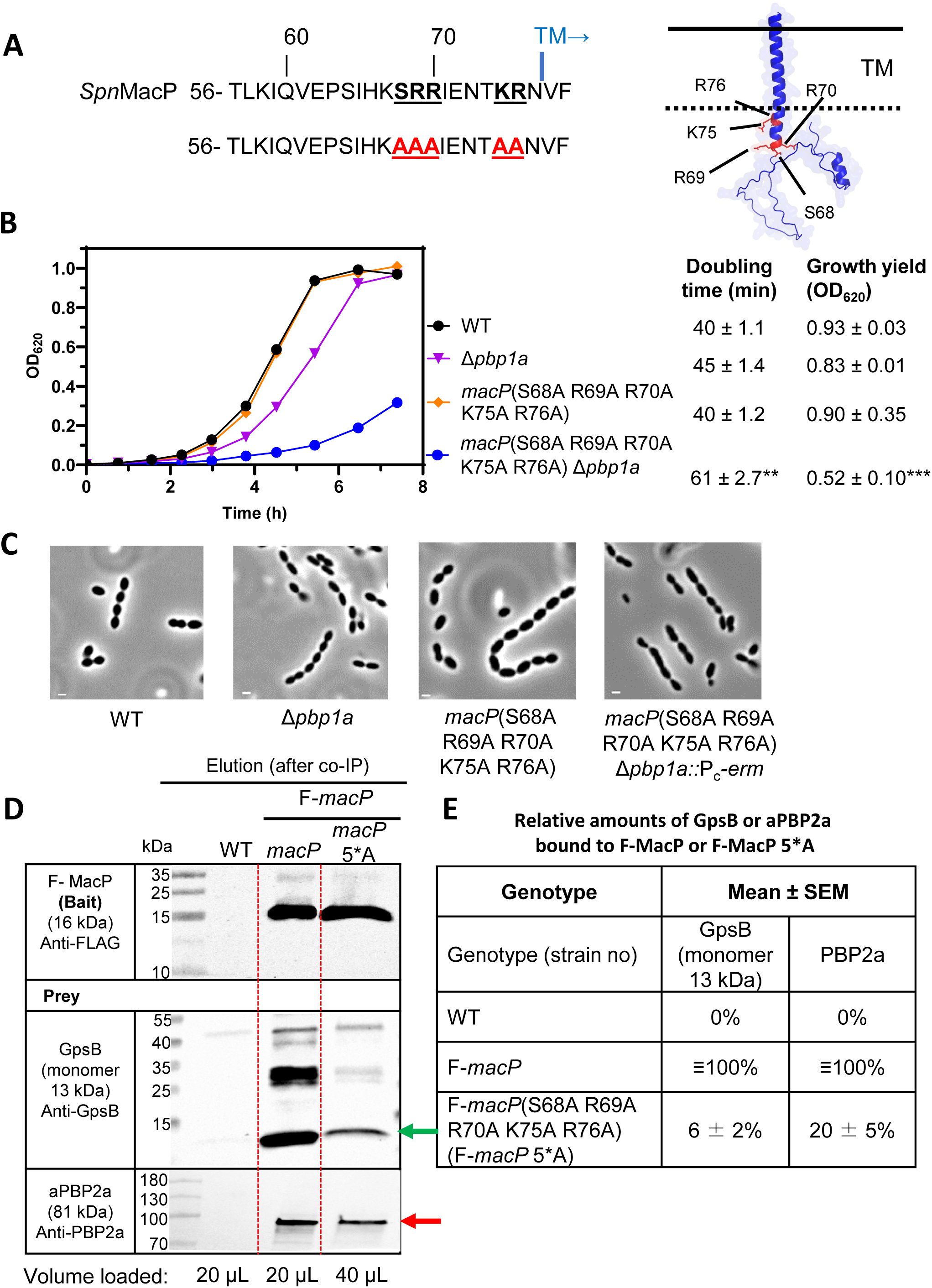
The conserved <u>SRR</u>IENT<u>KR</u> motif in MacP is important for stability, GpsB binding, and aPBP2a activation. **(A)** Left, MacP sequence containing the conserved, Arg-rich **<u>SRR</u>**xxxx**KR** motif (**Fig. 7C**), similar to that of the GpsB-binding motif in aPBP2a (**Fig. 7A)** (Cleverley *et al*., 2019). Right, AlphaFold3-predicted structure of MacP (pTM = 0.29), indicating the amino acids (in red) changed in the MacP(S68A R69A R70A K75A R76A) pentuple mutant. **(B)** Representative growth curves in BHI broth of the WT strain (IU1824), the Δ*pbp1a* mutant (IU6741); the *macP*(S68A R69A R70A K75A R76A; abbreviated: 5*A) pentuple mutant (IU21139), and the double *macP*(S68A R69A R70A K75A R76A) Δ*pbp1a* mutant (IU21169), which was impaired for growth compared to the other strains (see text and **Fig. S9C)**. Means ± SEM of doubling times and growth yields are shown for two biological replicates. Statistical comparisons to WT were done by one-way ANOVA with Dunnett’s multiple-comparisons test, where \*\**p* < 0.01 and \*\*\**p* < 0.001. **(C)** Representative phase-contrast micrographs of strains from **Fig 8B**. The *macP*(5*A) pentuple mutant and the *macP*(5*A) pentuple Δ*pbp1a* mutant exhibited aberrant cell morphologies compared to the other strains. Growth curves and phase microscopy were performed as described in *Experimental procedures*. Scale bar (left corners) = 1.0 µm. **(D)** Representative western blot of pairwise co-IP showing a substantial decrease in binding of GpsB or aPBP2a to F-MacP(S68A R69A R70A K75A R76A; 5*A) compared to WT F-MacP. Quantitative western blotting with anti-FLAG antibody showed that the amount of F-MacP(S68A R69A R70A K75A R76A) was ≈34% of the F-MacP amount (**Fig. S9A** and **S9B**); therefore, twice as much protein was loaded in the F-*macP*(5*A) lane. Co-IP with western blotting using polyclonal antibodies to GpsB (green arrow) or aPBP2a (red arrow) was performed and quantitated as described in *Experimental procedures* on strains: WT (IU1824); F-*macP* (IU17032); and F-*macP*(S68A R69A R70A K75A R76A) (IU21376). See **Fig. S9D** for input controls. **(E**) Summary table (means ± SEM) of relative amounts of GpsB or aPBP2a bound to F-*macP* or F-*macP*(5*A) for three biological replicates.

**Fig. 9.**
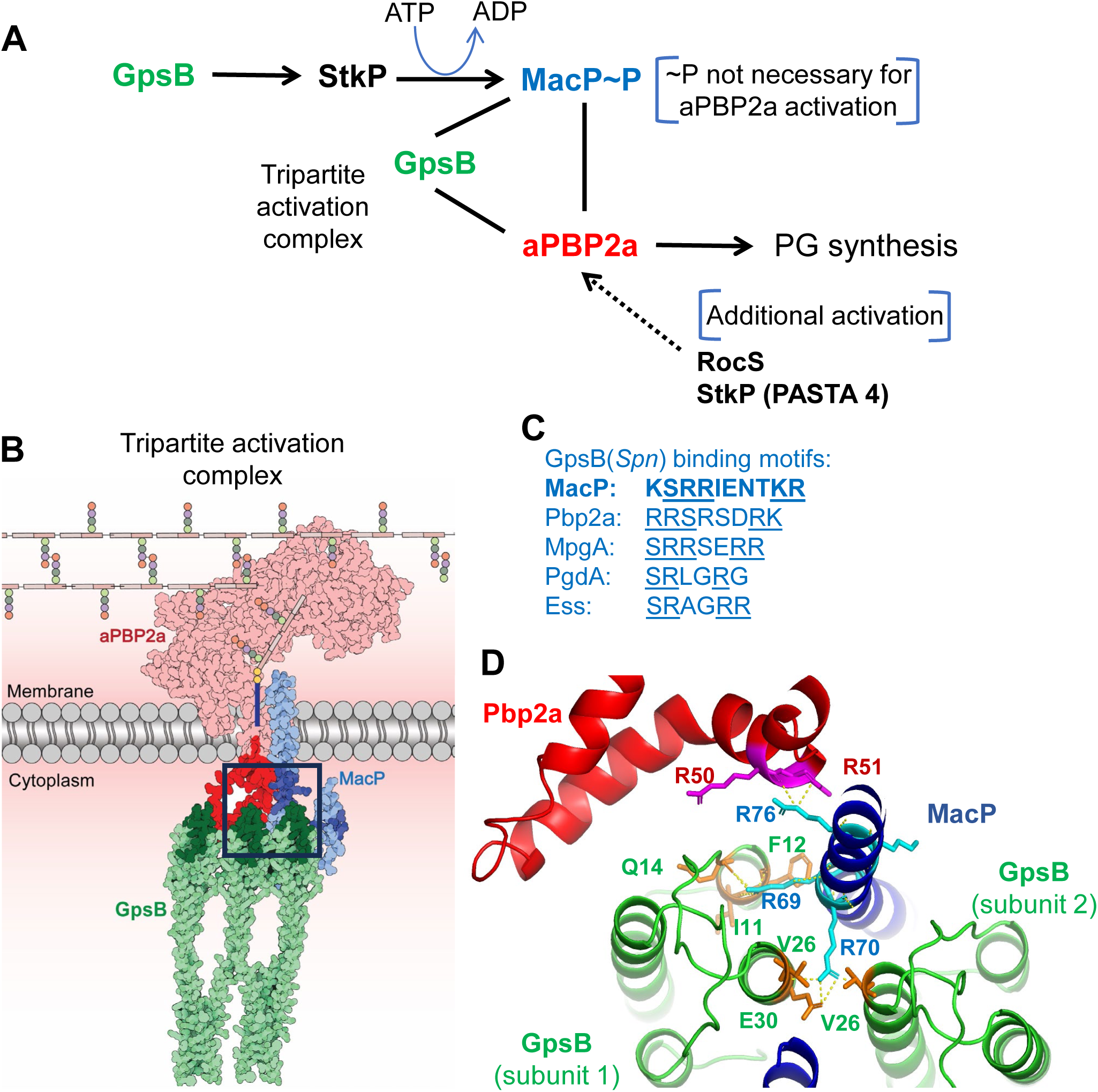
Summary of tripartite model of aPBP2a activation by MacP and GpsB. **(A)** GpsB acts as an adapter that facilitates interaction between MacP and aPBP2a to stimulate aPBP2a PG synthase activity in *Spn*. GpsB also activates the Ser/Thr protein kinase activity of StkP to phosphorylate Thr32 and Thr56 of MacP; however, MacP phosphorylation is not required for aPBP2a activation in transformation assays or during exponential growth of *Spn* cells. Besides MacP and GpsB, genetic evidence suggests aPBP2a is regulated by additional direct or indirect mechanisms that involve RocS, protein phosphorylation, and the P4-PASTA domain of StkP. See text for additional details. **(B)** Model structure of tripartite complex formed between a hexamer of GpsB and monomers of MacP and aPBP2a predicted by AlphaFold3. The complex structure is of low global reliability (pTM = 0.61 for MacP-aPBP2a-GpsB and pTM = 0.47 for MacP-aPBP2a-6XGpsB), where the cytoplasmic region of MacP is itself predicted to be largely unstructured (**Fig. 1A**). In this structure, GpsB hexamer is a trimer of three dimers, whose amino termini each have the capacity to bind to six GpsB-binding motifs in different target proteins (Cleverley *et al*., 2019). Here, GpsB simultaneously binds to the GpsB-binding motifs in the amino terminus of aPBP2a (**Fig. 7A**) and in the conserved membrane-proximal domain of MacP (**Fig. 7C**, **8A**, and **9C**). There are additional important functional interactions in the TM interface of aPBP2a and MacP (**Fig. 4**). **(C)** Comparison of the spaced Arg residues in the GpsB-binding motif of MacP reported here (**Fig. 8A**) with those in aPBP2a, MpgA, PgdA, and Ess (Cleverley *et al*., 2019, Millat *et al*., 2026). **(D)** The AlphaFold3 prediction of the structure of the interface between the N-terminal domain of GpsB and MacP corresponds to the GpsB-binding motif of MacP determined experimentally here (**Fig. 8**).

To demonstrate decreased binding of MacP(S68A R69A R70A K75A R76A) to GpsB, we performed co-IP experiments analyzed by quantitative western blotting (**Fig. 8D** and **S9D**). Twice as much F-MacP(S68A R69A R70A K75A R76A) extract compared to WT extract was used as bait in quantitative western blotting to compensate for the reduced expression of the mutant protein (**Fig. S9D**). Compared to WT MacP, the relative amount of GpsB or aPBP2a bound to MacP(S68A R69A R70A K75A R76A) dropped about 17-fold or 5-fold, respectively (**Fig. 8D** and **8E**). This result supports the conclusion that the MacP(68-KS<u>RR</u>IENTK<u>R</u>-76) motif mediates an interaction with GpsB. In addition, decreased binding between MacP and GpsB seems to decrease MacP binding to aPBP2a, consistent with GpsB acting as an adapter that promotes optimal activation of aPBP2a activity by MacP.

### 2.8. *phpP* or *rocS* mutants are synthetically lethal with Δ*pbp1a*

We reexamined Tn-Seq data of a Δ*pbp1a* mutant during exponential growth in BHI broth for other possible regulators of aPBP2a. Insertions in two genes, *phpP* and *rocS*, were absent in a Δ*pbp1a* mutant compared to WT (**Fig. S10** and **S11**, respectively). *phpP* encodes the sole Ser/Thr protein phosphatase in *S. pneumoniae* and is transcribed in an operon upstream of *stkP*, which encodes its cognate StkP Ser/Thr protein kinase (**Fig. S10A**). We reported previously that a Δ*phpP::P_c_-erm* mutation was polar and reduced StkP kinase expression; consequently, Δ*phpP::P_c_-erm* mutations led to a decrease in cellular protein phosphorylation, instead of an expected increase (Rued *et al*., 2017). The Δ*phpP::P_c_-erm* mutant grew slower than WT (Rued *et al*., 2017), which may account for the sparser number of insertions in *phpP* compared to non-essential genes in the WT strain in Tn-Seq (**Fig. S10B**). In comparison, there were no insertions in *phpP* in the Δ*pbp1a* mutant (**Fig. S10B** and **S10C**). This synthetic lethality was confirmed by transformation assays where the Δ*phpP::P_c_-erm* allele could be moved into a WT strain, but not into a markerless Δ*pbp1a* mutant (**Fig. S10D**). Tn-Seq also indicated that the P4-pasta domain at the C-terminus of StkP was required for growth in the Δ*pbp1a* or Δ*macP* mutant, whereas P4 was not essential in the WT or other mutant strains (**Fig. S10B**).

Together, these results suggested that phosphorylation of proteins other than MacP could play a role in aPBP2a regulation. To test this idea, we transformed Δ*pbp1a* into a suppressed Δ*stkP* strain lacking Ser/Thr protein phosphorylation. Previously, we reported that the essentiality of StkP in growing D39 progenitor cells was due to its phosphorylation of IreB (Tsui *et al*., 2023). When unphosphorylated, IreB acts as a negative regulator of the enzymatic activity of redundant MurZ and MurA that catalyze the first committed step of PG precursor biosynthesis. Consequently, Δ*stkP* is suppressed by mutations in this pathway, such as *murZ*(D280Y), resulting in growth similar to that of the WT strain (Tsui *et al*., 2023). We found that Δ*macP* transformants of a *murZ*(D280Y) Δ*stkP* mutant formed WT-sized colonies (row 2, **Fig. S10E**), indicating that aPBP2a activity is not required in the absence of protein phosphorylation. Δ*pbp1a* transformants of a *murZ*(D280Y) Δ*stkP* mutant formed noticeably smaller colonies than WT that varied in size with the antibiotic selection (rows 2 and 3, **Fig. S10E**). Notably, the *murZ*(D280Y) Δ*stkP* Δ*pbp1a* mutant was not transformable with Δ*macP* (row 3, **Fig. S10E**). These results reinforce the conclusion that phosphorylation of MacP is not required for aPBP2a activation and that phosphorylation of a regulator other than MacP may contribute to aPBP2a activation.

Finally, *rocS* showed a synthetic lethal relationship with Δ*pbp1a* and possibly with Δ*macP* in Tn-Seq screens (**Fig. S11A**). RocS is a critical regulator that couples cell division to chromosome segregation in *S. pneumoniae* (Mercy *et al*., 2019). Absence of RocS results in poor growth in culture and few insertions in Tn-seq. A synthetic lethal relationship between Δ*rocS* and Δ*pbp1a*, but not Δ*macP*, was confirmed by transformation assays (**Fig. S11B**). The synthetic lethality of *phpP* or *rocS* mutations with Δ*pbp1a* in the context of PBP2a regulation is discussed below.

## 3. DISCUSSION

Previous work concluded that *Spn* aPBP2a activity is positively regulated by MacP phosphorylated at Thr32 (Fenton *et al*., 2018) and by GpsB (Rued *et al*., 2017), which is an adapter protein involved in PG synthesis and cell division in Gram-positive bacteria (Cleverley *et al*., 2019). Because GpsB also acts as a positive regulator of the StkP Ser/Thr protein kinase (**Fig. 9A**) (Stauberová *et al*., 2024, Rued *et al*., 2017), one model is that GpsB activation of aPBP2a is indirect and occurs by GpsB positively regulating StkP phosphorylation of MacP (Fenton *et al*., 2018, Stauberová *et al*., 2024). In addition, recent phosphoproteomic studies showed that MacP is phosphorylated in growing cells at both Thr32 and Thr56 as well as at Thr7 and Thr37 under stress conditions (Hirschfeld *et al*., 2019, Ulrych *et al*., 2021). We set out to confirm and quantitate MacP phosphorylation at Thr32 and Thr56 and test whether positive regulation of aPBP2a by GpsB was indirect through GpsB activation of the StkP Ser/Thr protein kinase.

We found that ≈41% of MacP was phosphorylated at both Thr32 and Thr56 and ≈36% of MacP was unphosphorylated in exponentially growing *Spn* cells (**Fig. 2A** and **S3H**). Moreover, MacP phosphorylated at both Thr32 and Thr56 was detected in biochemical reactions containing purified proteins (**Fig. 2C**). Remarkably, our results showed that phosphorylation of MacP at Thr32 and/or Thr56 was not required for activation of aPBP2a in transformation assays or growing *Spn* cells (**Fig. 1B** and **S3D**). Two pieces of evidence supported this conclusion. Phosphoablative *macP*(T32A and/or T56A) mutations were not synthetically lethal with a Δ*pbp1a* mutation in direct transformation assays, where the numbers and colony morphology of the mutant matched those of the WT strain (**Fig. 1B** and **S4A**). This result indicates that unphosphorylated MacP activates aPBP2a in the absence of aPBP1a. Likewise, phosphomimetic *macP* (T32E and/or T56E) mutations were not synthetically lethal with Δ*pbp1a*.

Additional structure-function studies showed that internal deletions in the cytoplasmic domain of MacP(Δ5-8; Δ21-58 (including Thr32 and Thr56); Δ30-33; or Δ46-53), (**Fig. 3B**) were not synthetically lethal with Δ*pbp1a* in transformation and growth assays (**Fig. 3B, S4A**, and **S5C**). Of these deletions, only *macP*(Δ21-58) destabilized MacP appreciably (**Fig. 3C**). But the remaining MacP(Δ21-58), which amounted to ≈11% of the WT level (**Fig. 3D**), was sufficient to activate aPBP2a similar to WT MacP in transformation (**Fig. 3B** and **S4A**) and growth assays (**S5C**) in the absence of aPBP1a. MacP(Δ21-58) also maintained interactions with aPBP2a or GpsB in co-IP experiments (**Fig. S4A** and **S5A**). Parenthetically, this aPBP2a activation at a low cellular amount of MacP(Δ21-58) suggests that WT MacP is present in excess in *Spn* cells. In contrast, MacP(N77 stop) or MacP(L87 stop) truncated in the TM region was unstable and not detected in western blots (**Fig. 3C** and **3D**). We conclude that phosphorylation of MacP and most of the N-terminal cytoplasmic domain are not required for aPBP2a activation in the absence of aPBP1a under the conditions tested.

These findings do not support the conclusion that MacP phosphorylation is required for aPBP2a activation in the absence of aPBP1a (Fenton *et al*., 2018). Our study and this previous study were performed in the same unencapsulated derivative of the D39W progenitor strain, which lacks many spontaneous mutations that affect PG synthesis and cell division in R6, Rx1, and R800 laboratory strains (Trouve *et al*., 2021, Tsui *et al*., 2023, Rued *et al*., 2017). Therefore, the conflicting conclusions may reflect the different bacterial constructs and assays used. In (Fenton *et al*., 2018), synthetic lethality was detected for a phosphoablative GFP-MacP(T32A) fusion protein ectopically expressed in a strain in which aPBP1a was ectopically depleted; raising the possibility that proteins were not produced at WT levels, that the GFP fusion somehow interfered with the function of MacP mutant proteins, or that the depletion on plates caused a stress condition. Here, unmarked chromosomal *macP* mutations were assayed directly by transformation and growth assays, which reiterated results from Tn-seq experiments (**Fig. S2**). Moreover, the previous study discussed two results that were apparently not consistent with a requirement of MacP phosphorylation for aPBP2a activation. Phosphomimetic GFP-*macP*(T32E) was synthetically lethal with Δ*pbp1a*, instead of being synthetically viable as expected (Fenton *et al*., 2018). Incongruously, *macP*(WT) and *macP*(T32A or T32E) alleles were viable upon aPBP1a depletion in a Δ*stkP* mutant, which lacked the sole pneumococcal Ser/Thr protein kinase (Fenton *et al*., 2018), but likely contained a Δ*stkP* suppressor mutation (Rued *et al*., 2017, Tsui *et al*., 2023). Together, these results indicate that phosphorylation of MacP *per se* is not required for activation of aPBP2a in transformation assays or in unstressed, growing *Spn* cells.

Further structure-function studies were performed to characterize the activation of aPBP2a by MacP. We used AlphaFold3 to model interacting amino acids in the TM domain interface of MacP or the juxtamembrane region of aPBP2a (**Fig. 4, S1C, S4A**, and **S4B**). MacP(L83A I86A L87A) or aPBP2a(R51A H53A K56A) abolished aPBP2a activation in a Δ*pbp1a* mutant (**Fig. 5A, S4A**, and **S4B**). This loss of activation was not correlated with a large decrease in the cellular amounts of MacP(L83A I86A L87A) (≈46%) or aPBP2a(R51A H53A K56A) (≈140%) compared to WT MacP (**Fig. 5B**-**5D**). Instead, the amino acid changes in MacP(L83A I86A L87A) abolished interaction with aPBP2a by ≈95% (**Fig. S7E**), while not causing mislocalization of MacP(L83A I86A L87A) from midcell (**Fig.S7F**). Taken together, these results demonstrate that certain amino acids in the MacP(TM) interface (e.g., MacP(L83 I86 L87)) and the aPBP2a juxtamembrane region (e.g., aPBP2a(R51 H53 K56)) are required for MacP activation of aPBP2a, and in the case of MacP(L83 I86 L87) interaction with aPBP2a (**Fig. 9B** and **9D**).

Finally, MacP(M101A L103A), with two amino acid changes in the short extracellular domain of MacP (**Fig. 5A** and **S1D**), was produced at ≈90% of the WT MacP amount (**Fig. 5B** and **5D**). MacP(M101A L103A) still activated aPBP2a in the absence of aPBP1a (**Fig. 5A** and **S4A**) and showed only a slight (<30%) reduction in binding to aPBP2a in co-IP assays (**Fig. S7E**). Notably, an AlphaFold3 model predicted that the short extracellular peptide of MacP containing M101 and L103 interacts with the regulatory domain of aPBP2a containing A77 (**Fig. 4A**), which mediates MacP activation of aPBP2a GTase activity (Midonet *et al*., 2023). The modestly decreased interaction of MacP(M101A L103A) with aPBP2a is consistent with this interaction.

We next turned to how GpsB positively regulates aPBP2a activity, given that phosphorylation of MacP is not required. B2H assays confirmed that GpsB interacted with MacP (**Fig. 6A**), and co-IP assays showed that GpsB and MacP were in a complex at some stage of the cell cycle (**Fig. 6B** and **6C**) (Stauberová *et al*., 2024). As expected from previous results (Fenton *et al*., 2018, Cleverley *et al*., 2019), GpsB or MacP interacted with aPBP2a in both assays. Unexpectedly, MacP interacted with members of the divisome (EzrA, FtsL, FtsQ, and bPBP2x), class A aPBP1a but not aPBP1b, members of the PG elongasome (bPBP2b, RodA, RodZ, MreC), and the MpgA muramidase in B2H assays (**Fig. 6A**). Co-IP assays confirmed complex formation between MacP and bPBP2x, bPBP2b, MreC, RodZ, and MpgA (**Fig. 6B**-**6D** and **Table S4**) at unknown stages of the cell cycle, including the initial FtsZ ring that contains all proteins required for sPG and ePG synthesis (Briggs *et al*., 2021, Massidda *et al*., 2013, Vollmer *et al*., 2019). We focused on the interaction between MacP and GpsB in the activation of aPBP2a, leaving investigation of other MacP interactions for future studies.

We showed before that GpsB interacts with an Arg-rich domain (5’-<u>RR</u>S<u>R</u>SD<u>R</u>K-3’) in the cytoplasmic N-terminal region of *Spn* aPBP2a (**Fig. 7A** and **S1D**) (Cleverley *et al*., 2019). As expected, short amino-acid deletions in this region in aPBP2a diminished binding to GpsB in B2H assays (**Fig. 7A**), but minimally changed binding of aPBP2a to MacP, indicating that the Arg-rich N-terminal domain of aPBP2a did not mediate binding to MacP. Conversely, amino acids in the N-terminal domain of GpsB dimers required for binding to the Arg-rich motif in aPBP2a were also required for binding to MacP (**Fig. 7B**), albeit with somewhat different strengths. Reexamination of the MacP sequence disclosed a cytoplasmic Arg-rich sequence reminiscent of the GpsB-binding motif of aPBP2a right before the MacP(TM) domain (**Fig. 7C** and **S1D**).

A pentuple mutant of MacP(S68A R69A R70A K75A R76A) that changed five amino acids in the Arg-rich region (**Fig. 8A**) grew slowly when aPBP1a was depleted or absent (**Fig. 8BA, S4A**, and **S9C**), and the *macP*(S68A R69A R70A K75A R76A) Δ*pbp1a* mutant exhibited defects in cell morphology and division different from those caused by Δ*pbp1a* alone (**Fig. 8C**). The amount of MacP(S68A R69A R70A K75A R76A) in growing cells was lower (≈34%) than that of WT (**Fig. S9A** and **S9B**), but above the amount (≈11%) of MacP(Δ21-58) (**Fig. 3D**) sufficient to stimulate aPBP2a activity in a Δ*pbp1a* mutant (**Fig. 3B** and **S5C**). Notably, co-IP experiments demonstrated that MacP(S68A R69A R70A K75A R76A) bound GpsB or aPBP2a at only 6% or 20% of the WT level, respectively (**Fig. 8D**, **8E**, and **S9D**).

Taken together, these results support a model in which both GpsB and MacP can bind separately to aPBP2a, and GpsB acts to adapt or position MacP and aPBP2a together to stimulate aPBP2a activity (**Fig. 9A**). In this model, the Arg-rich motif of aPBP2a (amino acids 30-40; **Fig. 7A**) interacts with one N-terminal domain of the GpsB hexamer (Cleverley *et al*., 2019), while the Arg-rich motif of MacP (amino acids 67-76; **Fig. 7C** and **8A**) interacts with a second N-terminal domain in the same GpsB hexamer (**Fig**. **9B**). Consistent with this interpretation, AlphaFold3 modeling predicted a possible interaction between the N-terminal domain of GpsB and the Arg-rich motif of MacP that had be been changed in the pentuple mutant (**Fig**. **9D**); however, this structure was of low overall reliability (pTM = 0.47) partly because of difficulty modeling the unstructured N-terminal domain of MacP (**Fig. 1A**) and the hexamer structure of GpsB. Further work is required to determine the order and strength of binding GpsB to MacP or aPBP2a and whether binding of one pair influences the binding of the other pair. Another conclusion from this study is that there is variability in the Arg-rich, GpsB-binding motif sequence; therefore, GpsB may act as an adapter for additional PG-synthesis and cell-division proteins from those previously identified that contain iterations of the original GpsB-binding consensus motif (**Fig. 9C**) (Cleverley *et al*., 2019, Millat *et al*., 2026). Potential Arg-rich, GpsB-binding motifs in other proteins reported here to interact with GpsB (**Fig. 6A**) include: N<u>R</u>FKKS in MreC, N<u>RRR</u>VG in bPBP2x, L<u>R</u>LIKY in aPBP1a, and L<u>R</u>LA<u>R</u> in RodZ (Richard J. Lewis, personal communication).

The conclusion that phosphorylation of MacP does not act as a switch to activate aPBP2a activity in *Spn* cells grown under standard conditions aligns with the lack of phenotypes of Δ*pbp2a* mutants reported so far (Land *et al*., 2013, Tsui *et al*., 2016, Fenton *et al*., 2018, Cleverley *et al*., 2019). It remains to be determined whether aPBP2a activity and its regulation by MacP phosphorylation are required in cells subjected to antibiotic or other stresses, which greatly increase protein phosphorylation levels (Beilharz *et al*., 2012, Ulrych *et al*., 2021, Stauberová *et al*., 2024). Moreover, Tn-seq and corroborative transformation results reported here suggest that aPBP2a activity may be regulated by additional mechanisms. The PhpP Ser/Thr protein phosphatase or ultimate (P4) PASTA domain of the cognate StkP Ser/Thr protein kinase were synthetically lethal in the absence of aPBP1a (**Fig. S10B** and **S10D**). Given that insertion mutations in *phpP* reduce protein phosphorylation through polarity on *stkP* (Rued *et al*., 2017), these results suggest that phosphorylation of proteins other than MacP may be required for aPBP2a activity under these growth conditions.

Consistent with this idea, Δ*pbp1a* transformants of a suppressed *murZ*(D280Y) Δ*stkP* mutant, which lacked Ser/Thr protein phosphorylation, formed much smaller colonies compared to WT in the presence of different antibiotics (**Fig. S10E**). This result supports the conclusion that phosphorylation of MacP is not obligatory for aPBP2a activation. The slower growth is also consistent with another regulator of aPBP2a that requires phosphorylation. However, this second regulation would still depend on MacP, because no transformants were recovered when Δ*macP* was transformed into the *murZ*(D280Y) Δ*stkP* Δ*pbp1a* mutant (**Fig. S10E**). A complication in interpretating these experiments is that suppression of Δ*stkP* requires altering regulation of the PG precursor pathway (Tsui *et al*., 2023).

One candidate for a phosphorylated activator of aPBP2a, other than MacP, is the aPBP2a-coactivator GpsB, which is phosphorylated in growing and stressed *Spn* cells (Stauberová *et al*., 2024). A second candidate is RocS (Mercy *et al*., 2019, Demuysere *et al*., 2024), whose absence is synthetically lethal with Δ*pbp1a*, but not with Δ*macP* (**Fig. S11A** and **S11B**). The *rocS* gene is located one gene downstream from *macP* (**Fig. S1A**), and *macP* insertion mutations showed polarity consistent with a multigene operon, possibly including *rocS* (**Fig. S3C**). RocS is a phosphorylated, membrane-bound multimer that mediates chromosome partitioning in unencapsulated and encapsulated *Spn* cells (Mercy *et al*., 2019, Demuysere *et al*., 2024). In encapsulated strains, RocS additionally interacts with the CpsD regulator to coordinate FtsZ constriction with capsule secretion, thereby protecting the nucleoid from truncation (Mercy *et al*., 2019). To carry out these roles, the helix-turn-helix domain of RocS binds DNA, and RocS binds to cell division protein FtsZ, chromosome partitioning protein ParB, and chromosome maintenance protein Smc (Mercy *et al*., 2019, Demuysere *et al*., 2024). The genetic results presented here suggest that RocS could potentially regulate aPBP2a to coordinate PG synthesis with chromosome segregation.

Finally, it is noteworthy that the ultimate (P4) PASTA domain of the StkP Ser/Thr protein kinase was synthetically lethal with both aPBP1a or MacP, but not aPBP1b or aPBP2a in an unencapsulated derivative of a serotype-2 D39 strain (**Fig. S10B**). This result suggests a possible requirement of the StkP protein for aPBP2a activation by MacP. The P4 PASTA domain of StkP has been implicated in septal localization of StkP and interaction with the LytB PG hydrolase that separates fully divided *Spn* cells (Righino *et al*., 2018, Zucchini *et al*., 2018, Martínez-Caballero *et al*., 2023). In contrast, the essential P1-P3 PASTA domains bind to peptidoglycan fragments, β-lactam antibiotic ligands (Maestro *et al*., 2011), and possibly Lipid II (Hardt *et al*., 2017, Sun *et al*., 2023) and are required for activation of StkP (Novakova *et al*., 2010, Pensinger *et al*., 2018). Phenotypes of P4 PASTA domain mutants were observed in derivatives of laboratory strain R6 (Zucchini *et al*., 2018), which contains numerous mutations compared D39 progenitor strains (Tsui *et al*., 2023), and it remains to be determined whether these phenotypes are observed in D39 strains. In this regard, *murZ*(D280Y) Δ*stkP* mutants form wild-type shaped cells in exponentially growing Δ*cps* D39 strains (Tsui *et al*., 2023), indicating that StkP Ser/Thr protein is not obligatorily required for LytB function. How aPBP2a is regulated by RocS, StkP, and other proteins that are in complexes with MacP at some stage of cell division (**Fig. 6**) awaits further investigation in growing and stressed *Spn* cells.

## 4. EXPERIMENTAL PROCEDURES

### 4.1. Bacterial strains and growth conditions

*Spn* strains used in this study are listed in **Table S1**. All strains were derived from IU1824 (D39 Δ*cps rpsL1*) or IU1945 (D39 Δ*cps*), which are unencapsulated derivatives of *Spn* serotype-2 progenitor strain D39W (Lanie *et al*., 2007, Slager *et al*., 2018). Strains containing antibiotic markers were constructed by transformation of CSP1-induced competent pneumococcal cells with linear DNA amplicons synthesized by overlapping fusion PCR (Tsui *et al*., 2014). Strains containing markerless alleles in native chromosomal loci were constructed using allele replacement via the Janus cassette (P_c_-[*kan*-*rpsL*^+^]) (Sung *et al*., 2001) or Sweet Janus cassette (*sacB*-*kan*-*rpsL*^+^) (Echlin & Rosch, 2020). Primers used to synthesize different amplicons for strain constructions are listed in Table S2.

D39 strains were grown on plates containing trypticase soy agar II (modified; Becton-Dickinson) or Columbia agar and 5% (v/v) defibrinated sheep blood (TSAII-BA) at 37°C in an atmosphere of 5% CO_2_. TSAII-BA or Columbia agar plates for selections contained antibiotics at concentrations reported previously (Tsui *et al*., 2014, Tsui *et al*., 2016). Bacteria were cultured statically in Becton-Dickinson brain heart infusion (BHI) broth at 37°C in an atmosphere of 5% CO_2_, and growth was monitored by OD_620_, as described before (Tsui *et al*., 2016). Mutant constructs were confirmed by PCR and DNA sequencing of chromosomal regions corresponding to the amplicon region used for transformation. Ectopic expression of various genes was achieved from a P_Zn_ zinc-inducible promoter at the ectopic *bgaA* site (Tsui *et al*., 2014, Tsui *et al*., 2016). 0.2 to 0.5 mM concentrations of ZnCl_2_ containing 1/10 concentration of MnSO_4_ (Zn^2+^/(1/10)Mn^2+^) were added to TSAII-BA plates or BHI broth for inducing conditions. Mn^2+^ was added to Zn^2+^ conditions to prevent zinc toxicity (Tsui *et al*., 2023).

In all experiments, cells were inoculated from frozen glycerol stocks into BHI broth, serially diluted, and incubated for 12–15 h statically at 37°C in an atmosphere of 5% CO_2_. Parallel cultures were set up for each strain and condition to generate growth curves and collect samples for western blotting or microscopy. The next day, cultures at OD_620_ ≈0.1–0.4 were diluted to OD_620_ ≈0.003 in BHI broth with or without additional Zn^2+^/(1/10)Mn^2+^as indicated. Doubling times of exponential growth were determined from linear regions on semi-log plot using GraphPad Prism. Maximum growth yields were determined by the highest OD_620_ values reached on linear plots within 8-10 h of growth. Doubling times and maximal growth yields were compared to WT strains using one-way ANOVA analysis (GraphPad Prism, Dunnett’s test). Cultures were sampled for microscopy or western blot analysis at OD_620_ ≈0.1–0.2, which corresponded to early to mid-exponential phase.

### 4.2. Plasmid construction and site-directed mutagenesis for biochemical experiments

Plasmids constructed to produce proteins for biochemical studies are listed in **Table S3**. pETPHos-*macP*ΔTM and derived plasmids were used to express recombinant His6-MacPΔTM in *E. coli*. To generate the expression plasmid pETPHos-*macP*ΔTM, we amplified *macP*ΔTM (aa 1-85) using the primer pair LN385/LN386. The PCR product was cloned into the *Nde*I/*Bam*HI restriction sites of the modified pETPhos vector, named pETPhosLink (Novakova *et al*., 2010), under the control of the T7 promoter so that the protein was fused to a His-tag at the N-terminus. pETPhos-macPΔTM carrying mutations T32A, T56A, or T32A/T56A (2TA) was produced by site-directed mutagenesis using the mutagenic primers listed in Table S2.

### 4.3. Transformation assays

Transformation assays were performed as described previously (Tsui *et al*., 2023). For most transformation assays, Δ*pbp1a*::P_c_-*erm* and Δ*pbp2a*::P_c_-*erm* were used as test amplicons and Δ*pbp1b*::P_c_-*erm* was used as a control amplicon. All amplicons had ≈1 kb of flanking chromosomal DNA on each end. Amplicons were synthesized by PCR using the primers and templates listed in **Table S2.** Transformation experiments also included negative control without added DNA. The Δ*pbp1b*::P_c_-*erm* positive control or no DNA negative control showed >500 or no colonies, respectively and are not included in data tables. Volumes of transformation mixtures plated (50 to 300 µL) were adjusted to produce ≈150 to 500 colonies with the Δ*pbp1b*::P_c_-*erm* control amplicon. Transformations with positive control amplicons typically yielded >500 colonies per 1 mL of transformation mixture. The transformants were confirmed by PCR. Each transformation experiment was performed independently two or more times. The sizes of the colonies indicated are relative to colonies of the recipient strains transformed with the control Δ*pbp1b*::P_c_-*erm* amplicon. For transformation requiring induction of gene expression from the P_Zn_ zinc-inducible promoter in the ectopic *bgaA* site, 0.2 mM ZnCl_2_ + 0.02 mM MnSO_4_, or 0.4 mM ZnCl_2_ + 0.04 mM MnSO_4_ were added to transformation mixes and to the soft agar spread on blood plates (Rued *et al*., 2017). Photos of colony morphologies were taken from transformation plates after 20 h of incubation at 37°C with the illumination source under the plates. For experiments in **Fig. S10E** using *murZ*(D280Y) Δ*stkP*::P_c_-*erm* mutants, Δ*pbp1a*::P_c_*-*[*kan-rpsL^+^*], Δ*pbp2a*::P_c_*-*[*kan-rpsL^+^*], and Δ*pbp1b*::P_c_-*aad9* amplicons were used in transformation assays using the same protocol as described above with Δ*bgaA*::P_c_*-*[*kan-rpsL^+^*] and Δ*bgaA*::P_c_-*aad9* as the positive control amplicon.

### 4.4. Cell length and width measurements

Cell lengths and widths of bacterial cells growing exponentially in BHI broth were measured as previously described (Tsui *et al*., 2016). Unless indicated in the figure legends, more than 50 cells from at least 2 independent experiments were measured and plotted as scatter plots. *p* values were obtained by one-way ANOVA analysis using the nonparametric Kruskal-Wallis test in the GraphPad Prism.

### 4.5. Quantitative western blotting

Cell lysates were prepared using SEDS lysis buffer (0.1% deoxycholate (vol/vol), 150 mM NaCl, 0.2% SDS (vol/vol), 15 mM EDTA, pH 8.0), and western blotting was performed as previously described (Lamanna *et al*., 2022). Briefly, bacteria were grown exponentially in 5 ml BHI broth to an OD_620_ ≈ 0.15–0.2. Frozen pellets collected from 1.8 mL cultures at OD_620_ ≈0.16 were suspended in 80 μL of SEDS lysis buffer. The volume of the SEDS buffer was adjusted proportionally to the OD_620_ values. The μL amounts of protein lysates loaded in each lane are listed for each blot. The sources of antibodies used for western blotting are as follows. Primary antibodies were polyclonal rabbit antibodies: anti-FLAG (Sigma, F7425, 1:1,400), anti-pThr (Cell Signaling, #9381) (Rued *et al*., 2017), and anti-PBP2a (1:5,000) (Cleverley *et al*., 2019). Secondary antibodies were anti-rabbit IgG conjugated to horseradish peroxidase (GE Healthcare NA93AV, 1:10,000), or Licor IR Dye800 CW goat anti-rabbit (926–32,211, 1:14,000). Chemiluminescence signals obtained with secondary HRP-conjugated antibody and IR signals obtained with the Licor IR Dye800 CW secondary antibody were detected using Azure Biosystems 600, as described previously (Lamanna *et al*., 2022).

Relative expression levels of MacP were determined using FLAG-MacP expressed from its native chromosomal locus. To ensure linearity of western blot signal values versus. protein amounts, a range of amounts of (5.0, 7.5, 10.0, or 15.0 µL) protein samples of IU17032 (FLAG-*macP*) or IU1824 (*pbp2a^+^*) were loaded on the same gel as the experimental samples to provide a standard curve of µL protein amounts versus signal intensities (**Fig. 3E**) (Perez *et al*., 2024). These plots were obtained for each western quantitation experiment and were used to calculate the relative protein amounts in each sample lane by interpolation in the linear range of detection. TotalStain-Q (Azure Biosystems, AC2227) total protein stain was used to determine the relative amount of total protein in each lane and was used for normalization. Protein concentrations were not determined. Signal intensities obtained with anti-FLAG or anti-PBP2a antibodies were normalized to total protein stain of each lane stained with Totalstain-Q reagent Standard curves of dilutions of protein staining were generated to verify signal linearity (Perez *et al*., 2024). All experiments were performed at least twice independently.

### 4.6. Phos-tag SDS-PAGE and western blotting

Phos-tag SDS-PAGE to distinguish protein phosphorylation states of FLAG-MacP and standard western blotting were carried out as described previously with certain modifications (Tsui *et al*., 2014, Wayne *et al*., 2012). To determine the phosphorylation state of the FLAG-MacP protein, overnight BHI broth cultures were diluted and grown to OD_620_ ≈0.2 in 30 mL of BHI. Cells were centrifuged at 14,500 × g for 5 min at 4°C, and all subsequent steps were performed at 4°C. Pellets were lysed using a FastPrep homogenizer (MP biomedicals) in cold lysis buffer (20 mM Tris-HCl pH 7.0 and 1 protease inhibitor tablet (Thermo Fisher Scientific) per 10 mL buffer). Cell lysates were resolved by 12.5% SDS-PAGE supplemented with 75 μM Phos-Tag acrylamide (AAL-107; Wako) and 100 μM MnCl_2_, and standard 12.5 % SDS-PAGE was used as the control (Wayne *et al*., 2012). Samples were loaded after normalizing to OD_620_ (for OD_620_ ≈0.2; 10 µL of the sample was loaded). FLAG-MacP was detected by western blotting as described previously, using anti-FLAG (1:1,400) (Sigma, F7425) as the primary antibody (Tsui *et al*., 2014). Secondary antibodies were anti-rabbit IgG conjugated to horseradish peroxidase (GE healthcare NA93AV, 1:10,000). Chemiluminescent signal in protein bands was quantified using an IVIS imaging system, as described previously (Lamanna *et al*., 2022).

### 4.7. Co-immunoprecipitation (co-IP)

Co-immunoprecipitation (co-IP) was performed as previously described (Rued *et al*., 2017, Lamanna *et al*., 2022). Briefly, strains expressing FLAG-tagged proteins and control strains lacking tagged protein were grown exponentially in 400 mL of BHI and processed as previously described. Protein concentrations of samples were determined using the Bio-Rad DC^TM^ protein assay (Bio-Rad). 1 mL of lysate containing similar amounts of total protein (3-6 mg/mL) was added to tubes containing 50 μL anti-FLAG magnetic beads (Sigma, M8823). Equal amounts of protein were loaded onto the beads for strains expressing FLAG-tagged proteins and the corresponding control strains lacking FLAG-tagged proteins in each experiment. The protein was eluted from the beads with FLAG peptide, as previously described (Lamanna *et al*., 2022). 3-5 µL of input protein samples and 20 µL (unless otherwise specified in the figure legends) of each elution sample were separated by SDS-PAGE on 4-15% precast protein gels (Bio-Rad, Hercules, CA, USA) in Tris-glycine buffer. Gels were western blotted using rabbit anti-FLAG (1:1,400) (Sigma, F7425), anti-MreC (1:5,000) (Lamanna *et al*., 2022), anti-PBP2a (1:10,000), anti-PBP2b (1:10,000) (Lamanna *et al*., 2022), and anti-GpsB (1:5,000) as primary antibodies. Secondary antibody used was Licor IR Dye800 CW goat anti-rabbit (926–32,211, 1:14,000). IR signals obtained with the Licor IR Dye800 CW secondary antibody were detected using an Azure Biosystem 600 (Lamanna *et al*., 2022). Each experiment was independently performed at least twice unless specified in the legend. For co-IP LC-MS, magnetic beads containing cross-linked protein were submitted to the Laboratory for Biological Mass Spectrometry at Indiana University for protein identification.

To determine phosphorylation in vivo FLAG-MacP-WT, -T32A, -T56A and -2TA variants were immunoprecipitated as described above. 10 µL (=10 µg) of the input protein samples and 10 µL of each elution sample were separated by SDS-PAGE on 16% precast protein gels (Invitrogen) in Tris-glycine buffer. Gels were subjected to immunoblotting using custom made rabbit anti-MacP serum (1:20,000) (Stauberová, Kubeša et al., 2024) and rabbit anti-pThr (1:50,000) (Cell Signaling) to detect protein phosphorylation. Goat-anti-rabbit IgG IR700 (1:10,000; Advansta) was used as a secondary antibody. The fluorescent signal in the protein bands was detected using an AmershamTM Typhoon (GE Healthcare) instrument and quantified using ImageQuantTL software. The results are representative of four independent experiments.

### 4.8. 3D structure and residue alignment

Structures of MacP, PBP2a, and GpsB hexamer from *S. pneumoniae* D39 were predicted using AlphaFold3 (Abramson *et al*., 2024), and images were generated using PyMOL (Schrödinger, LLC). For amino acid sequence comparisons, amino acid sequences were obtained from the NCBI Protein database (https://www.ncbi.nlm.nih.gov/protein/) and aligned using the Clustal Omega web server. Protein–protein interaction interfaces between MacP and PBP2a were analyzed from the AlphaFold3-predicted structure, and residue–residue contacts were characterized using PDBsum.

### 4.9. Bacterial two-hybrid (B2H) assay

Plasmids used for B2H assays are listed in Table S3. The plasmids carrying *macP* WT were generated by amplifying with specific primers the corresponding gene by PCR from *Spn* D39 chromosomal DNA and cloning it into the respective XbaI/EcoRI sites of pKT25 and pUT18C to generate, respectively, the T25-MacP and T28-MacP hybrid proteins fused at the C-terminal ends of the T25 and T18 fragments (Stauberová *et al*., 2024). The plasmids carrying the phosphoablative or phosphomimetic *macP* variants, as well as the other mutated or truncated variants of *macP* or *pbp2a*, were generated by introducing the desired mutations or truncations *via* site-directed mutagenesis (SDM), using the SDM primers listed in Table S2. Plasmids carrying the mutated or truncated variants were transformed into the cloning host *E. coli* DH5a and transformants were selected on LB agar plates containing 50 μg/mL kanamycin (Kan^50^) or 100 μg/mL ampicillin (Amp^100^) and 0.4% glucose to repress leaky expression (Cleverley *et al*., 2019)The presence of the correct allele was verified by double-strand sequencing. B2H assay was performed as previously reported (Rued *et al*., 2017, Cleverley *et al*., 2019, Perez *et al*., 2019, Perez *et al*., 2021, Lamanna *et al*., 2022, Stauberová *et al*., 2024). Briefly, hybrid plasmids pairs were co-transformed into *E. coli cya-*BTH101 and co-transformants were spotted onto LB Amp^100^ Kan^50^, and 40 or 60 μg/mL 5-bromo-4-chloro-3-indolyl-β-D-galactopyranoside (Xgal^40^ or X-gal^60^) agar plates. Plates were inspected and photographed after 24 and 40 h at 30°C. The plasmid pairs pKT25/pUT18C and pKT25-zip/pUT18C-zip were used as negative and positive controls, respectively. All B2H experiments were repeated at least twice.

### 4.10. Expression and purification of recombinant proteins and in vitro protein phosphorylation

The kinase domain of StkP (StkP-KD) was expressed and purified by affinity chromatography using a His-tag, as described previously (Novakova *et al*., 2010). To express His-MacPΔTM and the corresponding mutant proteins, *E. coli* BL21 (DE3) was transformed with the expression plasmids pETPhos-macP-WT, -T32A, -T56A, and -2TA. The resulting expression strains were cultured at 30 °C until OD_600_ ≈0.6. Overproduction of recombinant proteins was induced by adding isopropyl-β-D-thiogalactopyranoside (IPTG) to a final concentration of 1 mM, and the cultures were then incubated at 16°C for 20 h. Recombinant proteins were purified at room temperature using Ni-nitrilotriacetic acid (NTA)-metal affinity resin (Qiagen), according to the manufacturer’s instructions. Purified proteins were dialyzed against a buffer containing 25 mM Tris-HCl (pH 7.5), 100 mM NaCl, and 10% (v/v) glycerol. The protein concentration was determined using a BCA protein assay (Thermo Fischer Scientific). The recombinant protein substrate (2 μg) was incubated with purified StkP-KD (0.3 μg) in reaction buffer containing 25 mM Tris-HCl (pH 7.5), 25 mM NaCl, 10 mM MnCl_2_, 10 μM ATP, and 37 kBq γ^32^P-ATP. The reaction was initiated by adding ATP and terminated after 30 m of incubation at 37°C by adding 5X SDS sample buffer. Protein samples were resolved by SDS-PAGE using 16% pre-cast Tris-Glycine gels (Invitrogen), and Coomassie blue-stained gel was exposed to a sensitive screen and scanned using Amersham Typhoon Biomolecular Imager in Phosphor imager mode (Cytiva). To quantify the phosphorylation signal, the images from four independent experiments were analyzed using ImageQuant TL software (GE Healthcare).

### 4.11. Tn-seq transposon library generation and insertion sequencing

Tn-seq was performed using the protocols described in (Lamanna *et al*., 2022). A transposon insertion library was generated for each of the following strains: WT D39 Δ*cps rpsL1* (IU1824), isogenic Δ*pbp1a* (IU14697), Δ*pbp1b* (IU14697), Δ*macP* (IU17032), and Δ*pbp2a* (IU13256), which were grown exponentially in BHI. Insertion data were visualized graphically using the Artemis genome browser (version 10.2) (Carver *et al*., 2011). Tn-seq primary data for the region between IG_SPD_0335 to IG_SPD_0340, SPD_0875 to SPD_0880, SPD_1541 to SPD_1544 and SPD_1820 to SPD_1823, are provided in Appendix S1. These regions include the *pbp1a*(SPD_0336) and *gpsB*(SPD_0339) loci, the *macP*(SPD_0876) and *rocS*(SPD_0878) loci, the *stkP*(SPD_1542) and *phpP*(SPD_1543) loci, and the *pbp2a*(SPD_1821) locus, respectively. Appendix S1 includes run summaries, the number of reads per TA site for each gene, and count ratios for each gene in the indicated mutants relative to WT. *p* values for comparisons of the number of reads per TA site in each gene were calculated by the Mann–Whitney test using GraphPad Prism (version 11.0).

## AUTHOR CONTRIBUTIONS

**Merrin Joseph:** Conceptualization; methodology; investigation; formal analysis; data curation; validation; writing–original draft. **Bohumil Kubeša:** Conceptualization; methodology; investigation; formal analysis; data curation; validation. **Ho-Ching T. Tsui:** Conceptualization; methodology; investigation; formal analysis; data curation; validation. **Mattia Benedet:** Conceptualization; methodology; investigation; formal analysis; data curation; validation. **Orietta Massidda.** Conceptualization; methodology; formal analysis; data curation; validation; supervision; funding acquisition. **Pavel Branny.** formal analysis; funding acquisition. **Linda Doubravová.** Conceptualization; methodology; formal analysis; data curation; validation; supervision; funding acquisition. **Malcolm E. Winkler.** Conceptualization; methodology; formal analysis; data curation; validation; supervision; funding acquisition.

## ACKNOWLEDGMENTS

We thank Irfan Manzoor for generating primary Tn-seq data, and Kevin Bruce and other members of the Winkler laboratory for discussions about this work. This work was supported by Czech Science Foundation grant 22-11062S, Ministry of Education, Youth, and Sports of the Czech Republic grant: Talking microbes – understanding microbial interactions within One Health framework CZ.02.01.01/00/22_008/0004597, and grant INTER-EXCELLENCE II, INTER-ACTION, LUAUS25092 (to PB), institutional research funds from the CIBIO Department at University of Trento (to OM), and NIH grant R35GM131767 (to MEW). Work done on the Carbonate Research supercomputer was supported in part by the Lilly Endowment, Inc., through its support of the Indiana University Pervasive Technology Institute.

## DATA AVAILABILITY STATEMENT

All data that supports the findings of this study are reported with indicated statistical analyses and numbers of biological repeats in this paper, including Supplemental Information, and the Appendix A Dataset containing primary data.

## ETHICS STATEMENT

This work did not include direct animal or human experimental subjects requiring formal approval or consent. Antibodies used in this study are available commercially, were published previously, or were prepared by companies approved by the Indiana University Bloomington Institutional Animal Care and Use Committee.

