## Supplemental_Information for "GpsB acts as an adapter for MacP-mediated activation of class A penicillin-binding protein aPBP2a in *Streptococcus pneumoniae*, independently of MacP phosphorylation"

**SUPPLEMENTAL TABLES: Tables S1-S4**

**SUPPLEMENTAL FIGURE LEGENDS: Figures S1-S11**

**SUPPLEMENTAL REFERENCES**

**SUPPLEMENTAL FIGURES: Figures S1-S11**

**Table S1.** *S. pneumoniae* and *E. coli* strains used in this study

| ***S. pneumoniae*** **strains** | | | |
|---|---|---|---|
| Strain number | Genotype (description)[a] | Antibiotic resistance[b] | Reference or source |
| IU1690 | D39W *cps*$^{+}$ | None | |
| IU1781 | D39W *cps*$^{+}$ *rpsL1* | Str$^{R}$ | (Lanie *et al.*, 2007) |
| IU1824 | D39W *rpsL1* Δ*cps2A'-cps2H'* = D39W *rpsL1* Δ*cps* | Str$^{R}$ | (Lanie *et al.*, 2007) |
| IU1945 | D39W Δ*cps2A'-cps2H'*= D39 Δ*cps* | None | (Lanie *et al.*, 2007) |
| E46 | D39 Δ*cps* Δ*bgaA*::P$_{c}$-*erm* | Erm$^{R}$ | (Rued *et al.*, 2017) |
| E177 | D39 Δ*cps* Δ*pbp1a*::P$_{c}$-*erm* | Erm$^{R}$ | (Land *et al.*, 2013) |
| E180 | D39 Δ*cps* Δ*pbp2a*::P$_{c}$-*erm* | Erm$^{R}$ | (Land *et al.*, 2013) |
| E193 | D39 Δ*cps* Δ*pbp1b*::P$_{c}$-*erm* | Erm$^{R}$ | (Land *et al.*, 2013) |
| E736 | D39 Δ*cps* Δ*phpP*::P$_{c}$-*erm* | Erm$^{R}$ | (Rued *et al.*, 2017) |
| E793 | D39 Δ*cps* Δ*macP*::P$_{c}$-*erm* (IU1945 X fusion Δ*macP*::P$_{c}$-*erm*) | Erm$^{R}$ | This study |
| K793 | D39 Δ*cps* Δ*macP*::P$_{c}$-[*kan-rpsL*$^{+}$] (IU1945 X fusion Δ*macP*::P$_{c}$-[*kan-rpsL*$^{+}$]) | Kan$^{R}$ | This study |
| K164 | D39W Δ*cps* Δ*pbp1a*::P$_{c}$-[*kan-rpsL*$^{+}$] | Kan$^{R}$ | (Tsui *et al.*, 2016) |
| K166 | D39W Δ*cps* Δ*pbp2a*::P$_{c}$-[*kan-rpsL*$^{+}$] | Kan$^{R}$ | (Tsui *et al.*, 2016) |
| IU4888 | D39 Δ*cps* Δ*gpsB*<>*aad9*//Δ*bgaA*::*kan*-t1t2-P$_{fcsK}$-*gpsB*$^{+}$ | Kan$^{R}$ Spec$^{R}$ | (Land *et al.*, 2013) |
| IU4970 | D39 Δ*cps* *mreC*-L-FLAG$^{3}$-P$_{c}$-*erm* | Erm$^{R}$ | (Land & Winkler, 2011) |
| IU6726 | D39 Δ*cps* *rpsL1* Δ*pbp1a*::P$_{c}$-[*kan-rpsL*$^{+}$] | Kan$^{R}$ | (Tsui *et al.*, 2016) |
| IU6741 | D39 Δ*cps* *rpsL1* Δ*pbp1a* markerless | Str$^{R}$ | (Tsui *et al.*, 2016) |
| IU7853 | D39 Δ*cps* *rpsL1* Δ*pbp2a*::P$_{c}$-[*kan-rpsL*$^{+}$] | Kan$^{R}$ | (Cleverley *et al.*, 2019) |
| IU7923 | D39 Δ*cps* Δ*stkP*::Pc-*erm* | Erm$^{R}$ | (Rued *et al.*, 2017) |
| IU9182 | D39 Δ*cps* *rpsL1* *gfp*-L1-*mapZ* markerless | Str$^{R}$ | (Perez *et al.*, 2019) |
| IU10063 | D39 Δcps Δ*bgaA*::*tet*-P$_{Zn}$-RBS$^{ftsA}$-*pbp2x* | Tet$^{R}$ | (Perez *et al.*, 2019) |
| IU12788 | D39 Δ*cps* *rpsL1* Δ*bgaA*::*kan*-t1t2-P$_{Zn}$-RBS$^{ftsA}$-*khpA* | Str$^{R}$ Kan$^{R}$ | (Zheng *et al.*, 2017) |

| | | | |
|---|---|---|---|
| IU13256 | D39 Δ*cps rpsL1* Δ*pbp2a* | Str$^R$ | (Cleverley *et al.*, 2019) |
| IU13444 | D39 Δ*cps rpsL1* Δ*pbp1a*::P$_c$-*erm* | Str$^R$ Erm$^R$ | (Cleverley *et al.*, 2019) |
| IU13604 | D39 Δ*cps rpsL1* Δ*ireB* markerless | Str$^R$ | (Tsui *et al.*, 2023) |
| IU13680 | D39 Δ*cps* Δ*pbp1b*::P$_c$-*aad9* | Spec$^R$ | (Tsui *et al.*, 2023) |
| IU14151 | D39 Δ*cps rpsL1 ireB*-L-FLAG$^3$-P$_c$-*erm* (IU1824 X fusion *ireB*-L-FLAG$^3$-P$_c$-*erm*) | Str$^R$ Erm$^R$ | This Study |
| IU14357 | D39 Δ*cps rpsL1* Δ*pbp1a*::P$_c$-*erm*//Δ*bgaA*::*tet*-P$_{Zn}$-RBS$^{ftsA}$-*pbp1a* | Tet$^R$ Erm$^R$ | (Land *et al.*, 2013) |
| IU14631 | D39 Δ*cps* Δ*macP*::P$_c$-*erm* Δ*pbp2a*::P$_c$-[*kan-rpsL$^+$*] (E793 X Δ*pbp2a*::P$_c$-[*kan-rpsL$^+$*] amplicon from K166) | Erm$^R$ Kan$^R$ | This study |
| IU14633 | D39 Δ*cps* Δ*macP*::P$_c$-[*kan-rpsL$^+$*] Δ*pbp2a*::P$_c$-*erm* (K793 X Δ*pbp2a*::P$_c$-*erm* amplicon from E180) | Erm$^R$ Kan$^R$ | This study |
| IU14650 | D39 Δ*cps rpsL1* Δ*macP*::P$_c$-[*kan-rpsL$^+$*] (IU1824 X Δ*macP*:: P$_c$-[*kan-rpsL$^+$*] amplicon from K793) | Kan$^R$ | This study |
| IU14697 | D39 Δ*cps rpsL1* Δ*pbp1b* | Str$^R$ | (Lamanna *et al.*, 2022) |
| IU14699 | D39 Δ*cps rpsL1* Δ*macP* markerless (IU14650 X fusion Δ*macP* markerless) | Str$^R$ | This study |
| IU14746 | D39 Δ*cps rpsL1* Δ*macP* Δ*pbp1b*::P$_c$-*erm* (IU14650 X Δ*pbp1b*::P$_c$-*erm* amplicon from E193) | Str$^R$ Erm$^R$ | This study |
| IU14820 IU14821 | D39 Δ*cps rpsL1 macP*(T32E) (IU14650 X fusion *macP* (T32E)) | Str$^R$ | This study |
| IU14822 IU14823 | D39 Δ*cps rpsL1 macP*(T32A) (IU14650 X fusion *macP* (T32A)) | Str$^R$ | This study |
| IU14966 | D39 Δ*cps* Δ*bgaA*::*tet*-P$_{Zn}$-RBS$^{ftsA}$-*macP* (IU14650 X fusion Δ*bgaA*::*tet*-P$_{Zn}$-RBS$^{ftsA}$-*macP*)) | Tet$^R$ | This study |
| IU14968 | D39 Δ*cps rpsL1* Δ*bgaA*::*tet*-P$_{Zn}$-RBS$^{ftsA}$-*macP* (IU1824 X fusion Δ*bgaA*::*tet*-P$_{Zn}$-RBS$^{ftsA}$-*macP*)) | Str$^R$ Tet$^R$ | This study |
| IU14972 | D39 Δ*cps* Δ*bgaA*::*kan*-t1t2-P$_{Zn}$-RBS$^{ftsA}$-*macP* (IU1945 X fusion Δ*bgaA*::*kan*-t1t2-P$_{Zn}$-RBS$^{ftsA}$-*macP*)) | Kan$^R$ | This study |
| IU14976 | D39 Δ*cps rpsL1* Δ*bgaA*::*kan*-t1t2-P$_{Zn}$-RBS$^{ftsA}$-*macP* (IU1824 X fusion Δ*bgaA*::*kan*-t1t2-P$_{Zn}$-RBS$^{ftsA}$-*macP*)) | Str$^R$ Kan$^R$ | This study |
| IU15084 | D39 Δ*cps rpsL1* Δ*macP*//Δ*bgaA*::*tet*-P$_{Zn}$-RBS$^{ftsA}$-*macP* (IU14650 X Δ*bgaA*::*tet*-P$_{Zn}$-RBS$^{ftsA}$-*macP* amplicon from IU14968)) | Str$^R$ Tet$^R$ | This study |
| IU15987 | D39 Δ*cps rpsL1* Δ*ireB* Δ*stkP*::P$_c$-*erm* (IU13604 X Δ*stkP*::P$_c$-*erm* amplicon from IU7923) | Str$^R$ Erm$^R$ | (Lamanna *et al.*, 2022) |
| IU16885 | D39 Δ*cps rpsL1 murZ*(D280Y) Δ*stkP*::P$_c$-*erm* | Str$^R$ Erm$^R$ | (Tsui *et al.*, 2023) |
| IU16616 | D39 Δ*cps rpsL1 macP*(T56E) (IU14650 X fusion *macP*(T56E)) | Str$^R$ | This study |
| IU16618 | D39 Δ*cps rpsL1 macP*(T56A) (IU14650 X fusion *macP*(T56A)) | Str$^R$ | This study |

| | | | |
|---|---|---|---|
| IU16722 | D39 Δ*cps rpsL1* Δ*macP* Δ*pbp2a*::P$_c$-*erm* (IU14650 X Δ*pbp2a*::P$_c$-*erm* amplicon from E180) | Str$^R$ Erm$^R$ | This study |
| IU16724 | D39 Δ*cps rpsL1 macP*(T32E) Δ*pbp1a*::P$_c$-*erm* (IU14820 X Δ*pbp1a*::P$_c$-*erm* amplicon from E177) | Str$^R$ Erm$^R$ | This study |
| IU16726 | D39 Δ*cps rpsL1 macP*(T32A) Δ*pbp1a*::P$_c$-*erm* (IU14822 X Δ*pbp1a*::P$_c$-*erm* amplicon from E177) | Str$^R$ Erm$^R$ | This study |
| IU16728 | D39 Δ*cps rpsL1 macP*(T32A) Δ*pbp2a*::P$_c$-*erm* (IU14820 X Δ*pbp2a*::P$_c$-*erm* amplicon from E180) | Str$^R$ Erm$^R$ | This study |
| IU16730 | D39 Δ*cps rpsL1 macP*(T56E) Δ*pbp1a*::P$_c$-*erm* (IU16615 X Δ*pbp1a*::P$_c$-*erm* amplicon from E177) | Str$^R$ Erm$^R$ | This study |
| IU16732 | D39 Δ*cps rpsL1 macP*(T56E) Δ*pbp2a*::P$_c$-*erm* (IU16615 X Δ*pbp2a*::P$_c$-*erm* amplicon from E180) | Str$^R$ Erm$^R$ | This study |
| IU16734 | D39 Δ*cps rpsL1 macP*(T56A) Δ*pbp1a*::P$_c$-*erm* (IU16617 X Δ*pbp1a*::P$_c$-*erm* amplicon from E177) | Str$^R$ Erm$^R$ | This study |
| IU16736 | D39 Δ*cps rpsL1 macP*(T56A) Δ*pbp2a*::P$_c$-*erm* (IU16617 X Δ*pbp2a*::P$_c$-*erm* amplicon from E180) | Str$^R$ Erm$^R$ | This study |
| IU16761 | D39 Δ*cps rpsL1* Δ*macP* markerless Δ*pbp2a*::P$_c$-*erm* (IU14650 X Δ*pbp2a*::P$_c$-*erm* amplicon from E180) | Str$^R$ Erm$^R$ | This study |
| IU16762 | D39 Δ*cps rpsL1* Δ*macP* markerless Δ*pbp1b*::P$_c$-*erm* (IU14699 X Δ*pbp1b*::P$_c$-*erm* amplicon from E193) | Str$^R$ Erm$^R$ | This study |
| IU16771 | D39 Δ*cps rpsL1 macP*(T32A) Δ*pbp1b*::P$_c$-*erm* (IU14823 X Δ*pbp1b*::P$_c$-*erm* amplicon from E193) | Str$^R$ Erm$^R$ | This study |
| IU16787 | D39 Δ*cps rpsL macP*(T32E T56E) (IU14650 X fusion *macP*(T32E T56E)) | Str$^R$ | This study |
| IU16788 | D39 Δ*cps rpsL macP*(T32A T56A) (IU14650 X fusion *macP*(T32A T56A)) | Str$^R$ | This study |
| IU16801 | D39 Δ*cps rpsL macP*(T32E T56E) Δ*pbp1a*::P$_c$-*erm* (IU16787 X Δ*pbp1a*::P$_c$-*erm* amplicon from E177) | Str$^R$ Erm$^R$ | This study |
| IU16805 | D39 Δ*cps rpsL macP*(T32A T56A) Δ*pbp1a*::P$_c$-*erm* (IU16788 X Δ*pbp1a*::P$_c$-*erm* amplicon from E177) | Str$^R$ Erm$^R$ | This study |
| IU16978 | D39 Δ*cps rpsL1* FLAG-*macP*(T32A) (IU14650 X fusion FLAG-*macP*(T32A)) | Str$^R$ | This study |
| IU16980 | D39 Δ*cps rpsL1* FLAG-*macP*(T56A) (IU14650 X fusion FLAG-*macP*(T56A)) | Str$^R$ | This study |
| IU17032 | D39 Δ*cps rpsL1* FLAG-*macP* (IU14650 X fusion FLAG-*macP*) | Str$^R$ | This study |
| IU17035 | D39 Δ*cps rpsL1* FLAG-*macP* (T32A T56A) (IU14650 X fusion FLAG-*macP*(T32A T56A)) | Str$^R$ | This study |
| IU17055 | D39 Δ*cps rpsL1* FLAG-*macP* Δ*pbp1a*::P$_c$-*erm* (IU17032 X Δ*pbp1a*::P$_c$-*erm* amplicon from E177) | Str$^R$ Erm$^R$ | This study |
| IU17187 | D39 Δ*cps rpsL1* Δ*macP*::P$_c$-*erm* (IU1824 X Δ*macP*::P$_c$-*erm* amplicon from E793) | Erm$^R$ | This study |
| IU17228 | D39 Δ*cps rpsL1 macP*(Δ5-8) (IU14650 X fusion *macP*(Δ5-8)) | Str$^R$ | This study |
| IU17232 | D39 Δ*cps rpsL1 macP*(Δ46-53) (IU14650 X fusion *macP*(Δ46-53)) | Str$^R$ | This study |

| | | | |
|---|---|---|---|
| IU17236 | D39 Δ*cps rpsL1 macP*(1-77) (IU14650 X fusion *macP*(1-77)) | Str$^R$ | This study |
| IU17240 | D39 Δ*cps rpsL1 macP*(1-87) (IU14650 X fusion *macP*(1-87)) | Str$^R$ | This study |
| IU17263 | D39 Δ*cps rpsL1* Δ*macP::*P$_c$-[*kan-rpsL*$^+$] (IU1824 X Δ*macP::*P$_c$-[*kan-rpsL*$^+$] amplicon from K793) | Kan$^R$ | This study |
| IU17266 | D39 Δ*cps rpsL1* Δ*pbp2a* markerless Δ*macP*::P$_c$-[*kan-rpsL*$^+$] (IU13256 X Δ*macP::*P$_c$-[*kan-rpsL*+] amplicon from K793) | Kan$^R$ | This study |
| IU17269 | D39 Δ*cps rpsL1* Δ*macP*//Δ*bgaA*::tet-P$_{Zn}$-RBS$^{ftsA}$-*macP* Δ*pbp1a*::P$_c$-*erm* (IU15084 X Δ*pbp1a*::P$_c$-*erm* amplicon from E177) | Str$^R$ Tet$^R$ Erm$^R$ | This study |
| IU17271 | D39 Δ*cps rpsL1 macP*(Δ5-8) Δ*pbp1a*::P$_c$-*erm* (IU17228 X Δ*pbp1a*::P$_c$-*erm* amplicon from E177) | Str$^R$ Erm$^R$ | This study |
| IU17273 | D39 Δ*cps rpsL1 macP*(Δ5-8) Δ*pbp2a*::P$_c$-*erm* (IU17228 X Δ*pbp2a*::P$_c$-*erm* amplicon from E180) | Str$^R$ Erm$^R$ | This study |
| IU17275 | D39 Δ*cps rpsL1 macP*(Δ46-53) Δ*pbp1a*::P$_c$-*erm* (IU17232 X Δ*pbp1a*::P$_c$-*erm* amplicon from E177) | Str$^R$ Erm$^R$ | This study |
| IU17277 | D39 Δ*cps rpsL1 macP*(Δ46-53) Δ*pbp2a*::P$_c$-*erm* (IU17232 X Δ*pbp2a*::P$_c$-*erm* amplicon from E180) | Str$^R$ Erm$^R$ | This study |
| IU17279 | D39 Δ*cps rpsL1 macP*(N77stop) Δ*pbp2a*::P$_c$-*erm* (IU17236 X Δ*pbp2a*::P$_c$-*erm* amplicon from E180) | Str$^R$ Erm$^R$ | This study |
| IU17281 | D39 Δ*cps rpsL1 macP*(L87stop) Δ*pbp2a*::P$_c$-*erm* (IU17240 X Δ*pbp2a*::P$_c$-*erm* amplicon from E180) | Str$^R$ Erm$^R$ | This study |
| IU17382 | D39 Δ*cps rpsL1 macP*$^+$-P$_c$-*erm* (IU14650 X fusion *macP*$^+$-P$_c$-*erm*) | Erm$^R$ | This study |
| IU17384 | D39 Δ*cps rpsL1 macP*(Δ30-33) (IU14650 X fusion *macP*(Δ30-33)) | Str$^R$ | This study |
| IU17386 | D39 Δ*cps rpsL1 macP*(Δ21-58) (IU14650 X fusion *macP*(Δ21-58)) | Str$^R$ | This study |
| IU17456 | D39 Δ*cps rpsL1 macP*(Δ30-33) Δ*pbp1a*::P$_c$-*erm* (IU17384 X Δ*pbp1a*::P$_c$-*erm* amplicon from E177) | Str$^R$ Erm$^R$ | This study |
| IU17458 | D39 Δ*cps rpsL1 macP*(Δ30-33) Δ*pbp2a*::P$_c$-*erm* (IU17384 X Δ*pbp2a*::P$_c$-*erm* amplicon from E180) | Str$^R$ Erm$^R$ | This study |
| IU17460 | D39 Δ*cps rpsL1 macP*(Δ21-58) Δ*pbp1a*::P$_c$-*erm* (IU17386 X Δ*pbp1a*::P$_c$-*erm* amplicon from E177) | Str$^R$ Erm$^R$ | This study |
| IU17462 | D39 Δ*cps rpsL1 macP*(Δ21-58) Δ*pbp2a*::P$_c$-*erm* (IU17386 X Δ*pbp2a*::P$_c$-*erm* amplicon from E180) | Str$^R$ Erm$^R$ | This study |
| IU18342<br>IU18358 | D39 Δ*cps rpsL1* FLAG-*macP*(T32E T56E) IU14650 X fusion FLAG-*macP*(T32E T56E)) | Str$^R$ | This study |
| IU18345 | D39 Δ*cps rpsL1* FLAG-*macP*(Δ46-53) (IU14650 X fusion FLAG-*macP*(Δ46-53)) | Str$^R$ | This study |
| IU18346 | D39 Δ*cps rpsL1* FLAG-*macP*(Δ30-33) (IU14650 X fusion FLAG-*macP*(Δ30-33)) | Str$^R$ | This study |
| IU18348 | D39 Δ*cps rpsL1* FLAG-*macP*(Δ21-58) (IU14650 X fusion FLAG-*macP*(Δ21-58)) | Str$^R$ | This study |
| IU18351 | D39 Δ*cps rpsL1* FLAG-*macP*(1-77) (IU14650 X fusion FLAG-*macP*(1-77)) | Str$^R$ | This study |
| IU18352 | D39 Δ*cps rpsL1* FLAG-*macP*(1-87) (IU14650 X fusion FLAG-*macP*(1-87)) | Str$^R$ | This study |

| | | | |
|---|---|---|---|
| IU18354 | D39 Δ*cps rpsL1* FLAG-*macP*(T32E) (IU14650 X fusion FLAG-*macP*(T32E)) | $Str^R$ | This study |
| IU18356 | D39 Δ*cps rpsL1* FLAG-*macP*(T56E) (IU14650 X fusion FLAG-*macP*(T56E)) | $Str^R$ | This study |
| IU18364 | D39 Δ*cps rpsL1* Δ*macP* Δ*pbp2a*::$P_c$-*erm* (IU14699 X Δ*pbp2a*::$P_c$-*erm* amplicon from E180) | $Str^R$ $Erm^R$ | This study |
| IU18366 | D39 Δ*cps rpsL1* Δ*macP* markerless Δ*pbp1b*::$P_c$-*erm* (IU14699 X Δ*pbp1b*::$P_c$-*erm* amplicon from E193) | $Str^R$ $Erm^R$ | This study |
| IU18368 | D39 Δ*cps rpsL1* FLAG-*macP*(Δ21-58) Δ*pbp1a*::$P_c$-*erm* (IU18348 X Δ*pbp1a*::$P_c$-*erm* amplicon from E177) | $Str^R$ $Erm^R$ | This study |
| IU18370 | D39 Δ*cps rpsL1* FLAG-*macP*(Δ21-58) Δ*pbp2a*::$P_c$-*erm* (IU18348 X Δ*pbp2a*::$P_c$-*erm* amplicon from E180) | $Str^R$ $Erm^R$ | This study |
| IU18372 | D39 Δ*cps rpsL1* FLAG-*macP*(Δ21-58) Δ*pbp1b*::$P_c$-*erm* (IU18348 X Δ*pbp1b*::$P_c$-*erm* amplicon from E193) | $Str^R$ $Erm^R$ | This study |
| IU18579 | D39 Δ*cps rpsL1* Δ*pbp1a* markerless | $Str^R$ | (Lamanna *et al.*, 2022) |
| IU18751 | D39 Δcps rpsL1 Δ*pbp1b* markerless Δ*rocS*::$P_c$-*erm* (IU14697 X fusion Δ*rocS*::$P_c$-*erm*) | $Str^R$ $Erm^R$ | This study |
| IU19214 | D39 Δ*cps rpsL1* Δ*macP*::$P_c$-*aad9* (IU14650 X fusion Δ*macP*::$P_c$-*aad9*) | $Spec^R$ | This study |
| IU19216 | D39 Δ*cps rpsL1* Δ*pbp1a*::$P_c$-*erm*// Δ*bgaA*::*tet*-$P_{Zn}$-RBS$^{ftsA}$-*pbp1a* Δ*pbp2a*::$P_c$-[*kan-rpsL*$^+$] (IU14357 X Δ*pbp2a*::$P_c$-[*kan-rpsL*$^+$] amplicon from K166) | $Erm^R$ $Kan^R$ $Tet^R$ | This study |
| IU19240 | D39 Δ*cps rpsL1* Δ*pbp1a*::$P_c$-*erm*// Δ*bgaA*::*tet*-$P_{Zn}$-RBS$^{ftsA}$-*pbp1a pbp2a*(A77T) (IU19216 X fusion *pbp2a*(A77T)) | $Erm^R$ $Str^R$ $Tet^R$ | This study |
| IU19242 | D39 Δ*cps rpsL1* Δ*pbp1a*::$P_c$-*erm*// Δ*bgaA*::*tet*-$P_{Zn}$-RBS$^{ftsA}$-*pbp1a pbp2a*(K78A) (IU19216 X fusion *pbp2a*(K78A)) | $Erm^R$ $Str^R$ $Tet^R$ | This study |
| IU19431 | D39 Δ*cps rpsL1* Δ*pbp1a*::$P_c$-*erm*// Δ*bgaA*::*tet*-$P_{Zn}$-RBS$^{ftsA}$-*pbp1a* Δ*macP*::$P_c$-*aad9* (IU14357 X Δ*macP*::$P_c$-*aad9* amplicon from IU19214) | $Erm^R$ $Spec^R$ $Tet^R$ | This study |
| IU19433 | D39 Δ*cps rpsL1* Δ*pbp1a*::$P_c$-*erm*// Δ*bgaA*::*tet*-$P_{Zn}$-RBS$^{ftsA}$-*pbp1a pbp2a*(A77T) Δ*macP*::$P_c$-*aad9* (IU19240 X Δ*macP*::$P_c$-*aad9* amplicon from IU19214) | $Erm^R$ $Spec^R$ $Tet^R$ $Str^R$ | This study |
| IU19435 | D39 Δ*cps rpsL1* Δ*pbp1a*::$P_c$-*erm*// Δ*bgaA*::*tet*-$P_{Zn}$-RBS$^{ftsA}$-*pbp1a pbp2a*(K78A) Δ*macP*::$P_c$-*aad9* (IU19242 X Δ*macP*::$P_c$-*aad9* amplicon from IU19214) | $Erm^R$ $Spec^R$ $Tet^R$ $Str^R$ | This study |
| IU19522 | D39 Δ*cps rpsL1 macP*(R76A N80A) (IU14650 X fusion *macP*(R76A N80A)) | $Str^R$ | This study |
| IU19524 | D39 Δ*cps rpsL1* FLAG-*macP*(R76A N80A) (IU14650 X fusion FLAG-*macP*(R76A N80A)) | $Str^R$ | This study |

| | | | |
|---|---|---|---|
| IU19526 | D39 Δ*cps rpsL1 macP*(L83A I86A L87A) (IU14650 X fusion *macP*(L83A I86A L87A)) | $Str^R$ | This study |
| IU19528 | D39 Δ*cps rpsL1* FLAG-*macP*(L83A I86A L87A) (IU14650 X fusion FLAG-*macP*(L83A I86A L87A)) | $Str^R$ | This study |
| IU19529 | D39 Δ*cps rpsL1* FLAG-*macP*(M101A L103A) (IU14650 X fusion FLAG-*macP*(M101A L103A)) | $Str^R$ | This study |
| IU19530 | D39 Δ*cps rpsL1* FLAG-*macP*(Δ5-8) (IU14650 X fusion F-*macP*(Δ5-8)) | $Str^R$ | This study |
| IU19533 | D39 Δ*cps rpsL1 pbp2a*(R51A H53A K56A) (IU7853 X fusion *pbp2a*(R51A H53A K56A)) | $Str^R$ | This study |
| IU19605 | D39 Δ*cps rpsL1 macP*(76A N80A) Δ*pbp1a*::$P_c$-*erm* (IU19522 X Δ*pbp1a*::$P_c$-*erm* amplicon from E177) | $Str^R$ $Erm^R$ | This study |
| IU19607 | D39 Δ*cps rpsL1 macP*(76A N80A) Δ*pbp2a*::$P_c$-*erm* (IU19522 X Δ*pbp2a*::$P_c$-*erm* amplicon from E180) | $Str^R$ $Erm^R$ | This study |
| IU19609 | D39 Δ*cps rpsL1* FLAG-*macP*(76A N80A) Δ*pbp1a*::$P_c$-*erm* (IU19524 X Δ*pbp1a*::$P_c$-*erm* amplicon from E177) | $Str^R$ $Erm^R$ | This study |
| IU19611 | D39 Δ*cps rpsL1* FLAG-*macP*(76A N80A) Δ*pbp2a*::$P_c$-*erm* (IU19524 X Δ*pbp2a*::$P_c$-*erm* amplicon from E180) | $Str^R$ $Erm^R$ | This study |
| IU19613 | D39 Δ*cps rpsL1 macP*(L83A I86A L87A) Δ*pbp2a*::$P_c$-*erm* (IU19526 X Δ*pbp2a*::$P_c$-*erm* amplicon from E180) | $Str^R$ $Erm^R$ | This study |
| IU19615 | D39 Δ*cps rpsL1* FLAG-*macP*(L83A I86A L87A) Δ*pbp2a*::$P_c$-*erm* (IU19528 X Δ*pbp2a*::$P_c$-*erm* amplicon from E180) | $Str^R$ $Erm^R$ | This study |
| IU19619 | D39 Δ*cps rpsL1* FLAG-*macP*(M101A L103A) Δ*pbp1a*::$P_c$-*erm* (IU19529 X Δ*pbp1a*::$P_c$-*erm* amplicon from E177) | $Str^R$ $Erm^R$ | This study |
| IU19621 | D39 Δ*cps rpsL1* FLAG-*macP*(M101A L103A) Δ*pbp2a*::$P_c$-*erm* (IU19529 X Δ*pbp2a*::$P_c$-*erm* amplicon from E180) | $Str^R$ $Erm^R$ | This study |
| IU19623 | D39 Δ*cps rpsL1* FLAG-*macP*(Δ5-8) Δ*pbp1a*::$P_c$-*erm* (IU19530 X Δ*pbp1a*::$P_c$-*erm* amplicon from E177) | $Str^R$ $Erm^R$ | This study |
| IU19625 | D39 Δ*cps rpsL1* FLAG-*macP*(Δ5-8) Δ*pbp2a*::$P_c$-*erm* (IU19530 X Δ*pbp2a*::$P_c$-*erm* amplicon from E180) | $Str^R$ $Erm^R$ | This study |
| IU19628 | D39 Δ*cps rpsL1 macP*(T56E) Δ*pbp1a*::$P_c$-*erm* (IU16616 X Δ*pbp1a*::$P_c$-*erm* amplicon from E177) | $Str^R$ $Erm^R$ | This study |
| IU19629 | D39 Δ*cps rpsL1 macP*(T56E) Δ*pbp2a*::$P_c$-*erm* (IU16616 X Δ*pbp2a*::$P_c$-*erm* amplicon from E180) | $Str^R$ $Erm^R$ | This study |
| IU19631 | D39 Δ*cps rpsL1 macP*(Δ30-33) Δ*pbp1a*::$P_c$-*erm* (IU17384 X Δ*pbp1a*::$P_c$-*erm* amplicon from E177) | $Str^R$ $Erm^R$ | This study |
| IU19633 | D39 Δ*cps rpsL1 macP*(Δ30-33) Δ*pbp2a*::$P_c$-*erm* (IU17384 X Δ*pbp2a*::$P_c$-*erm* amplicon from E180) | $Str^R$ $Erm^R$ | This study |

| | | | |
|---|---|---|---|
| IU19635 | D39 Δ*cps rpsL1 macP*(Δ21-58) Δ*pbp1a*::P$_c$-*erm* (IU17386 X Δ*pbp1a*::P$_c$-*erm* amplicon from E177) | Str$^R$ Erm$^R$ | This study |
| IU19637 | D39 Δ*cps rpsL1 macP*(Δ21-58) Δ*pbp2a*::P$_c$-*erm* (IU17386 X Δ*pbp2a*::P$_c$-*erm* amplicon from E180) | Str$^R$ Erm$^R$ | This study |
| IU19735 | D39 Δ*cps rpsL1* Δ*bgaA*::P$_c$-*aad9* (IU1824 X fusion Δ*bgaA*::P$_c$-*aad9*) | Spec$^R$ | This study |
| IU19908 | D39 Δ*cps rpsL1 murZ*(D280Y) Δ*stkP*::P$_c$-*erm* Δ*pbp1a*::P$_c$-[*kan-rpsL*$^+$] (IU16885 X Δ*pbp1a*::P$_c$-[*kan-rpsL*$^+$] amplicon from IU6726) | Erm$^R$ Kan$^R$ | This study |
| IU19947 | D39 Δ*cps rpsL1 gfp*-L-*macP* (IU14650 X fusion *gfp*-L-*macP*) | Str$^R$ | This study |
| IU20380 | D39 Δ*cps rpsL1 pbp2a*(A77T) (IU7853 X *pbp2a*(A77T) amplicon from IU19240) | Str$^R$ | This study |
| IU20382 | D39 Δ*cps rpsL1 pbp2a*(K78A) (IU7853 X *pbp2a*(K78A) amplicon from IU19242) | Str$^R$ | This study |
| IU21139 IU21140 | D39 Δ*cps rpsL1 macP*(S68A R69A R70A K75A R76A) (IU14650 X fusion *macP*(S68A R69A R70A K75A R76A)) | Str$^R$ | This study |
| IU21169 | D39 Δ*cps rpsL1 macP*(S68A R69A R70A K75A R76A) Δ*pbp1a*::P$_c$-*erm* (IU21139 X Δ*pbp1a*::P$_c$-*erm* amplicon from E177) | Str$^R$ Erm$^R$ | This study |
| IU21336 | D39 Δ*cps rpsL1 gfp*-L-*macP* Δ*pbp2a*::P$_c$-[*kan-rpsL*$^+$] (IU19947 X Δ*pbp2a*::P$_c$-[*kan-rpsL*$^+$] amplicon from K166) | Str$^R$ Kan$^R$ | This study |
| IU21339 | D39 Δ*cps rpsL1 gfp*-L-*macP*(R76A N80A) (IU14650 X fusion *gfp*-L-*macP*(R76A N80A)) | Str$^R$ | This study |
| IU21340 | D39 Δ*cps rpsL1 gfp*-L-*macP*(L83A I86A L87A) (IU14650 X fusion *gfp*-L-*macP*(R76A N80A)) | Str$^R$ | This study |
| IU21345 | D39 Δ*cps rpsL1 gfp*-L-*macP*(S68A R69A R70A K75A R76A) (IU14650 X fusion *gfp*-L-*macP*(S68A R69A R70A K75A R76A)) | Str$^R$ | This study |
| IU21376 IU21377 | D39 Δ*cps rpsL1* FLAG-*macP*(S68A R69A R70A K75A R76A) (IU14650 X fusion FLAG-*macP*(S68A R69A R70A K75A R76A)) | Str$^R$ | This study |
| Sp393 | D39 Δ*cps macP*::Sweet Janus cassette (*sacB-kan-rpsL*$^+$) (IU1945 X fusion amplicon *macP*::Sweet Janus cassette (*sacB-kan-rpsL*$^+$) | Kan$^R$ | This study |
| Sp480 | D39 Δ*cps* FLAG-L-*macP* (Sp393 X fusion amplicon FLAG-L-*macP*) | None | This study |
| Sp481 | D39 Δ*cps* FLAG-L-*macP*-(T32A T56A) (Sp393 X fusion amplicon FLAG-L-*macP*(T32A T56A)) | None | This study |
| Sp486 | D39 Δ*cps* FLAG-L-*macP*(T32A) (Sp393 X fusion amplicon FLAG-L-*macP*(T32A)) | None | This study |
| Sp487 | D39 Δ*cps* FLAG-L-*macP*(T56A) (Sp393 X fusion amplicon FLAG-L-*macP*-(T56A)) | None | This study |

| ***E. coli*** **strains** | | | |
|---|---|---|---|
| Strain name | Genotype/description | Antibiotic resistance[b] | Reference or source |

| | | | |
|---|---|---|---|
| DH5α | *F- Φ80lacZΔM15 Δ(lacZYA-argF) U169 recA1 endA1 hsdR17 (rk-, mk+) phoA supE44 λ- thi-1 gyrA96 relA1* | None | Invitrogen |
| BL21 | *F- ompT gal [dcm][lon] hsdSB (rB- mB-) (DE3)* | None | Novagen |
| BTH101 | *F-, cya-99, araD139, galE15, galK16, rpsL1 (Str R), hsdR2, mcrA1, mcrB1.* | Str$^R$ | Euromedex |

[a]Strains were constructed as described in *Experimental procedures.*

[b]Antibiotic resistance markers: Erm$^R$, erythromycin; Kan$^R$, kanamycin; Spec$^R$, spectinomycin; Str$^R$, streptomycin; Tet$^R$, tetracycline; Cm$^R$, chloramphenicol.

**Table S2.** Oligonucleotides used in this study

| **Primers used to construct strains** | | | |
|---|---|---|---|
| Primer | Sequence (5' to 3') | Template[a] | Amplicon Product |
| **For construction of E793 (Δ*macP*:: $P_c$-*erm*)** | | | |
| P1715 | TATCGGAAACTGTGAAGTGGGAAGCAACGT | D39 | upstream + 5’ 60 bp of *macP* |
| P1717 | CATTATCCATTAAAAATCAAACGGATCCTAAATTTTTTCGCCTCTATTAGCTCTTTCAAT | | |
| kan-rpsL forward | TAGGATCCGTTTGATTTTTAATGGATAATG | $P_c$-*erm* cassette[b] | $P_c$-*erm* |
| kan-rpsL reverse | GGGCCCCTTTCCTTATGCTTTTG | | |
| P1718 | AACGTCCAAAAGCATAAGGAAAGGGGCCCATCTTATTTGCGGTCATCTTTCTCTTGATTT | D39 | 3' 60 bp of *macP* + downstream |
| P1716 | CTTTGACACGGTTATTGATTGCTTGGCGAG | | |
| **For construction of K793 (Δ*macP*:: $P_c$-[*kan-rpsL*$^+$])** | | | |
| P1715 | TATCGGAAACTGTGAAGTGGGAAGCAACGT | D39 | upstream + 5’ 60 bp of *macP* |
| P1717 | CATTATCCATTAAAAATCAAACGGATCCTAAATTTTTTCGCCTCTATTAGCTCTTTCAAT | | |
| kan-rpsL forward | TAGGATCCGTTTGATTTTTAATGGATAATG | $P_c$-[*kan-rpsL*$^+$] cassette[b] | $P_c$-[*kan-rpsL*$^+$] |
| Kan-rpsL reverse | GGGCCCCTTTCCTTATGCTTTTG | | |
| P1718 | AACGTCCAAAAGCATAAGGAAAGGGGCCCATCTTATTTGCGGTCATCTTTCTCTTGATTT | D39 | 3' 60 bp of *macP* + downstream |
| P1716 | CTTTGACACGGTTATTGATTGCTTGGCGAG | | |
| **For construction of IU14151 (*ireB*-L-$FLAG^3$-$P_c$-*erm*)[c,d]** | | | |
| P1711 | GAGTGTCAATGAAGTTCTCAATCTGATTATGGAAACACC | D39 | upstream + 5' 60 bp of *ireB* |
| AJP360 | GAACCAGCAGCGGAGCCAGCGGAACCTAGATCGACTCCTTGTCCTTTGAGATAGTA | | |
| JQ179 | GGTTCCGCTGGCTCCGCTGCTGGTTCTGGC | IU4970 | L-$FLAG^3$-$P_c$-*erm* |
| JQ184 | TTATTTCCTCCCGTTAAATAATAGATAACTAT | | |
| AJP363 | GAACCAGCAGCGGAGCCAGCGGAACCTAGATCGACTCCTTGTCCTTTGAGATAGTA | D39 | downstream |
| P1712 | TGAACCTGAAATCCCCCTGTAACCAGAACT | | |
| **For construction of IU14699 (Δ*macP* markerless)** | | | |
| P1715 | TATCGGAAACTGTGAAGTGGGAAGCAACGT | D39 | upstream + 5’ 60 bp of *macP* |
| TT1103 | GAGAAAGATGACCGCAAATAAGATAATTTTTTCGCCTCTATTAGCTCTTTCAAT | | |

| | | | |
|---|---|---|---|
| TT1104 | AAGAGCTAATAGAGGCGAAAAAATTATCTTATTTGCGGTCATCTTTCTCTTGATTT | D39 | 60 bp of 3’ *macP* + downstream |
| P1716 | CTTTGACACGGTTATTGATTGCTTGGCGAG | | |
| **For construction of IU14820 and IU14821 (*macP*(T32E))** | | | |
| P1715 | TATCGGAAACTGTGAAGTGGGAAGCAACGT | D39 | upstream + 5’ *macP*(T32E) |
| TT1133 | GAGGTTGGTAAAATCTTTTCTTCCTCATTATCATCTAGCA | | |
| TT1134 | TGCTAGATGATAATGAGGAAGAAAAGATTTTACCAACCTC | D39 | 3’ *macP*(T32E) + downstream |
| P1716 | CTTTGACACGGTTATTGATTGCTTGGCGAG | | |
| **For construction of IU14822 and IU14823 (*macP*(T32A))** | | | |
| P1715 | TATCGGAAACTGTGAAGTGGGAAGCAACGT | D39 | upstream + 5’ *macP*(T32A) |
| TT1131 | AGGTTGGTAAAATCTTAGCTTCCTCATTATCATCTAGCA | | |
| TT1132 | TGCTAGATGATAATGAGGAAGCTAAGATTTTACCAACCT | D39 | 3’ *macP*(T32A) + downstream |
| P1716 | CTTTGACACGGTTATTGATTGCTTGGCGAG | | |
| **For construction of IU14966 and IU14968 (Δ*bgaA*::*tet*-$P_{Zn}$-RBS$^{ftsA}$-*macP*)** | | | |
| TT657 | CGCCCCAAGTTCATCACCAATGACATCAAC | IU10063 | *bgaA’ tet*-$P_{Zn}$-*RBS*$^{ftsA}$ |
| TT1135 | TCATCCGTTAATAAAGATTTACCCATTACATCGCTTCCTCTCTATCTTCCTTGT | | |
| TT1136 | GGAAGATAGAGAGGAAGCGATGTAATGGGTAAATCTTTATTAACGGATGAAATG | D39 | *macP*$^{+}$ |
| TT1137 | AGCAACTGGTTTATGAGAAAGTAAGTTCTTTTACAAAAGTTTCATTGCTAAAACAAGCAA | | |
| TT1138 | TGCTTGTTTTAGCAATGAAACTTTTGTAAAAGAACTTACTTTCTCATAAACCAGTTGCTG | D39 | *bgaA’* to downstream |
| CS121 | GCTTTCTTGAGGCAATTCACTTGGTGC | | |
| **For construction of IU14972, IU14976 and IU15084(Δ*bgaA*::*kan*-t1t2-$P_{Zn}$-RBS$^{ftsA}$-*macP)*** | | | |
| TT657 | CGCCCCAAGTTCATCACCAATGACATCAAC | IU12788 | 5’ Δ*bgaA*::*kan*-t1t2-$P_{Zn}$-RBS$^{ftsA}$- |
| TT1135 | TCATCCGTTAATAAAGATTTACCCATTACATCGCTTCCTCTCTATCTTCCTTGT | | |
| TT1136 | GGAAGATAGAGAGGAAGCGATGTAATGGGTAAATCTTTATTAACGGATGAAATG | D39 | *macP*$^{+}$ |
| TT1137 | AGCAACTGGTTTATGAGAAAGTAAGTTCTTTTACAAAAGTTTCATTGCTAAAACAAGCAA | | |
| TT1138 | TGCTTGTTTTAGCAATGAAACTTTTGTAAAAGAACTTACTTTCTCATAAACCAGTTGCTG | D39 | *bgaA’* to downstream |
| CS121 | GCTTTCTTGAGGCAATTCACTTGGTGC | | |
| **For construction of IU16616 (*macP*(T56E))** | | | |
| P1715 | TATCGGAAACTGTGAAGTGGGAAGCAACGT | D39 | upstream + 5’ *macP*(T56E) |
| MJ021 | TTCGACCTGAATCTTCAACTCTTCCTGGCTAAAACCATGAT | | |

| | | | |
|---|---|---|---|
| MJ020 | ATCATGGTTTTAGCCAGGAAGAGTTGAAGAT TCAGGTCGAA | D39 | 3’ *macP*(T56E) + downstream |
| P1716 | CTTTGACACGGTTATTGATTGCTTGGCGAG | | |
| **For construction of IU16618 (*macP*(T56A))** | | | |
| P1715 | TATCGGAAACTGTGAAGTGGGAAGCAACGT | D39 | upstream + 5’ *macP*(T56A) |
| MJ023 | TTCGACCTGAATCTTCAAGGCTTCCTGGCT AAAACCATGAT | | |
| MJ022 | ATCATGGTTTTAGCCAGGAAGCCTTGAAGAT TCAGGTCGAA | D39 | 3’ *macP*(T56A) + downstream |
| P1716 | CTTTGACACGGTTATTGATTGCTTGGCGAG | | |
| **For construction of IU16787 (*macP*(T32E T56E))** | | | |
| P1715 | TATCGGAAACTGTGAAGTGGGAAGCAACGT | IU14820 | upstream + 5’ *macP*(T32E T56E) |
| MJ021 | TTCGACCTGAATCTTCAACTCTTCCTGGCTA AAACCATGAT | | |
| MJ020 | ATCATGGTTTTAGCCAGGAAGAGTTGAAGAT TCAGGTCGAA | D39 | 3’ *macP*(T32E T56E) + downstream |
| P1716 | CTTTGACACGGTTATTGATTGCTTGGCGAG | | |
| **For construction of IU16788 (*macP*(T32A T56A))** | | | |
| P1715 | TATCGGAAACTGTGAAGTGGGAAGCAACGT | IU14822 | upstream + 5’ *macP*(T32A T56A) |
| MJ023 | TTCGACCTGAATCTTCAAGGCTTCCTGGCT AAAACCATGAT | | |
| MJ022 | ATCATGGTTTTAGCCAGGAAGCCTTGAAGAT TCAGGTCGAA | D39 | 3’ *macP*(T32A T56A) + downstream |
| P1716 | CTTTGACACGGTTATTGATTGCTTGGCGAG | | |
| **For construction of IU16978 (FLAG-*macP*(T32A))** | | | |
| P1715 | TATCGGAAACTGTGAAGTGGGAAGCAACGT | D39 | upstream + FLAG |
| MJ050 | TACCTTTATCATCATCATCTTTATAATCCATAC TGACCTCCTCTATTTTTTCTGCAA | | |
| MJ051 | TGGATTATAAAGATGATGATGATAAAGGTAAA TCTTTATTAACGGATGAAATGATTG | IU14822 | FLAG-*macP*(T32A) + downstream |
| P1716 | CTTTGACACGGTTATTGATTGCTTGGCGAG | | |
| **For construction of IU16980 (FLAG-*macP*(T56A))** | | | |
| P1715 | TATCGGAAACTGTGAAGTGGGAAGCAACGT | D39 | upstream + FLAG |
| MJ050 | TACCTTTATCATCATCATCTTTATAATCCATAC TGACCTCCTCTATTTTTTCTGCAA | | |
| MJ051 | TGGATTATAAAGATGATGATGATAAAGGTAAA TCTTTATTAACGGATGAAATGATTG | IU16618 | FLAG-*macP*(T56A) + downstream |
| P1716 | CTTTGACACGGTTATTGATTGCTTGGCGAG | | |
| **For construction of IU17032 (FLAG-*macP*)[d]** | | | |
| P1715 | TATCGGAAACTGTGAAGTGGGAAGCAACGT | D39 | upstream + FLAG |
| MJ050 | TACCTTTATCATCATCATCTTTATAATCCATAC TGACCTCCTCTATTTTTTCTGCAA | | |

| | | | |
|---|---|---|---|
| MJ051 | TGGATTATAAAGATGATGATGATAAAGGTAAA TCTTTATTAACGGATGAAATGATTG | D39 | FLAG-*macP* + downstream |
| P1716 | CTTTGACACGGTTATTGATTGCTTGGCGAG | | |
| **For construction of IU17035 (FLAG-*macP*(T32A T56A))** | | | |
| P1715 | TATCGGAAACTGTGAAGTGGGAAGCAACGT | IU17032 | upstream + 5' FLAG-*macP*(T32A T56A) |
| MJ023 | TTCGACCTGAATCTTCAAGGCTTCCTGGCT AAAACCATGAT | | |
| MJ022 | ATCATGGTTTTAGCCAGGAAGCCTTGAAGAT TCAGGTCGAA | IU16788 | 3' FLAG-*macP*(T32A T56A) + downstream |
| P1716 | CTTTGACACGGTTATTGATTGCTTGGCGAG | | |
| **For construction of IU17228 (*macP*(Δ5-8))** | | | |
| P1715 | TATCGGAAACTGTGAAGTGGGAAGCAACGT | D39 | upstream + 5' *macP*(Δ5-8) |
| TT1328 | GCCTCTATTAGCTCTTTCAATCATTTCAGATT TACCCATACTGACCTCCTCTATTTTT | | |
| TT1329 | AAAAATAGAGGAGGTCAGTATGGGTAAATCT GAAATGATTGAAAGAGCTAATAGAGGC | D39 | 3' *macP* *macP*(Δ5-8) + downstream |
| P1716 | CTTTGACACGGTTATTGATTGCTTGGCGAG | | |
| **For construction of IU17232 (*macP*(Δ46-53))** | | | |
| P1715 | TATCGGAAACTGTGAAGTGGGAAGCAACGT | D39 | upstream + 5' *macP*(Δ46-53) |
| TT1330 | CCTGAATCTTCAAGGTTTCCTGGGCATAAC CAAAACGGGAAGAAGA | | |
| TT1331 | TTCCCGTTTTGGTTATGCCCAGGAAACCTT GAAGATTCAGGTCG | D39 | 3' *macP* *macP*(Δ46-53) + downstream |
| P1716 | CTTTGACACGGTTATTGATTGCTTGGCGAG | | |
| **For construction of IU17236 (*macP*(1-N77))** | | | |
| P1715 | TATCGGAAACTGTGAAGTGGGAAGCAACGT | D39 | upstream + 5' *macP*(1-N77) |
| TT1332 | TTATTCAACTTAGAATTGAAGACTTAATTTCT CTTGGTATTTTCAATACGACGG | | |
| TT1333 | AAATACCAAGAGAAATTAAGTCTTCAATTCTA AGTTGAATAAAATCTTATTTGC | D39 | 3' *macP*(1-N77) + downstream |
| P1716 | CTTTGACACGGTTATTGATTGCTTGGCGAG | | |
| **For construction of IU17240 (*macP*(1-L87))** | | | |
| P1715 | TATCGGAAACTGTGAAGTGGGAAGCAACGT | D39 | upstream + 5' *macP*(1-L87) |
| TT1334 | GAGAAAGATGACCGCAAATTATAAGATTTTAT TCAACTTAGAATTGAAGACATTTCT | | |
| TT1335 | TAAGTTGAATAAAATCTTATAATTTGCGGTCA TCTTTCTCTTGATTTT | D39 | 3' *macP*(1-L87) + downstream |
| P1716 | CTTTGACACGGTTATTGATTGCTTGGCGAG | | |
| **For construction of IU17382 (*macP*$^{+}$-P$_{c}$-*erm*)** | | | |
| P1715 | TATCGGAAACTGTGAAGTGGGAAGCAACGT | D39 | upstream + *macP* |
| TT1337 | CATTATCCATTAAAAATCAAACGGATCCTATT CAATTCCTTTTCTATTACAAAAGTTTCA | | |

| | | | |
|---|---|---|---|
| kan-rpsL forward | TAGGATCCGTTTGATTTTTAATGGATAATG | $P_c$-*erm* cassette[b] | $P_c$-*erm* |
| kan-rpsL reverse | GGGCCCCTTTCCTTATGCTTTTG | | |
| P1718 | ATCTTATTTGCGGTCATCTTTCTCTTGATTT | D39 | downstream |
| P1716 | CTTTGACACGGTTATTGATTGCTTGGCGAG | | |
| **For construction of IU17384 (*macP*(Δ30-33))** | | | |
| P1715 | TATCGGAAACTGTGAAGTGGGAAGCAACGT | D39 | upstream + 5' *macP*(Δ30-33) |
| TT1338 | ACCAAAACGGGAAGAAGAGGTTGGTAAAAT ATTATCATCTAGCAAAGGAGGACCTGAAAT | | |
| TT1339 | ATTTCAGGTCCTCCTTTGCTAGATGATAATAT TTTACCAACCTCTTCTTCCCGTTTTGGT | D39 | 3' *macP*(Δ30-33)) + downstream |
| P1716 | CTTTGACACGGTTATTGATTGCTTGGCGAG | | |
| **For construction of IU17386 (*macP*(Δ21-58))** | | | |
| P1715 | TATCGGAAACTGTGAAGTGGGAAGCAACGT | D39 | upstream + 5' *macP*(Δ21-58) |
| TT1340 | GCTTTTATGAATAGATGGTTCGACCTGAATA ATTTTTTCGCCTCTATTAGCTCTTTCAAT | | |
| TT1341 | ATTGAAAGAGCTAATAGAGGCGAAAAAATTA TTCAGGTCGAACCATCTATTCATAAAAGC | D39 | 3' *macP*(Δ21-58)) + downstream |
| P1716 | CTTTGACACGGTTATTGATTGCTTGGCGAG | | |
| P1716 | CTTTGACACGGTTATTGTTGCTTGGCGAG | | |
| **For construction of IU18342 (FLAG-*macP*(T32E T56E))** | | | |
| P1715 | TATCGGAAACTGTGAAGTGGGAAGCAACGT | IU17032 | upstream + 5' FLAG-*macP*(T32E T56E) |
| MJ021 | TTCGACCTGAATCTTCAACTCTTCCTGGCTA AAACCATGAT | | |
| MJ020 | ATCATGGTTTTAGCCAGGAAGAGTTGAAGAT TCAGGTCGAA | IU16787 | 3' FLAG-*macP*(T32E T56E) + downstream |
| P1716 | CTTTGACACGGTTATTGATTGCTTGGCGAG | | |
| **For construction of IU18345 (FLAG-*macP*(Δ46-53))** | | | |
| P1715 | TATCGGAAACTGTGAAGTGGGAAGCAACGT | IU17032 | upstream + 5' FLAG- *macP*(Δ46-53) |
| TT1330 | CCTGAATCTTCAAGGTTTCCTGGGCATAAC CAAAACGGGAAGAAGA | | |
| TT1331 | TTCCCGTTTTGGTTATGCCCAGGAAACCTT GAAGATTCAGGTCG | IU17232 | 3' FLAG-*macP*(Δ46-53) + downstream |
| P1716 | CTTTGACACGGTTATTGATTGCTTGGCGAG | | |
| **For construction of IU18346 (FLAG-*macP*(Δ30-33))** | | | |
| P1715 | TATCGGAAACTGTGAAGTGGGAAGCAACGT | IU17032 | upstream + 5' FLAG-*macP*(Δ30-33) |
| TT1338 | ACCAAAACGGGAAGAAGAGGTTGGTAAAAT ATTATCATCTAGCAAAGGAGGACCTGAAAT | | |
| TT1339 | ATTTCAGGTCCTCCTTTGCTAGATGATAATAT TTTACCAACCTCTTCTTCCCGTTTTGGT | IU17384 | 3' F-*macP*(Δ30-33) + downstream of *macP* |
| P1716 | CTTTGACACGGTTATTGATTGCTTGGCGAG | | |

| | | | |
|---|---|---|---|
| **For construction of IU18348 (FLAG- *macP*(Δ21-58))** | | | |
| P1715 | TATCGGAAACTGTGAAGTGGGAAGCAACGT | IU17032 | upstream + 5' FLAG-*macP*(Δ21-58) |
| TT1340 | GCTTTTATGAATAGATGGTTCGACCTGAATAATTTTTTCGCCTCTATTAGCTCTTTCAAT | | |
| TT1341 | ATTGAAAGAGCTAATAGAGGCGAAAAAATTATTCAGGTCGAACCATCTATTCATAAAAGC | IU17386 | 3' FLAG-*macP*(Δ21-58) + downstream of *macP* |
| P1716 | CTTTGACACGGTTATTGATTGCTTGGCGAG | | |
| **For construction of IU18351 (FLAG-*macP*(1-N77))** | | | |
| P1715 | TATCGGAAACTGTGAAGTGGGAAGCAACGT | IU17032 | upstream + 5' FLAG-*macP*(1-N77) |
| TT1332 | TTATTCAACTTAGAATTGAAGACTTAATTTCTCTTGGTATTTTCAATACGACGG | | |
| TT1333 | AAATACCAAGAGAAATTAAGTCTTCAATTCTAAGTTGAATAAAATCTTATTTGC | IU17240 | 3' FLAG-*macP*(1-N77) + downstream of *macP* |
| P1716 | CTTTGACACGGTTATTGATTGCTTGGCGAG | | |
| **For construction of IU18352 (FLAG-*macP*(1-L87))** | | | |
| P1715 | TATCGGAAACTGTGAAGTGGGAAGCAACGT | IU17032 | upstream + 5' FLAG-*macP*(1-L87) |
| TT1334 | GAGAAAGATGACCGCAAATTATAAGATTTTATTCAACTTAGAATTGAAGACATTTCT | | |
| TT1335 | TAAGTTGAATAAAATCTTATAATTTGCGGTCATCTTTCTCTTGATTTT | IU17240 | 3' FLAG-*macP*(1-L87) + downstream of *macP* |
| P1716 | CTTTGACACGGTTATTGATTGCTTGGCGAG | | |
| **For construction of IU18354 (FLAG- *macP*(T32E))** | | | |
| P1715 | TATCGGAAACTGTGAAGTGGGAAGCAACGT | IU17032 | upstream + 5' FLAG-*macP*(T32E) |
| TT1133 | GAGGTTGGTAAAATCTTTTCTTCCTCATTATCATCTAGCA | | |
| TT1134 | TGCTAGATGATAATGAGGAAGAAAAGATTTTACCAACCTC | IU14820 | 3' *macP*(T32E) + downstream |
| P1716 | CTTTGACACGGTTATTGATTGCTTGGCGAG | | |
| **For construction of IU18356 (FLAG- *macP*(T56E))** | | | |
| P1715 | TATCGGAAACTGTGAAGTGGGAAGCAACGT | IU17032 | upstream + 5' FLAG-*macP*(T56E) |
| MJ021 | TTCGACCTGAATCTTCAACTCTTCCTGGCTAAAACCATGAT | | |
| MJ020 | ATCATGGTTTTAGCCAGGAAGAGTTGAAGATTCAGGTCGAA | IU16616 | 3' *macP*(T56E) + downstream |
| P1716 | CTTTGACACGGTTATTGATTGCTTGGCGAG | | |
| **For construction of IU18751 (Δ*rocS*::$P_c$-*erm*)** | | | |
| P1737 | ACCAACCTCTTCTTCCCGTTTTGGTTATGC | D39 | upstream + 5' 60 bp of *rocS* |
| P1738 | CATTATCCATTAAAAATCAAACGGATCCTATGCTTGGCGAGATAATCCTAAGACCTCTGC | | |

| | | | |
|---|---|---|---|
| kan-rpsL forward | TAGGATCCGTTTGATTTTTAATGGATAATG | $P_c$-*erm* cassette[b] | $P_c$-*erm* |
| kan-rpsL reverse | GGGCCCCTTTCCTTATGCTTTTG | | |
| P1739 | TAAACGTCCAAAAGCATAAGGAAAGGGGCCCGCAAAAGAAGAAGTCCAATCCACTAAAAA | D39 | 3' 60 bp of *rocS* + downstream |
| P1740 | GCAACCAGAAGGTTAATTACAATCAAGGCT | | |
| **For construction of IU19214 (Δ*macP*::$P_c$-*aad9*)** | | | |
| P1715 | TATCGGAAACTGTGAAGTGGGAAGCAACGT | D39 | upstream + 5' 60 bp of *macP* |
| P1717 | CATTATCCATTAAAAATCAAACGGATCCTAAATTTTTTCGCCTCTATTAGCTCTTTCAAT | | |
| kan-rpsL forward | TAGGATCCGTTTGATTTTTAATGGATAATG | $P_c$-*aad9* cassette[b] | $P_c$-*aad9* |
| kan-rpsL reverse | GGGCCCCTTTCCTTATGCTTTTG | | |
| P1718 | AACGTCCAAAAGCATAAGGAAAGGGGCCCATCTTATTTGCGGTCATCTTTCTCTTGATTT | D39 | 3' 60 bp of *macP* + downstream |
| P1716 | CTTTGACACGGTTATTGATTGCTTGGCGAG | | |
| **For construction of IU19240 (*pbp2a*(A77T))** | | | |
| P226 | GGTACGACAACGAAATGTCATACACTGCAC | D39 | upstream + 5' *pbp2a*(A77T) |
| MJ119 | TCATTGACATTGGTCGACTTTGTTACAGCAAACAAATAGATTC | | |
| MJ118 | GAATCTATTTGTTTGCTGTAACAAAGTCGACCAATGTCAATGA | D39 | 3' *pbp2a*(A77T) + downstream |
| P227 | TCTGTTCCCGTGTGATCCGACAAATCCT | | |
| **For construction of IU19242 (*pbp2a*(K78A))** | | | |
| P226 | GGTACGACAACGAAATGTCATACACTGCAC | D39 | upstream + 5' *pbp2a*(K78A) |
| MJ121 | AAATCATTGACATTGGTCGAAGCGGCTACAGCAAACAAATAGA | | |
| MJ120 | TCTATTTGTTTGCTGTAGCCGCTTCGACCAATGTCAATGATTT | D39 | 3' *pbp2a*(K78A) + downstream |
| P227 | TCTGTTCCCGTGTGATCCGACAAATCCT | | |
| **For construction of IU19522 (*macP*(R76A N80A))** | | | |
| P1715 | TATCGGAAACTGTGAAGTGGGAAGCAACGT | D39 | upstream + 5' *macP*(R76A N80A) |
| MJ143 | ATTGAAAATACCAAGGCTAATGTCTTCGCTTCTAAGTTG | | |
| MJ144 | CAACTTAGAAGCGAAGACATTAGCCTTGGTATTTTCAAT | D39 | 3' *macP*(R76A N80A) + downstream |
| P1716 | CTTTGACACGGTTATTGATTGCTTGGCGAG | | |
| **For construction of IU19524 (FLAG-*macP*(R76A N80A))** | | | |
| P1715 | TATCGGAAACTGTGAAGTGGGAAGCAACGT | IU17032 | upstream + 5' FLAG- |
| MJ143 | ATTGAAAATACCAAGGCTAATGTCTTCGCTTCTAAGTTG | | |

| | | | |
|---|---|---|---|
| | | | *macP*(R76A N80A) |
| MJ144 | CAACTTAGAAGCGAAGACATTAGCCTTGGTATTTTCAAT | D39 | 3' *macP*(R76A N80A) + downstream |
| P1716 | CTTTGACACGGTTATTGATTGCTTGGCGAG | | |
| **For construction of IU19526 (*macP*(L83A I86A L87A))** | | | |
| P1715 | TATCGGAAACTGTGAAGTGGGAAGCAACGT | D39 | upstream + 5' *macP*(L83A I86A L87A) |
| MJ158 | CTTCAATTCTAAGGCTAATAAAGCTGCTTTTGCGGTCATC | | |
| MJ159 | GATGACCGCAAAAGCAGCTTTATTAGCCTTAGAATTGAAG | D39 | 3' *macP*(L83A I86A L87A) + downstream |
| P1716 | CTTTGACACGGTTATTGATTGCTTGGCGAG | | |
| **For construction of IU19528 (FLAG-*macP*(L83A I86A L87A))** | | | |
| P1715 | TATCGGAAACTGTGAAGTGGGAAGCAACGT | IU17032 | upstream + 5' FLAG- *macP*(L83A I86A L87A) |
| MJ158 | CTTCAATTCTAAGGCTAATAAAGCTGCTTTTGCGGTCATC | | |
| MJ159 | GATGACCGCAAAAGCAGCTTTATTAGCCTTAGAATTGAAG | D39 | 3' *macP*(L83A I86A L87A) + downstream |
| P1716 | CTTTGACACGGTTATTGATTGCTTGGCGAG | | |
| **For construction of IU19529 (FLAG-*macP*(M101A L103A))** | | | |
| P1715 | TATCGGAAACTGTGAAGTGGGAAGCAACGT | IU17032 | upstream + 5' FLAG- *macP*(M101A L103A) |
| MJ147 | GTTTTAGCAGCTAAAGCTTTGTAATAGAAAAGGAATTGAA | | |
| MJ148 | TTCAATTCCTTTTCTATTACAAAGCTTTAGCTGCTAAAAC | D39 | 3' FLAG- *macP*(M101A L103A) + downstream |
| P1716 | CTTTGACACGGTTATTGATTGCTTGGCGAG | | |
| **For construction of IU19530 (FLAG-*macP*(Δ5-8))** | | | |
| P1715 | TATCGGAAACTGTGAAGTGGGAAGCAACGT | IU17032 | upstream + 5' FLAG-*macP*(Δ5-8) |
| MJ160 | GGTCAGTATGGATTATAAAGATGATGATGATAAAGGTAAATCTGAAATGATTGAAAGAGC | | |
| MJ161 | AGATTTACCTTTATCATCATCATCTTTATAATCCATACTGACC | IU17228 | 3' *macP*(Δ5-8) + downstream |
| P1716 | CTTTGACACGGTTATTGATTGCTTGGCGAG | | |
| **For construction of IU19533 (*pbp2a*(R51A H53A K56A))** | | | |
| P226 | GGTACGACAACGAAATGTCATACACTGCAC | D39 | upstream + 5' *pbp2a*(R51A H53A K56A) |
| MJ135 | CTATATTTAGCGCTTAATTCAGCGGCAAGAGCAAATTCTTT | | |
| MJ134 | AAAGAATTTGCTCTTGCCGCTGAATTAAGCGCTAAATATAG | D39 | 3' *pbp2a*(R51A H53A K56A) + downstream |
| P227 | TCTGTTCCCGTGTGATCCGACAAATCCT | | |
| **For construction of IU19735 (Δ*bgaA*::$P_c$-*aad9*)** | | | |

| | | | |
|---|---|---|---|
| P146 | TGGCCATTCATCGCTGGTCGTGCTGAAAT | D39 | upstream + 5' 60 bp of *bgaA* |
| P148 | CATTATCCATTAAAAATCAAACGGATCCTATCCCACAGCAAACTTACGAATGCTATAAAC | | |
| kan-rpsL forward | TAGGATCCGTTTGATTTTTAATGGATAATG | $P_c$-*aad9* cassette[b] | $P_c$-*aad9* |
| kan-rpsL reverse | GGGCCCCTTTCCTTATGCTTTTG | | |
| P149 | CAAAAGCATAAGGAAAGGGGCCCGCTCTTCTAGGTTTGAGTGCAGGATTAG | D39 | 3' 60 bp of *bgaA* + downstream |
| P147 | TACGCCTTCTATCATGCCTTTGATCGCCCGT | | |
| **For construction of IU19947 (*gfp*-L-*macP*)** | | | |
| P1715 | TATCGGAAACTGTGAAGTGGGAAGCAACGT | D39 | upstream + 5' *gfp* |
| MJ170 | CTGTAAACAATTCTTCACCTTTAGAAATCATACTGACCTCCTCTATTTTTTCTGCAA | | |
| MJ171 | AGTTGCAGAAAAAATAGAGGAGGTCAGTATGATTTCTAAAGGTGAAGAATTGTTTACAGG | IU9182 | *gfp-L* |
| TT757 | TCCGGATCCCTCGAGTTTATACAATTCATCC | | |
| MJ172 | ATGAATTGTATAAACTCGAGGGATCCGGAGGTAAATCTTTATTAACGGATGAAATGATTG | D39 | L + *macP* + *downstream* |
| P1716 | CTTTGACACGGTTATTGATTGCTTGGCGAG | | |
| **For construction of IU21139 (*macP*(S68A R69A R70A K75A R76A))** | | | |
| P1715 | TATCGGAAACTGTGAAGTGGGAAGCAACGT | D39 | upstream + 5' *macP*(S68A R69A R70A K75A R76A) |
| MJ191 | TTAGAATTGAAGACATTAGCAGCGGTATTTTCAATAGCAGCAGCTTTATGAATAGATGG | | |
| MJ190 | CCATCTATTCATAAAGCTGCTGCTATTGAAAATACCGCTGCTAATGTCTTCAATTCTAA | D39 | 3' *macP*(S68A R69A R70A K75A R76A) + downstream |
| P1716 | CTTTGACACGGTTATTGATTGCTTGGCGAG | | |
| **For construction of IU21339 (*gfp*-L-*macP*(R76A N80A))** | | | |
| P1715 | TATCGGAAACTGTGAAGTGGGAAGCAACGT | IU19947 | upstream + 5' *gfp*-L-*macP* |
| MJ144 | CAACTTAGAAGCGAAGACATTAGCCTTGGTATTTTCAAT | | |
| MJ143 | ATTGAAAATACCAAGGCTAATGTCTTCGCTTCTAAGTTG | IU19522 | *macP*(R76A N80A) + downstream |
| P1716 | CTTTGACACGGTTATTGATTGCTTGGCGAG | | |
| **For construction of IU21340 (*gfp*-L-*macP*(L83A I86A L87A))** | | | |
| P1715 | TATCGGAAACTGTGAAGTGGGAAGCAACGT | IU19947 | upstream + 5' *gfp*-L-*macP* |
| MJ159 | GATGACCGCAAAAGCAGCTTTATTAGCCTTAGAATTGAAG | | |
| MJ158 | CTTCAATTCTAAGGCTAATAAAGCTGCTTTTGCGGTCATC | IU19526 | *macP*(L83A I86A L87A) + downstream |
| P1716 | CTTTGACACGGTTATTGATTGCTTGGCGAG | | |
| **For construction of IU21345 (*gfp*-L-*macP*(S68A R69A R70A K75A R76A))** | | | |

| | | | |
|---|---|---|---|
| P1715 | TATCGGAAACTGTGAAGTGGGAAGCAACGT | IU19947 | upstream + 5' *gfp*-L-*macP* |
| MJ191 | TTAGAATTGAAGACATTAGCAGCGGTATTTT CAATAGCAGCAGCTTTATGAATAGATGG | | |
| MJ190 | CCATCTATTCATAAAGCTGCTGCTATTGAAAA TACCGCTGCTAATGTCTTCAATTCTAA | IU21140 | *macP*(S68A R69A R70A K75A R76A) + downstream |
| P1716 | CTTTGACACGGTTATTGATTGCTTGGCGAG | | |
| **For construction of IU21376 (FLAG-*macP*(S68A R69A R70A K75A R76A))** | | | |
| P1715 | TATCGGAAACTGTGAAGTGGGAAGCAACGT | IU17032 | upstream + 5' FLAG-*macP*(S68A R69A R70A K75A R76A) |
| MJ191 | TTAGAATTGAAGACATTAGCAGCGGTATTTT CAATAGCAGCAGCTTTATGAATAGATGG | | |
| MJ190 | CCATCTATTCATAAAGCTGCTGCTATTGAAAA TACCGCTGCTAATGTCTTCAATTCTAA | IU21139 | 3' *macP*(S68A R69A R70A K75A R76A) + downstream |
| P1716 | CTTTGACACGGTTATTGATTGCTTGGCGAG | | |
| **For construction of Sp393 (*macP::SWJ*)** | | | |
| DP1 | TTAAGGATCGATCCGTTTGATT | Sweet Janus cassette (*sacB-kan-rpsL*$^+$) | Sweet Janus cassette (*sacB-kan-rpsL*$^+$) |
| DP2 | TTATGCTTTTGGACGTTTAGTA | | |
| AU135 | CAAGTTCATATTGGTAACTTTG | D39 | *macP* upstream flanking sequence; fusion to Sweet Janus cassette to generate *macP*::Sweet Janus cassette deletion construct |
| AU136 | GAAAAAATAGAGGAGGTCAGTATTAAGGATC GATCCGTTTGATT | | |
| AU137 | TACTAAACGTCCAAAAGCATAATAGAAAAGG AATTGAAATGAAAAT | D39 | *macP* downstream flanking sequence; fusion to Sweet Janus cassette to generate *macP*::Sweet Janus cassette deletion construct |
| AU138 | GTTGTTACCAGAAGTGGCTT | | |
| **For construction of Sp480/Sp481/Sp486/Sp487 (FLAG-L-*macP*-WT/2TA/T32A/T56A)** | | | |
| LN377 | GATTACAAGGATGACGACGATAAGGGTTCC GCTGGCTCCGCTGCTGGTTCTGGCGGTAA ATCTTTATTAACGGAT | pKT25_*spd_0876-wt/mut* | *flag-L-macP-WT or mut* |
| LN333 | TTACAAAAGTTTCATTGCTAAAAC | | |
| AU135 | CAAGTTCATATTGGTAACTTTG | D39 | |

| | | | |
|---|---|---|---|
| LN378 | CTTATCGTCGTCATCCTTGTAATCCATACTGACCTCCTCTATTTT | | *macP* upstream flanking sequence; fusion to *flag-L-macP* to generate construct for allelic exchange |
| LN334 | GTTTTAGCAATGAAACTTTTGTAA | D39 | *macP* downstream flanking sequence; fusion to *flag-L-macP* to generate construct for allelic exchange |
| AU138 | GTTGTTACCAGAAGTGGCTT | | |
| **For construction of pETphos-*macP*ΔTM-T32A (mutagenesis)** | | | |
| LN304 | AGAAGAGGTTGGTAAAATCTTAGCTTCCTCATTATCATCTAGCAAAG | pETphos-*macP* | pETPhos-*macP*ΔTM -T32A |
| LN305 | CTTTGCTAGATGATAATGAGGAAGCTAAGATTTTACCAACCTCTTCT | | |
| **For construction of pETphos-*macP*-ΔTM -T56A (mutagenesis)** | | | |
| LN306 | GACCTGAATCTTCAAGGCTTCCTGGCTAAAACCATGAT | pETphos-*macP* | pETPhos-*macP*ΔTM -T56A |
| LN307 | ATCATGGTTTTAGCCAGGAAGCCTTGAAGATTCAGGTC | | |
| **For construction of pETphos-*macP*ΔTM -T32A T56A (2TA) (mutagenesis)** | | | |
| LN304 | AGAAGAGGTTGGTAAAATCTTAGCTTCCTCATTATCATCTAGCAAAG | pETPhos-*macP*-T56A | pETPhos-*macP*ΔTM T32A/T56A (=2TA) |
| LN305 | CTTTGCTAGATGATAATGAGGAAGCTAAGATTTTACCAACCTCTTCT | | |
| **For transformation assays** | | | |
| P234 | CCCTTGTGTTCATAGCGAGGATAAGCA | E177 | Δ*pbp1a*::$P_c$-*erm* |
| P235 | AGGCAAGCCTGCAACCATGGTCTTGAAA | | |
| P226 | GGTACGACAACGAAATGTCATACACTGCAC | E180 | Δ*pbp2a*::$P_c$-*erm* |
| P227 | TCTGTTCCCGTGTGATCCGACAAATCCT | | |
| P222 | CGTTCGTGTGGCGCTGCTTCAAATTGTT | E193 | Δ*pbp1b*::$P_c$-*erm* |
| P522 | AACGGCAACCACCAAAGGAGAAACCAAGGA | | |
| P1485 | CCAAGCCTTGTTGGAGGCGAATAATTCCCT | E736 | Δ*phpP*::$P_c$-*erm* |
| TT574 | CGCCTGCTCTGGTGACAAGTAATGAACTGA | | |
| P146 | TGGCCATTCATCGCTGGTCGTGCTGAAAT | E46 | Δ*bgaA*::$P_c$-*erm* |
| P147 | TACGCCTTCTATCATGCCTTTGATCGCCCGT | | |
| P234 | CCCTTGTGTTCATAGCGAGGATAAGCA | K164 | Δ*pbp1a*::$P_c$-[*kan-rpsL*$^+$] |
| P235 | AGGCAAGCCTGCAACCATGGTCTTGAAA | | |
| P226 | GGTACGACAACGAAATGTCATACACTGCAC | K166 | Δ*pbp2a*::$P_c$-[*kan-rpsL*$^+$] |
| P227 | TCTGTTCCCGTGTGATCCGACAAATCCT | | |

| | | | |
|---|---|---|---|
| P1715 | TATCGGAAACTGTGAAGTGGGAAGCAACGT | IU19214 | Δ*macP*::$P_c$-*aad9* |
| P1716 | CTTTGACACGGTTATTGATTGCTTGGCGAG | | |
| P146 | TGGCCATTCATCGCTGGTCGTGCTGAAAT | IU19735 | Δ*bgaA*::$P_c$-*aad9* |
| P147 | TACGCCTTCTATCATGCCTTTGATCGCCCGT | | |
| P222 | CGTTCGTGTGGCGCTGCTTCAAATTGTT | IU13680 | Δ*pbp1b*::$P_c$-*aad9* |
| P522 | AACGGCAACCACCAAAGGAGAAACCAAGGA | | |
| P1737 | ACCAACCTCTTCTTCCCGTTTTGGTTATGC | IU18751 | Δ*rocS*::$P_c$-*erm* |
| P1740 | GCAACCAGAAGGTTAATTACAATCAAGGCT | | |
| **For construction of B2H plasmids** | | | |
| **Primer** | **Sequence** | **Template** | |
| **Construction of pKT25/pUT18C-*macP*-T32A (mutagenesis)** | | | |
| LN304 | AGAAGAGGTTGGTAAAATCTTAGCTTCCTCATTATCATCTAGCAAAG | pKT25/pUT18C-*macP*-WT | |
| LN305 | CTTTGCTAGATGATAATGAGGAAGCTAAGATTTTACCAACCTCTTCT | | |
| **Construction of pKT25/pUT18C-*macP*-T56A (mutagenesis)** | | | |
| LN306 | GACCTGAATCTTCAAGGCTTCCTGGCTAAAACCATGAT | pKT25/pUT18C-*macP*-WT | |
| LN307 | ATCATGGTTTTAGCCAGGAAGCCTTGAAGATTCAGGTC | | |
| **Construction of pKT25/pUT18C-*macP*-T32A T56A (2TA) (mutagenesis)** | | | |
| LN306 | GACCTGAATCTTCAAGGCTTCCTGGCTAAAACCATGAT | pKT25/pUT18C-*macP*-T32A | |
| LN307 | ATCATGGTTTTAGCCAGGAAGCCTTGAAGATTCAGGTC | | |
| **Construction of pKT25/pUT18C-*macP*-1-77 (truncation)** | | | |
| LN420 | CGTCTAGAGATGGGTAAATCTTTATTAACGGATGAAATG | pKT25/pUT18C-*macP*-WT | |
| LN422 | GCGAATTCTTAATTTCTCTTGGTATTTTCAAT | | |
| **Construction of pKT25/pUT18C-*macP*-1-87 (truncation)** | | | |
| LN420 | CGTCTAGAGATGGGTAAATCTTTATTAACGGATGAAATG | pKT25/pUT18C-*macP*-WT | |
| LN424 | GCGAATTCTTATAAGATTTTATTCAACTTAGAATTG | | |
| **Construction of pKT25/pUT18C-*macP*-L83A I86A L87A (mutagenesis)** | | | |
| BK26 | GCTAATAAAGCTGCTTTTGCGGTCATCTTTCTCTTG | pKT25/pUT18C-*macP*-WT | |
| BK27 | CTTAGAATTGAAGACATTTCTCT | | |
| **Construction of pKT25/pUT18C-*pbp2a*-R51A H53A K56A (mutagenesis)** | | | |
| BK28 | GCTTATGCTCTAACAGCTATTATCCTTATACTAGGTTTGAG | pKT25/pUT18C-*pbp2a*-WT | |
| BK29 | ACGCCAGAATTTTCGAATCG | | |

[a]Strains were constructed as described in *Experimental procedures.*

[b]$P_c$-*erm* and $P_c$-[*kan-rpsL*$^+$] cassettes are described in (Tsui *et al.*, 2011). $P_c$-*aad9* as described in (Lamanna *et al.*, 2022).

[c]All FLAG-tag (FLAG) fusions were made at the N-terminus of MacP. The amino acid sequence of the FLAG epitope is DYKDDDDK (Wayne *et al.*, 2010). C-terminal tagging of MacP resulted in loss of function as determined by transformation assays with Δ*pbp1a* (data not shown).

[d]*ireB*-L-FLAG$^3$ (IU14151) used in this study contained a linker sequence (L; GSAGSAAGSG). (Wayne *et al.*, 2010), followed by three tandem copies of the FLAG epitope (FLAG$^3$).

**Table S3.** Plasmids used in this study

| Plasmids | Genotype (description) | Antibiotic resistance[a] | Reference or source |
|---|---|---|---|
| pETPhosLink | *amp, pET derived (*T7 promoter, lac operator, His6-Tag*)*[b] | Amp$^R$ | (Novakova *et al.*, 2010) |
| pETPhos-*macP*[b] | *amp,* $P_{T7}$*-His6-macP* | Amp$^R$ | This study |
| pETPhos-*macP*-T32A[c] | *amp,* $P_{T7}$*- His6-macP-T32A* | Amp$^R$ | This study |
| pETPhos-*macP*-T56A | *amp,* $P_{T7}$*- His6-macP-T56A* | Amp$^R$ | This study |
| pETPhos-*macP*-2TA | *amp,* $P_{T7}$*- His6-macP-T32A T56A* | Amp$^R$ | This study |

**Bacterial two-hybrid (B2H) assay**

| **Plasmids** | **Genotype (description)** | Expressed protein/BATCH construct | Antibiotic resistance[a] | Reference or source |
|---|---|---|---|---|
| pKT25_*spd_0876* | *kan* $P_{lac}$*-cya(T25)-spd_0876* | T25-MacP | Kan$^R$ | (Stauberová *et al.*, 2024) |
| pUT18C_*spd_0876* | *amp* $P_{lac}$*-cya(T18)-spd_0876* | T18-MacP | Amp$^R$ | (Stauberová *et al.*, 2024) |
| pKT25_*spd_0876*-T32A | *kan* $P_{lac}$*-cya(T25)-spd_0876*-T32A | T25-MacP T32A | Kan$^R$ | This study |
| pUT18C_*spd_0876*-T32A | *amp* $P_{lac}$*-cya(T18)-spd_0876*-T32A | T18-MacP T32A | Amp$^R$ | This study |
| pKT25_*spd_0876*-T56A | *kan Plac-cya(T25)-spd_0876*-T56A | T25-MacP T56A | Kan$^R$ | This study |
| pUT18C_*spd_0876*-T56A | *amp* $P_{lac}$*-cya(T18)-spd_0876*-T56A | T18-MacP T56A | Amp$^R$ | This study |
| pKT25_*spd_0876*-T32 T56A | *kan* $P_{lac}$*-cya(T25)-spd_0876*-T32 T56A | T25-MacP T32A T56A | Kan$^R$ | This study |
| pUT18C_*spd_0876*-T32 T56A | *amp* $P_{lac}$*-cya(T18)-spd_0876*-T32 T56A | T18-MacP T32A T56A | Amp$^R$ | This study |
| pKT25_*spd_0876*-T32E | *kan* $P_{lac}$*-cya(T25)-spd_0876*-T32E | T25-MacP T32E | Kan$^R$ | This study |
| pUT18C_*spd_0876*-T32E | *amp* $P_{lac}$*-cya(T18)-spd_0876*-T32E | T18-MacP T32E | Amp$^R$ | This study |
| pKT25_*spd_0876*-T56E | *kan* $P_{lac}$*-cya(T25)-spd_0876*-T56E | T25-MacP T56E | Kan$^R$ | This study |
| pUT18C_*spd_0876*-T56E | *amp* $P_{lac}$*-cya(T18)-spd_0876*-T56E | T18-MacP T56E | Amp$^R$ | This study |
| pKT25_*spd_0876*-T32E T56E | *kan* $P_{lac}$*-cya(T25)-spd_0876*-T32E T56E | T25-MacP T32E T56E | Kan$^R$ | This study |
| pUT18C_*spd_0876*-T32E T56E | *amp* $P_{lac}$*-cya(T18)-spd_0876*-T32E T56E | T18-MacP T32E T56E | Amp$^R$ | This study |

| | | | | |
|---|---|---|---|---|
| pKT25_*spd_0876*-1-78 | *kan* $P_{lac}$-*cya(T25)-spd_0876-1-77* | T25-MacP-1-77 | $Kan^R$ | This study |
| pUT18C_*spd_0876*-1-78 | *amp* $P_{lac}$-*cya(T18)-spd_0876-1-77* | T18-MacP-1-77 | $Amp^R$ | This study |
| pKT25_*spd_0876*-1-88 | *kan* $P_{lac}$-*cya(T25)-spd_0876-1-87* | T25-MacP-1-87 | $Kan^R$ | This study |
| pUT18C_*spd_0876*-1-88 | *amp* $P_{lac}$-*cya(T18)-spd_0876-1-87* | T18-MacP-1-87 | $Amp^R$ | This study |
| pKT25_*spd_0876-L83A I86A L87A* | *kan* $P_{lac}$-*cya(T25)-spd_0876- L83A I86A L87A* | T25-MacP-L83 I86 L87A | $Kan^R$ | This study |
| pUT18C_*spd_0876- L83A I86A L87A* | *amp* $P_{lac}$-*cya(T18)-spd_0876- L83A I86A L87A* | T18-MacP-L83 I86 L87A | $Amp^R$ | This study |
| pKNT25_*spd_0339* | *kan* $P_{lac}$-*spd_0339-cya(T25)* | GpsB-T25 | $Kan^R$ | (Rued *et al.*, 2017) |
| pUT18_*spd_0339* | *amp* $P_{lac}$-*spd_0339-cya(T18)* | GpsB-T18 | $Amp^R$ | (Rued *et al.*, 2017) |
| pUT18_*spd_0339*-Y23A | *amp* $P_{lac}$-*spd_0339-Y23A-cya(T18)* | GpsB-Y23A-T18 | $Amp^R$ | (Cleverley *et al.*, 2019) |
| pUT18_*spd_0339*-V28A | *amp* $P_{lac}$-*spd_0339-V28A-cya(T18)* | GpsB-V28A-T18 | $Amp^R$ | (Cleverley *et al.*, 2019) |
| pUT18_*spd_0339*-D29A | *amp* $P_{lac}$-*spd_0339-D29A-cya(T18)* | GpsB-D29A-T18 | $Amp^R$ | (Cleverley *et al.*, 2019) |
| pUT18_*spd_0339*-L32A | *amp* $P_{lac}$-*spd_0339-L32A-cya(T18)* | GpsB-L32A-T18 | $Amp^R$ | (Cleverley *et al.*, 2019) |
| pUT18_*spd_0339*-D33A | *amp* $P_{lac}$-*spd_0339-D33A-cya(T18)* | GpsB-D33A-T18 | $Amp^R$ | (Cleverley *et al.*, 2019) |
| pUT18_*spd_0339*-I36A | *amp* $P_{lac}$-*spd_0339-I36A-cya(T18)* | GpsB-I36A-T18 | $Amp^R$ | (Cleverley *et al.*, 2019) |
| pKT25_*spd_1821* | *kan* $P_{lac}$-*spd_1821-cya(T25)* | T25-PBP2a | $Kan^R$ | (Cleverley *et al.*, 2019) |
| pUT18C_*spd_1821* | *amp* $P_{lac}$-*spd_1821-cya(T18)* | T18-PBP2a | $Amp^R$ | (Cleverley *et al.*, 2019) |
| pKT25_*spd_1821*-Δ32-37 | *kan* $P_{lac}$-*spd_1821-Δ32-37-cya(T25)* | T25-PBP2a-Δ32-37 | $Kan^R$ | (Cleverley *et al.*, 2019) |
| pKT25_*spd_1821*-Δ27-38 | *kan* $P_{lac}$-*spd_1821-Δ27-38-cya(T25)* | T25-PBP2a-Δ27-38 | $Kan^R$ | (Cleverley *et al.*, 2019) |
| pKT25_*spd_1821*-Δ26-45 | *kan* $P_{lac}$-*spd_1821-Δ26-45-cya(T25)* | T25-PBP2a-Δ26-45 | $Kan^R$ | (Cleverley *et al.*, 2019) |
| pKT25_*spd_1821*- R51A H53A K56A | *kan* $P_{lac}$-*spd_1821- R51A H53A K56A-cya(T25)* | T25-PBP2a-R51A H53A K56A | $Kan^R$ | This study |
| pUT18C_*spd_1821*- R51A H53A K56A | *amp* $P_{lac}$-*spd_1821- R51A H53A K56A-cya(T18)* | T18-PBP2a-R51A H53A K56A | $Amp^R$ | This study |
| pKNT25_spd_1479 | *kan* $P_{lac}$-*spd_1479-cya(T25)* | FtsZ-T25 | $Kan^R$ | (Rued *et al.*, 2017) |

| | | | | |
|---|---|---|---|---|
| pUT18_spd_1479 | *amp* $P_{lac}$-*spd_1479-cya(T18)* | FtsZ-T18 | $Amp^R$ | (Rued *et al.*, 2017) |
| pKT25_spd_1480 | *kan* $P_{lac}$-*cya(T25)-spd_1480* | T25-FtsA | $Kan^R$ | (Krupka *et al.*, 2012) |
| pUT18C_spd_1480 | *amp* $P_{lac}$-*cya(T18)-spd_1480* | T18-FtsA | $Amp^R$ | (Krupka *et al.*, 2012) |
| pKNT25_spd_0710 | *kan* $P_{lac}$-*spd_0710-cya(T25)* | EzrA-T25 | $Kan^R$ | (Rued *et al.*, 2017) |
| pUT18_spd_0710 | *amp* $P_{lac}$-*spd_0710-cya(T18)* | EzrA-T18 | $Amp^R$ | (Rued *et al.*, 2017) |
| pKT25_spd_1542 | *kan* $P_{lac}$-*spd_1542-cya(T25)* | StkP-T25 | $Kan^R$ | (Rued *et al.*, 2017) |
| pUT18C_spd_1542 | *amp* $P_{lac}$-*spd_1542-cya(T18)* | StkP-T18 | $Amp^R$ | (Rued *et al.*, 2017) |
| pKT25_spd_1474 | *kan* $P_{lac}$-*spd_1474-cya(T25)* | DivIVA-T25 | $Kan^R$ | (Rued *et al.*, 2017) |
| pUT18C_spd_1474 | *amp* $P_{lac}$-*spd_1474-cya(T18)* | DivIVA-T18 | $Amp^R$ | (Rued *et al.*, 2017) |
| pKT25_*spd_0305* | *kan* $P_{lac}$-*cya(T25)-spd_0305* | T25-FtsL | $Kan^R$ | (Perez *et al.*, 2021) |
| pUT18C_*spd_0305* | *amp* $P_{lac}$-*cya(T18)-spd_0305* | T18-FtsL | $Amp^R$ | (Perez *et al.*, 2021) |
| pKT25_*spd_0008* | *kan* $P_{lac}$-*cya(T25)-spd_0008* | T25-FtsB | $Kan^R$ | (Perez *et al.*, 2021) |
| pUT18C_*spd_0008* | *amp* $P_{lac}$-*cya(T18)-spd_0008* | T18-FtsB | $Amp^R$ | (Perez *et al.*, 2021) |
| pKT25_*spd_0600* | *kan* $P_{lac}$-*cya(T25)-spd_0600* | T25-FtsQ | $Kan^R$ | (Perez *et al.*, 2021) |
| pUT18C_*spd_0600* | *amp* $P_{lac}$-*cya(T18)-spd_0600* | T18-FtsQ | $Amp^R$ | (Perez *et al.*, 2021) |
| pKT25_*spd_0952* | *kan* $P_{lac}$-*cya(T25)-spd_0952* | T25-FtsW | $Kan^R$ | (Perez *et al.*, 2021) |
| pUT18C_*spd_0952* | *amp* $P_{lac}$-*cya(T18)-spd_0952* | T18-FtsW | $Amp^R$ | (Perez *et al.*, 2021) |
| pKT25_*spd_0306* | *kan* $P_{lac}$-*cya(T25)-spd_0306* | T25-PBP2x | $Kan^R$ | (Cleverley *et al.*, 2019) |
| pUT18C_*spd_0306* | *amp* $P_{lac}$-*cya(T18)-spd_0306* | T18-PBP2x | $Amp^R$ | (Cleverley *et al.*, 2019) |
| pKT25_*spd_0336* | *kan* $P_{lac}$-*cya(T25)-spd_0336* | T25-PBP1a | $Kan^R$ | (Cleverley *et al.*, 2019) |
| pUT18C_*spd_0336* | *amp* $P_{lac}$-*cya(T18)-spd_0336* | T18-PBP1a | $Amp^R$ | (Cleverley *et al.*, 2019) |
| pKT25_*spd_1486* | *kan* $P_{lac}$-*cya(T25)-spd_1486* | T25-PBP2b | $Kan^R$ | (Cleverley *et al.*, 2019) |
| pUT18C_*spd_1486* | *amp* $P_{lac}$-*cya(T18)-spd_1486* | T18-PBP2b | $Amp^R$ | (Cleverley *et al.*, 2019) |
| pKT25_*spd_0706* | *kan* $P_{lac}$-*cya(T25)-spd_0706* | T25-RodA | $Kan^R$ | (Perez *et al.*, 2021) |
| pUT18C_*spd_0706* | *amp* $P_{lac}$-*cya(T18)-spd_0706* | T18-RodA | $Amp^R$ | (Perez *et al.*, 2021) |

| | | | | |
|---|---|---|---|---|
| pKT25_*spd_2050* | *kan* $P_{lac}$-*cya(T25)-spd_2050* | T25-RodZ | Kan$^R$ | (Perez *et al.*, 2021) |
| pUT18C_*spd_2050* | *amp* $P_{lac}$-*cya(T18)-spd_2050* | T18-RodZ | Amp$^R$ | (Perez *et al.*, 2021) |
| pKT25_*spd_2045* | *kan* $P_{lac}$-*cya(T25)-spd_2045* | T25-MreC | Kan$^R$ | (Cleverley *et al.*, 2019) |
| pUT18C_*spd_2045* | *amp* $P_{lac}$-*cya(T18)-spd_2045* | T18-MreC | Amp$^R$ | (Cleverley *et al.*, 2019) |
| pKT25_*spd_1346* | *kan* $P_{lac}$-*cya(T25)-spd_1346* | T25-MpgA | Kan$^R$ | (Perez *et al.*, 2021) |
| pUT18C_*spd_1346* | *amp* $P_{lac}$-*cya(T18)-spd_1346* | T18-MpgA | Amp$^R$ | (Perez *et al.*, 2021) |
| pKNT25_spd_2044 | *kan* $P_{lac}$-*spd_2044-cya(T25)* | MreD-T25 | Kan$^R$ | (Perez *et al.*, 2021) |
| pUT18_spd_2044 | *amp* $P_{lac}$-*spd_2044-cya(T18)* | MreD-T18 | Amp$^R$ | (Perez *et al.*, 2021) |
| pKT25_*spd_1925* | *kan* $P_{lac}$-*cya(T25)-spd_1925* | T25-PBP1b | Kan$^R$ | (Lamanna *et al.*, 2022) |
| pUT18C_*spd_1925* | *amp* $P_{lac}$-*cya(T18)-spd_1925* | T18-PBP1b | Amp$^R$ | (Lamanna *et al.*, 2022) |
| pKT25_*spd_0180* | *kan* $P_{lac}$-*cya(T25)-ireB* | T25-IreB | Kan$^R$ | {Stauberová, 2024 #93} |
| pUT18C_*spd_0180* | *amp* $P_{lac}$-*cya(T18)-ireB* | T18-IreB | Amp$^R$ | {Stauberová, 2024 #93} |
| pKT25_*zip* | *kan* $P_{lac}$-*cya(T25)-zip* | T25-Zip | Kan$^R$ | (Karimova *et al.*, 1998) |
| pUT18C_*zip* | *amp* $P_{lac}$-*cya(T18)-zip* | T18-Zip | Amp$^R$ | (Karimova *et al.*, 1998) |
| pKT25 | *kan* $P_{lac}$-*cya(T25)* | T25 | Kan$^R$ | (Karimova *et al.*, 1998) |
| pUT18C | *amp* $P_{lac}$-*cya(T18)* | T18 | Amp$^R$ | (Karimova *et al.*, 1998) |

[a]Antibiotic resistance markers: Amp$^R$, ampicillin resistance; Kan$^R$, kanamycin resistance.

[b]pETPhosLink is a pET-derived expression vector containing a T7 promoter, *lac* operator, and an N-terminal His$_6$-tag (Novakova *et al.*, 2010) and was used as the backbone to construct pETPhos-*macP* and its derivative plasmids. pETPhos-derived plasmids were used for expression of recombinant His$_6$-tagged MacP proteins in *E. coli*.

[c]*macP* and mutant alleles were generated by site-directed mutagenesis as described in *Experimental procedures*.

**Table S4. Putative interactors of MacP determined by unbiased co-IP/MS[a]**

| Proteins that are in a complex with F-MacP (based on co-IP MS) | | | | |
|---|---|---|---|---|
| Protein | # unique peptides | | Function | Fold-change[c] |
| | $F^{-}$-*macP* | WT[b] | | |
| MacP | 6 (bait) | 0 | Regulator of aPBP2a | - |
| aPBP2a | 12 | 0 | ePG synthesis | - |
| IreB | 3 | 0 | Regulator of MurZ and MurA | - |
| PgdA | 3 | 0 | Peptidoglycan GlcNAc deacetylase | - |
| FtsX | 3 | 0 | ePG synthesis | - |
| GpsB | 1 | 0 | PG regulation | - |
| bPbp2x | 2 | 0 | sPG synthesis | - |
| bPbp2b | 2 | 0 | ePG synthesis | - |
| RodZ | 2 | 0 | ePG synthesis | - |
| MreC | 6 | 0 | ePG synthesis | - |
| MpgA | 9 | 3 | ePG synthesis | 13.36 |
| aPbp1b | 7 | 1 | ePG synthesis (?) | 4.28 |

[a] Number of peptides eluted from $F^{-}$-*macP* (IU17032) and WT (IU1824) strains following unbiased co-IP MS as described in *Experimental procedures*.

[b] Zero indicates peptides not detected in the untagged WT co-IP MS sample.

[c]"–" indicates that no peptide was detected in the WT sample. Fold change was calculated as the ratio of peptide abundance (LFQ intensity) in F-MacP relative to the WT control and was determined only for peptides detected in both samples.

### SUPPLEMENTAL FIGURE LEGENDS

**FIGURE S1. Genomic context, domain organization, and structural features of pneumococcal MacP and aPBP2a. (A)** Genomic region surrounding *macP* in the *S. pneumoniae* D39 chromosome. Genes shown include: *glmU* (*spd_0874*)-*spd_0875* (ADP-ribose pyrophosphatase) (operon_359): ***macP* (*spd_0876*)**: *mtnN* (*spd_0877*): *spd_0878* (*rocS/orfX*): *dnaQ* (*spd_0879*): and *whyD* (*spd_0880*). *macP* is predicted to be expressed as an independent transcriptional unit, but polarity reported here suggest that it is part of an operon. Immediately downstream of *macP* are: *mntN*, which encodes a bifunctional 5′-methylthioadenosine/S-adenosylhomocysteine nucleosidase; *spd_0878* (*rocS/orfX*), which encodes a chromosome partition regulator (Demuysere *et al.*, 2024, Mercy *et al.*, 2019); and *whyD* (*spd_0880*), which encodes a wall-teichoic acid (WTA) hydrolase (Flores-Kim *et al.*, 2022). **(B)** 2D-domain maps of MacP and PBP2a (not drawn to scale). Black lines indicate residues not assigned to known domains. Predicted transmembrane (TM) domains are shown in blue. Experimentally confirmed phosphorylated Thr32 and Thr56 in MacP are highlighted by green boxes. In PBP2a, the glycosyltransferase (GT), linker regulatory domain, and transpeptidase (TP) domains are indicated in green, gray, and red, respectively (Midonet *et al.*, 2023). **(C)** Predicted 3D structures of MacP and PBP2a generated using AlphaFold3. **(D)** Amino acid sequences of MacP and PBP2a, colored as in **(B)** to indicate domain organization and structural features. Predicted alpha-helical regions of MarP are boxed in gray. Phosphorylated Thr32 and Thr56 in MacP are in solid green boxes, and Thr7 and Thr37 that are potentially phosphorylated under stress conditions are in green dotted boxes. The GpsB-binding motifs in MacP and aPBP2a, the juxtamembrane region of aPBP2a, A77 that regulates GT activity, and the predicted disordered and α-helical region in the carboxyl terminus of aPBP2a are indicated. See text for additional details. **(E)** ConSurf conservation analysis of MacP across Streptococcus species. Residue annotations follow ConSurf's conventions (Glaser *et al.*, 2003): **e**- exposed residues; **b**- buried residues; **f**- predicted functional residues (highly conserved and exposed); **s**- predicted structural residues (highly conserved and buried); and **x**- insufficient data.

**Figure S2. Δ*macP* is synthetically lethal with Δ*pbp1a*, but not with Δ*pbp2a or* Δ*pbpb1b.* (A)** Tn-seq transposon insertion profiles in genomic regions containing *macP*, *pbp1a*, and *pbp2a*, in wild-type (*S. pneumoniae* D39 Δ*cps rpsL1*, strain IU1824), Δ*pbp1a* (IU18579), Δ*pbp2a* (IU13256), and Δ*pbp1b* (IU14697) strains. In vitro transposition reactions were performed with purified genomic DNA, *Magellan6* plasmid DNA, and purified MarC9 mariner transposase, followed by transformation, selection of transposon-inserted mutants, high-throughput sequencing (NextSeq 75, high-output), and analysis as described in the *Experimental procedures*. **(B)**Tables showing transposon insertion count ratios for *macP*, *pbp1a*, and *pbp2a* in each mutant background relative to WT. *p*-values were calculated using the Mann-Whitney U test.

**FIGURE S3. MacP phosphomimetic and phosphoablative mutants do not change the interaction with PBP2a and show no growth phenotype in the Δ*pbp1a* genetic background. (A)** Representative growth curves, doubling times, and growth yields in BHI broth for *macP* phosphomimetic and phosphoablative mutants, whose strain numbers are indicated in (**B**). Means ± SEMs are based on at least two biological replicates, and there were no statistically significant differences based on one-way ANOVA analysis with Dunnett's test. **(B)** Measurements of cell dimensions and representative phase-contrast micrographs (scale bar in left corner = 1 μm) of *macP* phosphoablative (T→A) mutants. Scatter plots (mean ± SEM) show cell length, width, aspect ratio, and relative volume. *p* values were obtained by one-way ANOVA analysis with the Kruskal-Wallis test. ns, not significant; *$p < 0.05$; **$p < 0.01$; ***$p < 0.001$ compared to WT. Data were from two biological replicates of ≥ 50 cells each, except for IU16788, which was from one experiment of 50 cells. **(C)** Growth curves, doubling times, growth yields, and representative micrographs showing a polar effect of a Δ*macP* mutant containing an insertion of the Pc-*erm* antibiotic cassette. Strains used: WT (IU1824), Δ*macP* (IU14699), Δ*macP*::$P_c$-[*kan-rpsL*$^+$] (IU14650), Δ*macP*::$P_c$-erm (IU17187) and *macP*$^+$-$P_c$-erm (IU17382). Growth curves were representative of two biological replicates. Statistical comparisons of significance was determined by one-way ANOVA with Dunnett's multiple-comparisons test: *$p < 0.05$; **$p < 0.01$; ***$p < 0.001$ compared to WT. **(D)** Representative growth curves of Δ*pbp1a* transformants*,* showing no growth defects in *macP* phosphoablative (T→A) or phosphomimetic (T→E) mutants. Strains used: WT D39 Δ*cps*  parent (IU1824), Δ*pbp1a* (IU13444), *macP*(T32A) Δ*pbp1a* (IU16726), *macP*(T56A) Δ*pbp1a* (IU16734), *macP*(T32A T56A) Δ*pbp1a* (IU16805)*, macP*(T32E) Δ*pbp1a* (IU16724), *macP*(T56E) Δ*pbp1a* (IU16730), and *macP*(T32E T56E) Δ*pbp1a* (IU16801). Growth curves are representative of two biological replicates, and there were no statistically significant differences in growth rates or yields of the mutants from WT by Dunnett's test. **(E)** Example of standard curve to ensure that relative protein amounts detected in western blots were in the linear range of detection. For each western blot, different amounts of protein samples (in this case, 2.5, 5.0, 10.0, 15.0 or 20.0 µL (black dots) of strain IU17032 (FLAG-*macP*)) were loaded on the same gel as the experimental samples to provide a standard curve of µL protein amount versus signal intensities (**Fig. 3E**) (Perez *et al.*, 2024). These plots were used to calculate the relative protein amounts in each sample lane by interpolation in the linear range of detection. TotalStain-Q NC (Azure Biosystems, AC2227) was used on the same blots to determine the relative amount of total protein in each lane and was used for normalization. Protein concentrations were not determined directly. Signal intensities obtained with different antibodies were normalized to total protein of each lane stained with Totalstain-Q. Standard curves of different amounts of protein samples stained with Totalstain-Q (not shown here) were also generated to verify signal linearity (Perez *et al.*, 2024). **(F)** Phosphoablative F-MacP mutants showed the same affinity as WT F-MacP for aPBP2a in co-IP experiments. Representative blots of the bait (F-macP) and prey (aPBP2a are shown). Four µL of lysate sample and 20 µL of output were loaded in each lane. The unknown band above the aPBP2a band (see **Fig. 5C**) was faint in this blot, and the position of aPBP2a is marked by a red arrow. Only the band corresponding to aPBP2a was detected after co-IP (**Fig. 6C***)*. Relative amounts of immunoprecipitated aPBP2a are indicated for this blot. The single phosphoablative mutants were tested once, and the double *macP*(T32A T56A) mutant (IU17035) was tested twice with similar results. Strains used: F-*macP* (IU17032), F-*macP*(T32A) (IU16978), F-*macP*(T56A) (IU16980) and F-*macP*(T32A T56A) (IU17035). **(G)** B2H assay results confirming interaction of MacP phosphoablative variants with aPBP2a. Plasmid pairs pKT25/pUT18C and pKT25-*zip*/pUT18C-*zip* were used as negative (-ve) and positive (+ve) controls, respectively. Plates were imaged after 40 h at 30 °C. B2H assays were performed as described in the *Experimental procedures*. **(H)** Phosphorylation of F-MacP at Thr32 and Thr56 determined by IP and detection with anti P-Thr antibody. Left panels, representative western blots of precipitated F-tag proteins probed with anti-MacP (top) and anti-P-Thr (bottom) antibodies. Right panel, quantitation of the phosphorylation signal from two independent experiments, each with two technical replicates. Strains used: WT F-MacP (Sp480); F-MacP(T32A) (Sp486); F-MacP(T56A) (Sp487); and F-MacP (T32 56A) (Sp481). Control (Ctrl) represents parental strain Sp339 (IU1945).

**FIGURE S4. Summary of results for MacP and aPBP2a mutants analyzed in this study. (A)** Summary of MacP mutants indicating locations of amino acid changes, synthetic lethality with Δ*pbp1a*; expression of F-tag variants relative to WT F-MacP (biological replicates are in parentheses); and whether a complex was detected between the F-macP variants and aPBP2a. **(B)** Summary of aPBP2a mutants indicating locations of amino acid changes and synthetic lethality with Δ*pbp1a*. Data were compiled from figures and tables that are described in the text.

**FIGURE S5. MacP cytoplasmic deletion mutants retain activation of aPBP2a. (A)** Phosphoablative, phosphomimetic, and cytoplasmic deletion mutants of MacP retained interaction with aPBP2a (red arrow) and GpsB (green arrow). Representative western blots are shown of co-IP experiments of interactions between F-MacP variants (bait) and aPBP2a (prey) performed as described in *Experimental procedures*. 4 µL of lysate input (left panel), and 15 µL of eluted protein (right panel) were loaded per lane. Similar results were obtained from two biological replicates showing no changes in binding of the F-MacP mutant variants compared to WT F-MacP to aPBP2a or GpsB. Strains used: F-*macP* WT (IU17032); F-*macP*(T32A T56A)

(IU16978); F-*macP*(T32E T56E) (IU16978); F-*macP*(Δ46-53) (IU18345); F-*macP*(Δ30-33) (IU18346); and F-*macP*(Δ21-58) (IU18348). **(B)** Mutant variants with deletions in the cytoplasmic domain of MacP did not change growth compared to WT in BHI broth. Representative growth curves are shown with doubling times and growth yields from at least two biological replicates. Strains used: MacP WT (IU1824); Δ*macP* (IU14699); *macP*(Δ5-8) (IU17228); *macP*(Δ21-58) (IU17386); *macP*(Δ30-33) (IU17384); *macP*(Δ46-53) (IU17232); *macP*(1-77) (IU17236); and *macP*(1-87) (IU17240). **(C)** Mutant variants with deletions in the cytoplasmic domain of MacP or MacP(R76A N80A) activated aPBP2a in the absence of aPBP1a. Representative growth curves are shown with doubling times and growth yields from at least two biological replicates. Strains used: MacP WT (IU1824); Δ*pbp1a* (IU13444); Δ*pbp1a macP(*Δ5-8) (IU17271); Δ*pbp1a macP(*Δ46-53) (IU17275); Δ*pbp1a macP(*Δ30-33) (IU17456); Δ*pbp1a macP(*Δ21-58) (IU17460); and Δ*pbp1a macP*(R76A N80A) (IU19605). **(D)** Mutant variants with deletions in the cytoplasmic domain of MacP or MacP(R76A N80A) and Δ*pbp1a* had the same shape and size as a Δ*pbp1a* mutant. Scatter plots (means ± SEM) are shown comparing cell length, width, aspect ratio (length/width), and relative volume are shown for the strains in (**C**). Cell parameters were compared to WT by one-way ANOVA analysis with the Kruskal-Wallis test: ns, not significant; $*p < 0.05$; $**p < 0.01$; $***p < 0.001$ compared to WT. Data are from two biological replicates with ≥ 50 cells analyzed per replicate.

**Figure S6. MacP is in a complex with IreB**. Silver-stained SDS-PAGE gel from co-IP experiments performed as described in *Experimental procedures* for WT strain IU1824 (non-FLAG-tag control) and strain IU14151 that produced IreB-L-FLAG$^3$ (bait). Lanes in the boxed area were analyzed by LC-MS. The white arrow marks the expected position of IreB-FLAG$^3$ in the right lane. The table shows the number of unique peptides from MacP identified by LC-MS from the IreB-L-FLAG$^3$ lane that were absent from the non-FLAG-tag control lane. A complex between MacP and IreB was also detected by a direct co-IP/MS procedure (see **Table S4**).

**FIGURE S7. Identification of amino acids in MacP(TM) and the juxtamembrane region of aPBP2a(TM) important for MacP activation of aPBP2a. (A)** Top panel, western blot of *E. coli* lysates showing that MacP(N77(stop)) or MacP(L87(stop) mutant variants lacking TM domains were stabilized in fusion constructs used for B2H assays in *E. coli*. This result contrasted with lack of expression of MacP(N77(stop)) or MacP(L87(stop) in *S. pneumoniae* (**Fig. 3C** and **3D**). Lysates of *E. coli* DH5α strains transformed with the B2H constructs indicated below the lanes were immunoblotted with anti-MacP antibody as described in *Experimental procedures*. Bottom panel, B2H assay of interactions between MacP(N77(stop)) or MacP(L87(stop)) with MacP, aPBP2a, and GpsB. WT MacP showed self-interaction and interaction with aPBP2a or GpsB, whereas MacP(N77(stop)) or MacP(L87(stop)) did not show self-interactions or detectable interactions with MacP, aPBP2a, or GpsB, despite expression of the fusion constructs. Plasmid pairs pKT25/pUT18C and pKT25-zip/pUT18C-zip were used as negative (-ve) and positive (+ve) controls. respectively. Similar results were obtained in two biological replicates. See text for additional details. **(B)** (Top) MacP (blue) and aPBP2a (red) residues that were predicted to contribute to the MacP-aPBP2a interaction interface. The interacting region was identified from the AlphaFold3 predicted structure on which polar contacts were identified using PyMOL with a cutoff distance of <4 Å. (Bottom) PDBsum interaction diagram illustrating residue–residue contacts between MacP (called Chain A) and aPBP2a (called Chain B). The majority of contacts are hydrophobic (Leu/Ile/Phe-rich clusters), complemented by polar interactions (Asn80–His53, Asn84–Lys56, Glu72–Arg51) and a predicted salt bridge involving Glu72–Arg51. **(C)** B2H assays showed some level of interaction between MacP(L83A I86A L87A) or PBP2a(R51A H53A K56A) and aPBP2a or MacP, respectively, in *E. coli*. B2H assays were done as described in *Experimental procedures*. Plasmid pairs pKT25/pUT18C and pKT25-*zip*/pUT18C-*zip* were used as negative (-ve) and positive (+ve) controls, respectively. Plates were imaged after 40 h 30 °C. Similar results were obtained in two biological replicates. **(D)** Co-IP assays showed a substantial decrease in binding between F- MacP(L83A I86A L87A) and aPBP2a (red arrow), and reduced binding between F-MacP(R76A N80A) and aPBP2a. Co-IP assays were performed as described in *Experimental procedures*. Numbers shown refer to relative aPBP2a amounts recovered in this representative blot. Strains used: WT (IU1824); F-*macP* (IU17032); F-*macP*(R76A N80A) (IU19524); F-*macP*(L83A I86A L87A) (IU19528); and F-*macP*(M101A L103A) (IU19529). **(E)** Summary table showing the mean ± SEM from two biological replicates of the relative amount of aPBP2a bound to variants of F-MacP. See text for additional details. **(F)** GFP-MacP(L83A I86A L87A) and other GFP-MacP variants localized to midcell similar to WT GFP-MacP, indicating that these amino acid changes did not alter subcellular localization. Phase-contrast, fluorescent-microscopy, and merged images are shown of cells grown exponentially to $OD_{620}$ ≈0.15 in BHI broth. Microscopy was performed at least twice as described in *Experimental procedures*, and scale bars (in left corners) = 1 µm.*,* Strains used: *macP*$^+$ WT (IU1824); *gfp*-L-*macP* (IU19947); *gfp*-L-*macP* Δ*pbp2a::*$P_c$-[*kan-rpsL*$^+$] (IU21336); *gfp*-L-*macP* (R76A N80A) (IU21339); *gfp*-L-*macP*(L83A I86A L87A) (IU21340), and *gfp*-L-*macP*(S68A R71A R70A K75A R76A) (IU21345).

**FIGURE S8. *pbp2a*(A77T) bypasses the requirement of MacP but only results in a minimal change in cell width. (A)** Representative growth curves in BHI broth of strains expressing MacP during aPBP1a depletion (top) and strains lacking MacP to activate aPBP2a upon aPBP1a depletion (bottom). The result confirms the previous finding that *pbp2a*(A77T) obviates the requirement for MacP activation in the absence of aPBP1a (Midonet, *et al*., 2023). In contrast, change of the adjacent amino acid aPBP2a(K78A) did not bypass the requirement for MacP activation in the absence of aPBP1a. Similar results were obtained for two biological replicates. +Zn (0.4 mM $Zn^{+2}$/0.04 mM $Mn^{+2}$), which was added to overnight cultures and used as starters for the growths shown. See *Experimental procedures* for other details about growths. Strains used in top panel: *pbp2a*$^+$ *pbp1a*$^+$ WT (IU1824); *pbp2a*$^+$ Δ*pbp1a*//$P_{Zn}$-*pbp1a* (IU14357); *pbp2a*(A77T) Δ*pbp1a*//$P_{Zn}$-*pbp1a* (IU19240); and *pbp2a*(K78A) Δ*pbp1a*//$P_{Zn}$-*pbp1a* (IU19242). (Bottom) Strains used in bottom panel: *pbp2a*$^+$ *macP*$^+$ *pbp1a*$^+$WT (IU1824), Δ*macP* Δ*pbp1a*//$P_{Zn}$-*pbp1a* (IU19431), *pbp2a*(A77T) Δ*macP* Δ*pbp1a*//$P_{Zn}$-*pbp1a* (IU19433), and *pbp2a*(K78A) Δ*macP* Δ*pbp1a*//$P_{Zn}$-

*pbp1a* (IU19435). **(B)** Representative growth curves with cell length and width measurements (mean ± SEM) of strains: WT (IU1824); Δ*pbp2a* (IU13256); *pbp2a*(A77T) (IU20380); and *pbp2a*(K78A) (IU20382) grown in BHI broth. Comparisons to WT lengths and widths were made by one-way ANOVA analysis with the Kruskal-Wallis test: ns, not significant; *$p < 0.05$; **$p < 0.01$; ***$p < 0.001$. Data are from two biological replicates with ≥ 50 cells analyzed per replicate. **(C)** Representative phase contrast microscopy images taken between 3 and 3.5 h of strains in **(B)**. Scale bar = 1 µm. **(D)** Representative growth curves of strains in **(B)** in THY broth. Growth curves are representative of two biological replicates and show no statistically significant difference from WT by Dunnett's test.

**FIGURE S9. Pentuple alanine (Ala) substitution in the SRRIENTKR motif of MacP impairs function and expression. (A)** Representative quantitative western blot showing the relative amount of F-MacP(S68A R69A R70A K75A R76A) compared to WT F-MacP in cells growing exponentially in BHI broth. See *Experimental procedures* for detail. Strains used: *MacP*$^+$ WT (IU1824); F-*macP* (IU17032); and F-*macP*(S68A R69A R70A K75A R76A) (IU21377). **(B)** Summary table (mean ± SEM) of the relative amount of MacP (S68A R69A R70A K75A R76A) from three biological replicates. **(C)** Transformation of *macP*(S68A R69A R70A K75A R76A) with Δ*pbp1a* resulted in tiny, sick colonies that streaked out and could be regrown. Plate pictures were taken after 20 h incubation at $37^{o}$C. Similar results were obtained in two biological replicates. **(D)** Left panel, input controls for co-IP assays shown in **Fig. 8D**, which is shown again in the right panel for comparison. Similar results were obtained in three biological replicates. See *Experimental procedures* and the text for additional details. Strain used: *macP*$^+$ WT (IU1824); F-*macP* (IU17032); and F-*macP*(S68A R69A R70A K75A R76A) (IU21377).

**FIGURE S10. Tn-seq demonstrates that Δ*pbp1a* is synthetically lethal with Δ*phpP* in cells growing exponentially in BHI broth**. **(A)** Predicted 3D structure of StkP(*Spn*) generated using AlphaFold3. Predicted extracellular PASTA domains P1, P2, P3, and P4 with indicated amino-acid numbers are indicated. **(B)** Mini-Mariner *Malgellan6* Tn-Seq transposon insertion profile for the genomic region covering *sun*, *phpP*, *stkP*, and *spd_1541* in the genomes of the unencapsulated WT parent (D39 Δ*cps rpsL1*, IU1824), Δ*pbp1a* (IU18579), Δ*pbp2a* (IU13256), or Δ*pbp1b* (IU14697). Transposition, library outgrowth, sequencing (NextSeq 75), and analysis were performed as described in *Experimental procedure*. Insertions in WT were detected in regions encoding P3 and P4, but not elsewhere in *stkP*. The first TA insertion in *stkP* occurred in the WT strain at the TAT(Y515) codon (indicated), where the Tn insertion created a TAA stop codon. while No insertion occurred at the upstream TTA(L512) codon, indicating that *stkP*(M1-L512) is essential for viability (Tsui, *et al.*, 2023). **(C)** Predicted three-dimensional structure of PhpP (*Spn*) generated using AlphaFold3. **(D)** Transformation assay showing synthetic lethality between Δ*pbp1a* and Δ*phpP*. Parent D39 Δ*cps rpsL1* strain (IU1824) and Δ*pbp1a* (IU18579) were transformed with Δ*phpP*::Pc-*erm* or control amplicon Δ*bgaA*::Pc-*erm* as described in the *Experimental procedures.* Representative TSAII-BA plates were imaged after ≈20 h incubation at 37°C. **(E)** Transformation efficiency and colony morphology in the suppressed *murZ*(D280Y) Δ*stkP* background, which lacks Ser/Thr protein phosphorylation. The indicated recipient strains were transformed with the listed amplicons as described in *Experimental procedures*, and colony numbers and appearance were assessed after 22 h incubation. Similar results were obtained in two biological replicates. N/A, not applicable because Δ*pbp1a* is already in the strain.

**FIGURE S11. Δ*pbp1a* is synthetically lethal with Δ*rocS*.** **(A)** (Left) Mini-Mariner *Malgellan6* Tn-Seq transposon insertion profile for the genome region containing *spd_0875*, *macP*, *mntN, rocS*, and *dnaQ* in strains: unencapsulated WT parent (D39 Δ*cps rpsL1*, IU1824); Δ*pbp1a* (IU18579); Δ*pbp2a* (IU13256); and Δ*pbp1b* (IU14697). Transposition reactions and analysis were performed as described in *Experimental procedures*. (Right) Predicted 3D structure of RocS(*Spn*) generated using AlphaFold3. **(B)** Transformation assay confirming synthetic lethality between Δ*pbp1a* and Δ*rocS*. The following strains: parent D39 Δ*cps rpsL1* strain (IU1824); Δ*pbp1a* (IU18597); and Δ*macP* (IU14699) were transformed with a Δ*rocS*::Pc-*erm* or control amplicons Δ*bgaA*::Pc-*erm* and Δ*pbp1b*::Pc-*erm,* as described in *Experimental procedures.* (Left) Representative transformation plates showing colonies on TSAII-BA plates after ≈20 h incubation. (Right) Summary of colony numbers and morphology after 22 h. No transformants were recovered for Δ*pbp1a* with Δ*rocS*::$P_c$-*erm*, whereas all controls yielded >500 colonies. Replicate numbers are in parentheses.

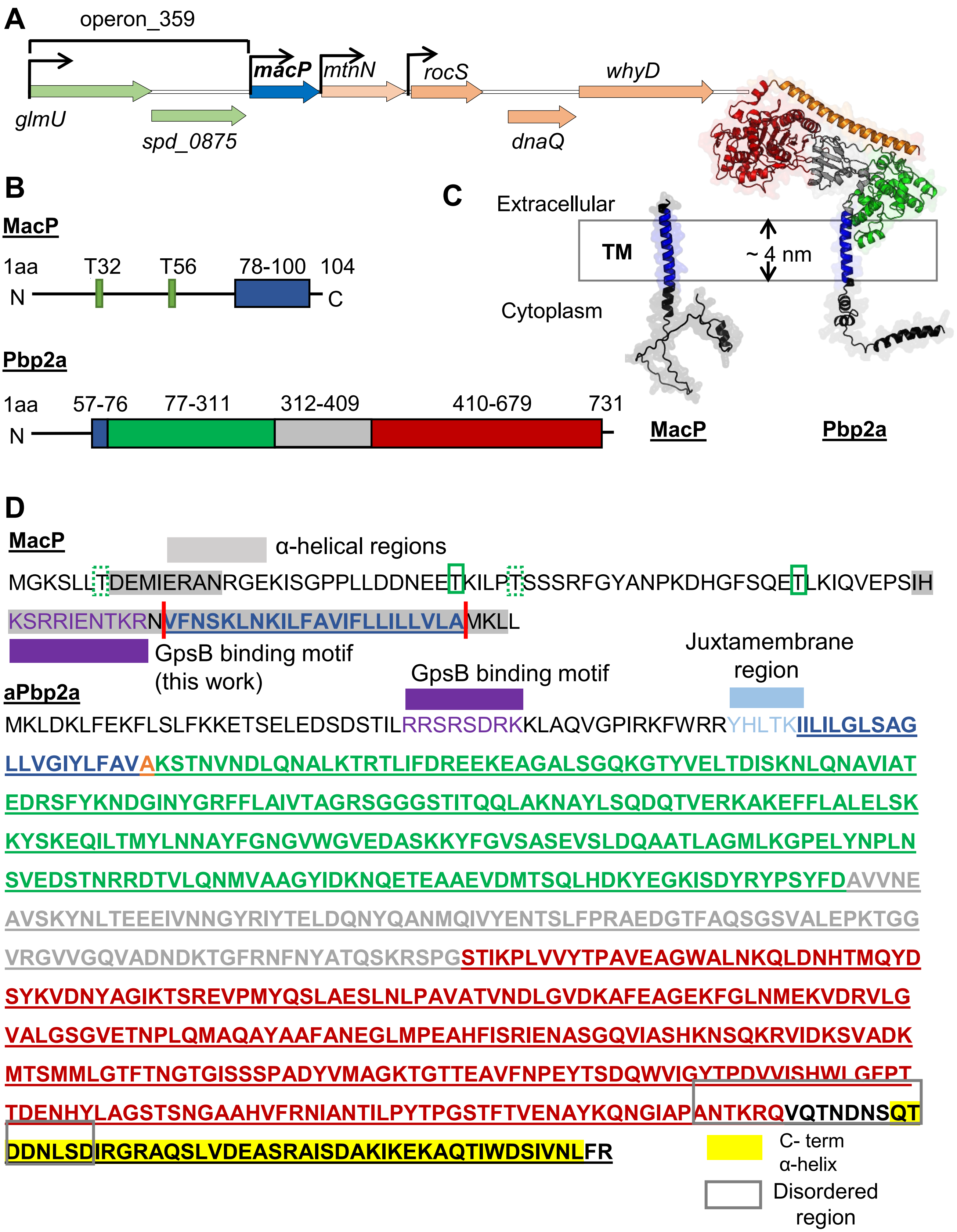


FIGURE S1 (continued)

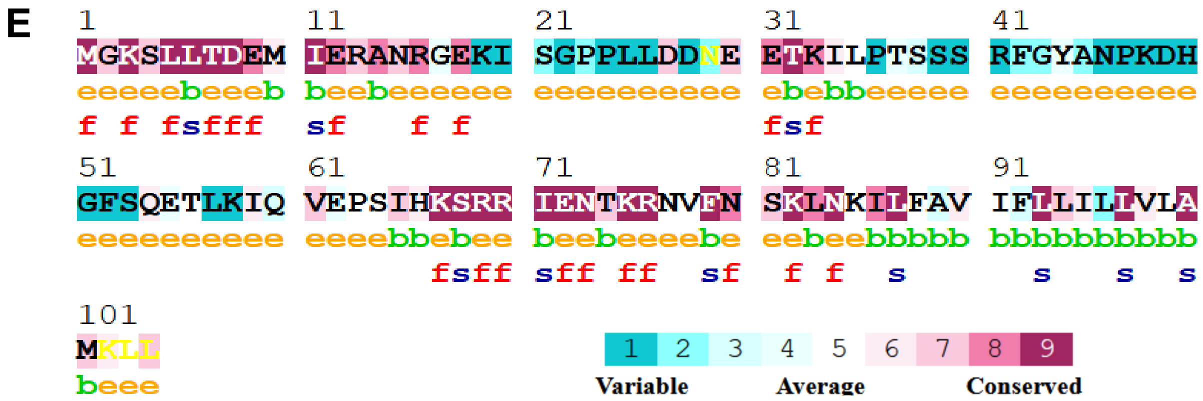


FIGURE S1 (continued)

**A**

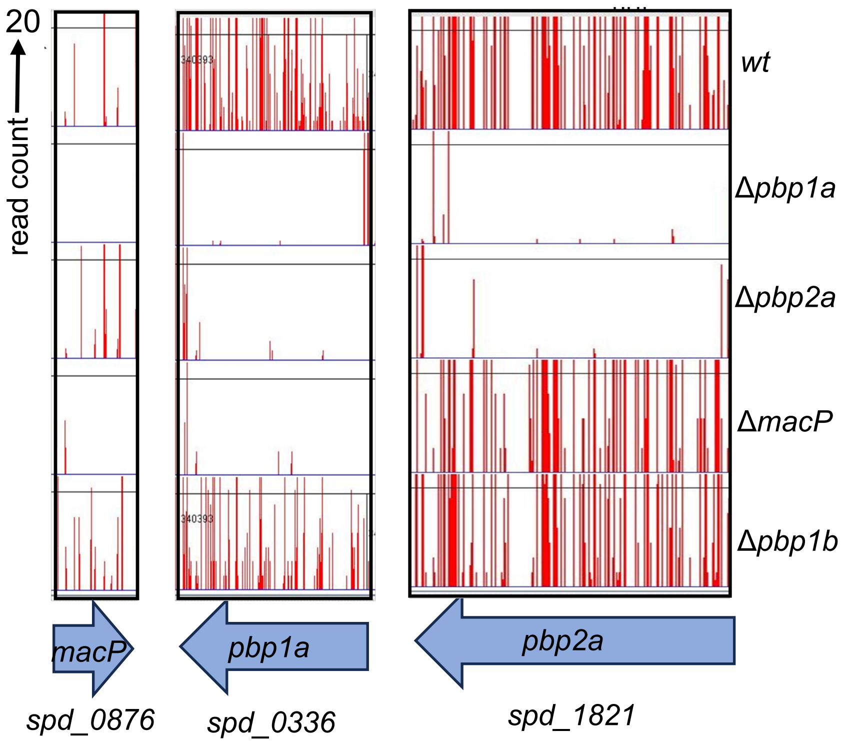


**B**

***macP* (*spd_0876*)**

| Strain | Count Ratio (mut/WT) | p-value |
|---|---|---|
| Δ*pbp1a* | 0.00 | 0.002 |
| Δ*pbp2a* | 0.95 | 0.405 |
| Δ*macP* | 0.08 | 0.036 |
| Δ*pbp1b* | 0.64 | 0.311 |

***pbp1a* (*spd_0336*)**

| Strain | Count Ratio (mut/WT) | p-value |
|---|---|---|
| Δ*pbp1a* | 0.07 | 0.000 |
| Δ*pbp2a* | 0.08 | 0.000 |
| Δ*macP* | 0.02 | 0.000 |
| Δ*pbp1b* | 0.41 | 0.001 |

***pbp2a* (*spd_1821*)**

| Strain | Count Ratio (mut/WT) | p-value |
|---|---|---|
| Δ*pbp1a* | 0.02 | 0.000 |
| Δ*pbp2a* | 0.03 | 0.000 |
| Δ*macP* | 1.06 | 0.042 |
| Δ*pbp1b* | 0.61 | 0.051 |

**FIGURE S2**

**A**

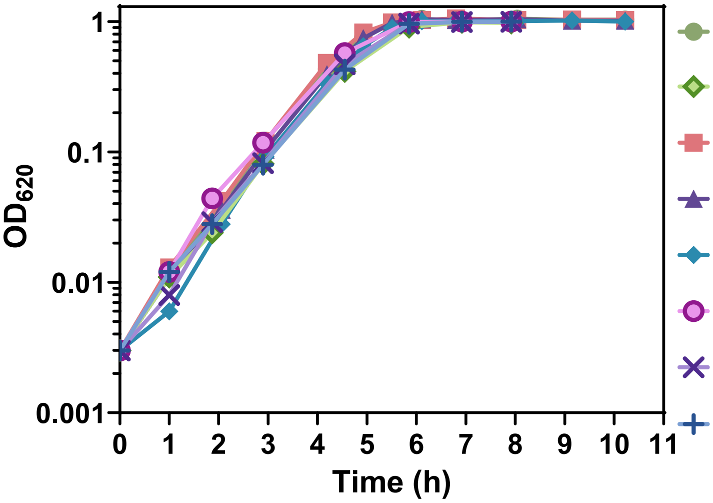


| | Mean ± SEM | |
|---|---|---|
| | Doubling time (min) | Growth yield ($OD_{620}$) |
| WT | 40.6 ± 1.0 | 1.00 ± 0.02 |
| Δ*macP* | 39.6 ± 1.5 | 0.99 ± 0.01 |
| *macP*(T32A) | 37.1 ± 0.9 | 1.01 ± 0.02 |
| *macP*(T56A) | 38.3 ± 1.2 | 0.98 ± 0.01 |
| *macP*(T32A T56A) | 38.5 ± 1.1 | 1.05 ± 0.03 |
| *macP*(T32E) | 41.7 ± 0.9 | 1.01 ± 0.02 |
| *macP*(T56E) | 38.4 ± 0.9 | 0.98 ± 0.01 |
| *macP*(T32E T56E) | 40.8 ± 0.4 | 1.03 ± 0.02 |

**B**

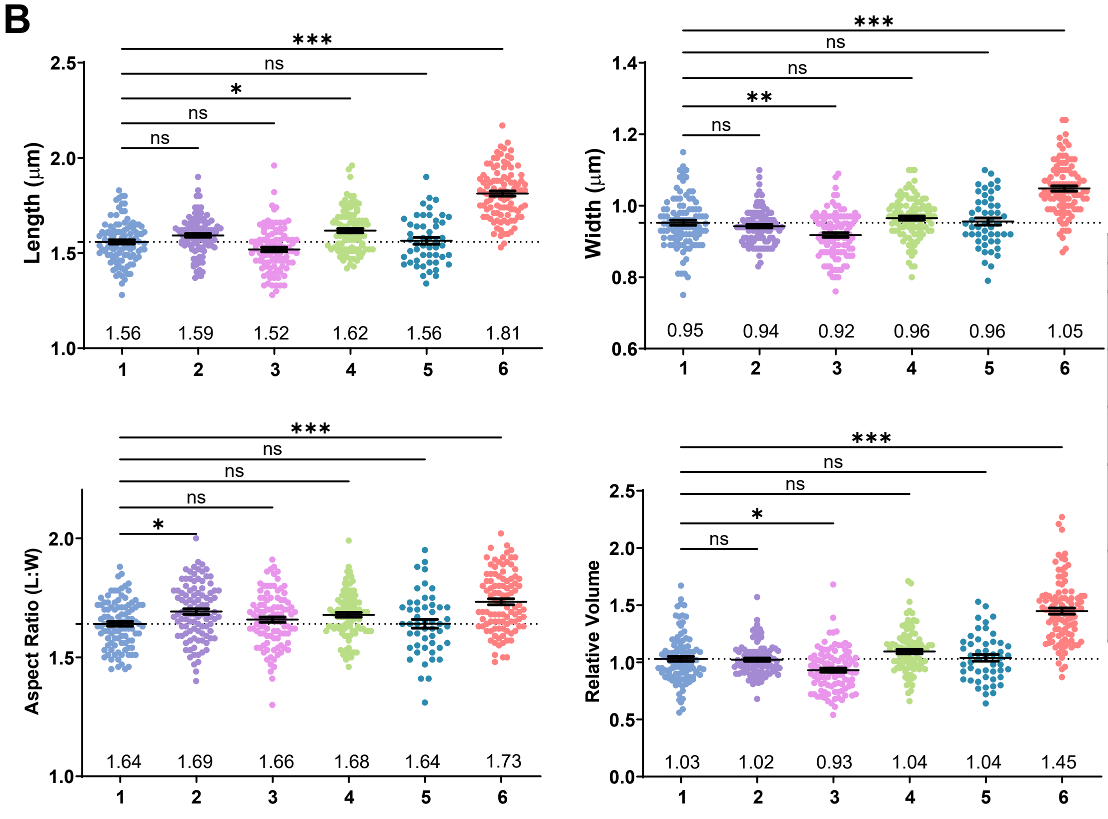


| | Genotype |
|---|---|
| 1 | WT |
| 2 | Δ*macP* (IU14699) |
| 3 | *macP*(T32A) (IU14822) |
| 4 | *macP*(T56A) (IU16618) |
| 5 | *macP*(T32A T56A) (IU16788) |
| 6 | Δ*macP*::$P_c$-[*kanrpsL*$^+$] (IU14650) |

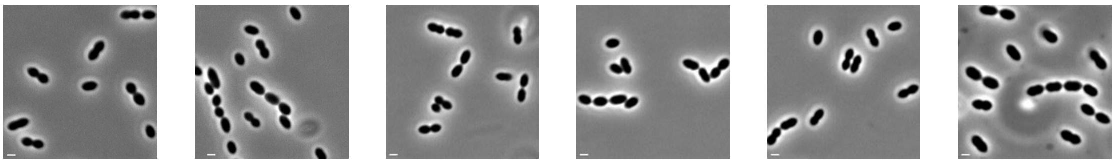

WT (IU1824) | Δ*macP* (IU14699) | *macP*(T32A) (IU14822) | *macP*(T56A) (IU16618) | *macP*(T32A T56A) (IU16788) | Δ*macP*::$P_c$-[*kanrpsL*$^+$] (IU14650)

**FIGURE S3 (continued)**

**C**

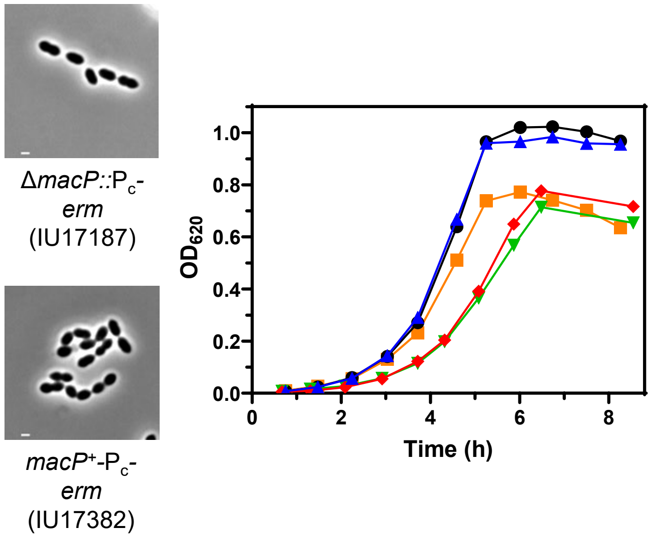


| Strain | Mean ± SEM: Doubling time (min) | Mean ± SEM: Growth yield ($OD_{620}$) |
|---|---|---|
| WT | 37.8 ± 1.3 | 0.99 ± 0.01 |
| Δ*macP* markerless | 37.0 ± 1.1 | 0.98 ± 0.01 |
| Δ*macP*::Pc-[*kan-rpsL*$^+$] | 39.6 ± 2.8 | 0.75 ± 0.01*** |
| Δ*macP*::Pc-*erm* | 47.0 ± 1.5** | 0.70 ± 0.01*** |
| *macP*$^+$-Pc-*erm* | 49.1 ± 2.6*** | 0.71 ± 0.04*** |

**D**

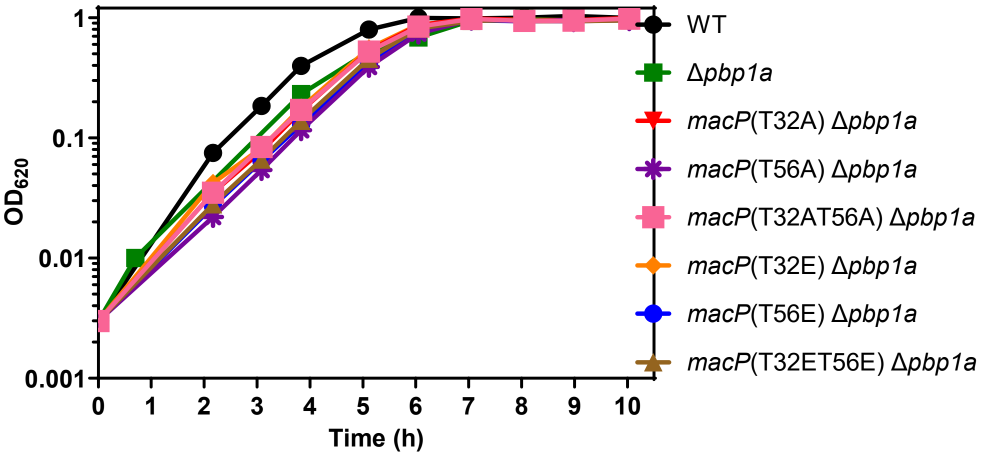


| Strain | Mean ± SEM: Doubling time (min) | Mean ± SEM: Growth yield ($OD_{620}$) |
|---|---|---|
| WT | 40 ± 0.7 | 1.00 ± 0.01 |
| Δ*pbp1a* | 41 ± 0.6 | 0.91 ± 0.01 |
| *macP*(T32A) Δ*pbp1a* | 41 ± 1.5 | 0.96 ± 0.01 |
| *macP*(T56A) Δ*pbp1a* | 41 ± 0.5 | 0.94 ± 0.01 |
| *macP*(T32AT56A) Δ*pbp1a* | 42 ± 1.6 | 0.95 ± 0.00 |
| *macP*(T32E) Δ*pbp1a* | 42 ± 2.4 | 0.95 ± 0.01 |
| *macP*(T56E) Δ*pbp1a* | 40 ± 0.7 | 0.95 ± 0.00 |
| *macP*(T32ET56E) Δ*pbp1a* | 43 ± 0.1 | 0.96 ± 0.00 |

**E**

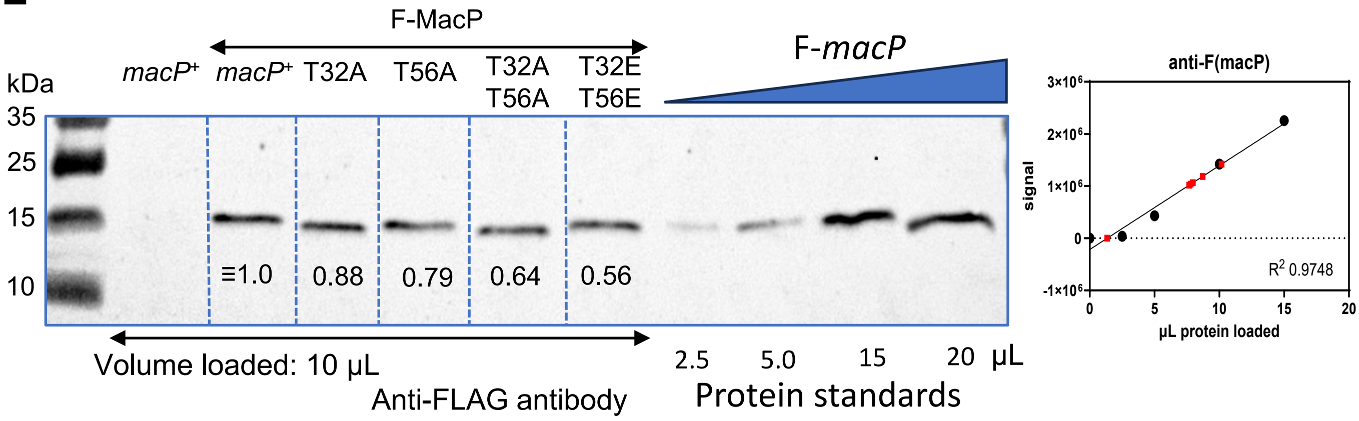


**FIGURE S3 (continued)**

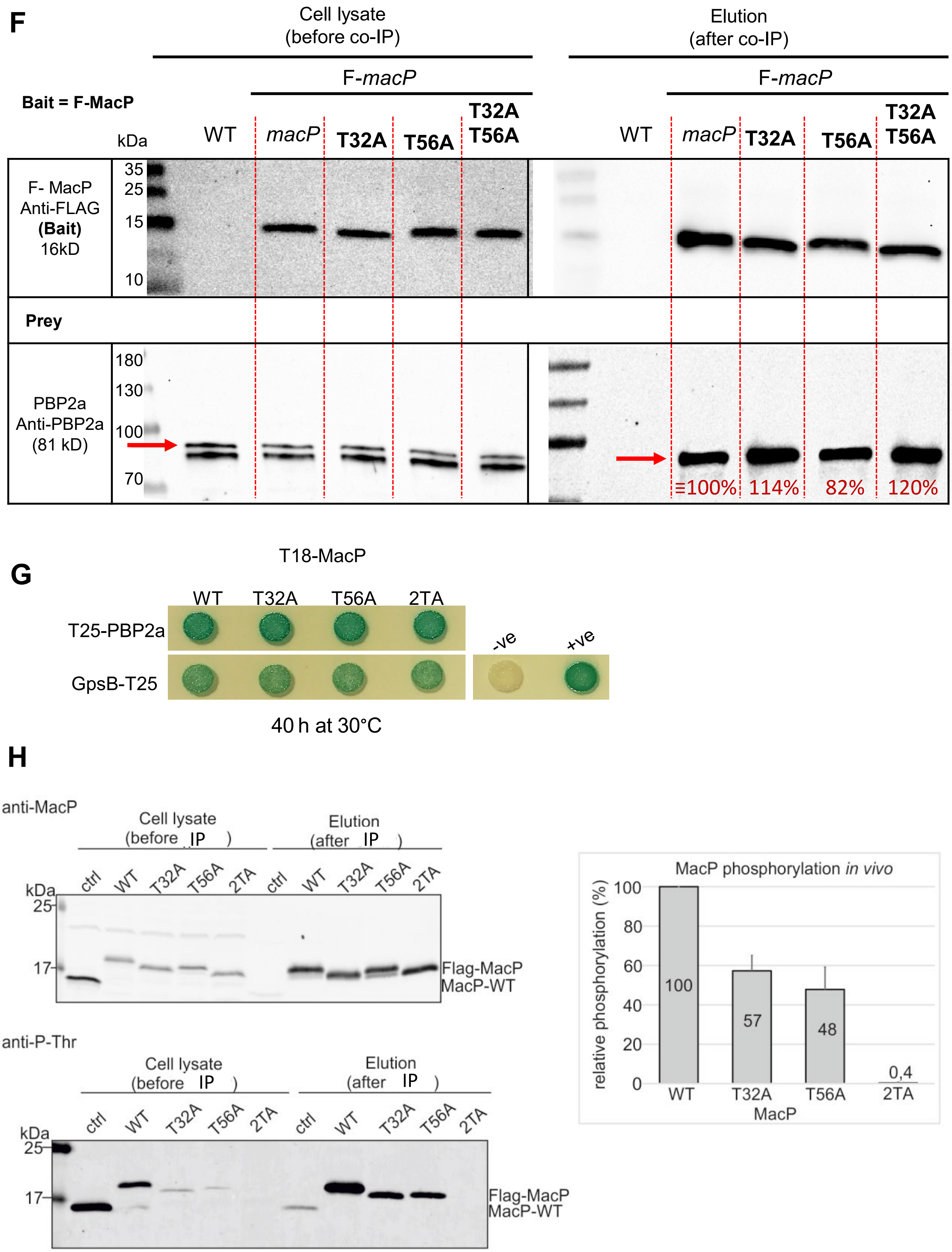


**FIGURE S3 (continued)**

**A**

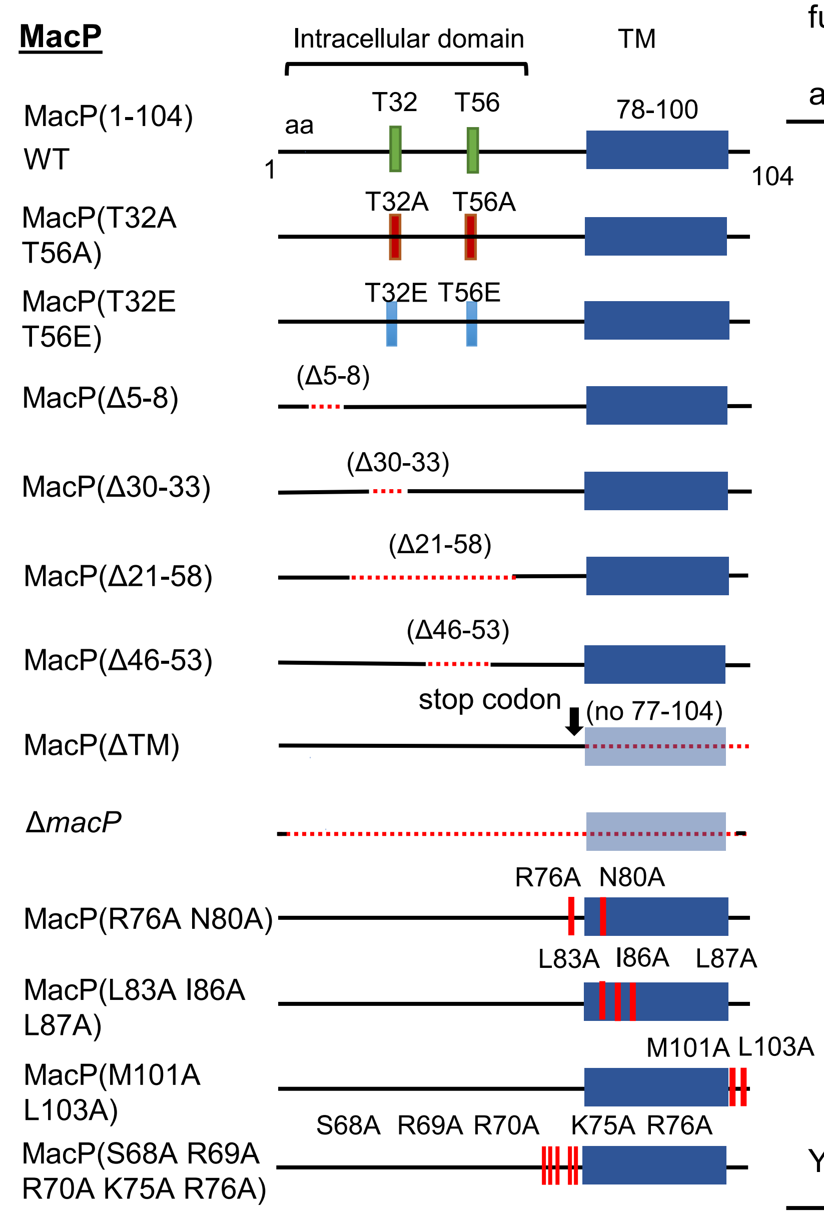


| Mutation functional w/o aPBP1a? | F-tagged protein expression level | Complex with aPbp2a |
|---|---|---|
| WT | 1.00 (3+) | Yes |
| Yes | 0.7 ± 0.3 (3+) | Yes |
| Yes | 0.7 ± 0.1 (3) | Yes |
| Yes | 0.8 ± 0.03 (2) | nd |
| Yes | 0.7 ± 0.1 (2) | Yes |
| Yes | 0.1 ± 0.03 (3) | Yes |
| Yes | 0.9 ± 0.1 (3) | Yes |
| No | 0.00 (3) | No |
| No | 0.00 (3+) | No |
| Yes | 0.7 ± 0.03 (2) | Yes |
| No | 0.5 ± 0.04 (2) | No |
| Yes | 0.9 ± 0.06 (2) | Yes |
| Yes (sick) | 0.3 ± 0.04 (3+) | Yes |

= phosphoablative = phosphomimetic

Key
= stop codon
= deleted region

**FIGURE S4 (continued)**

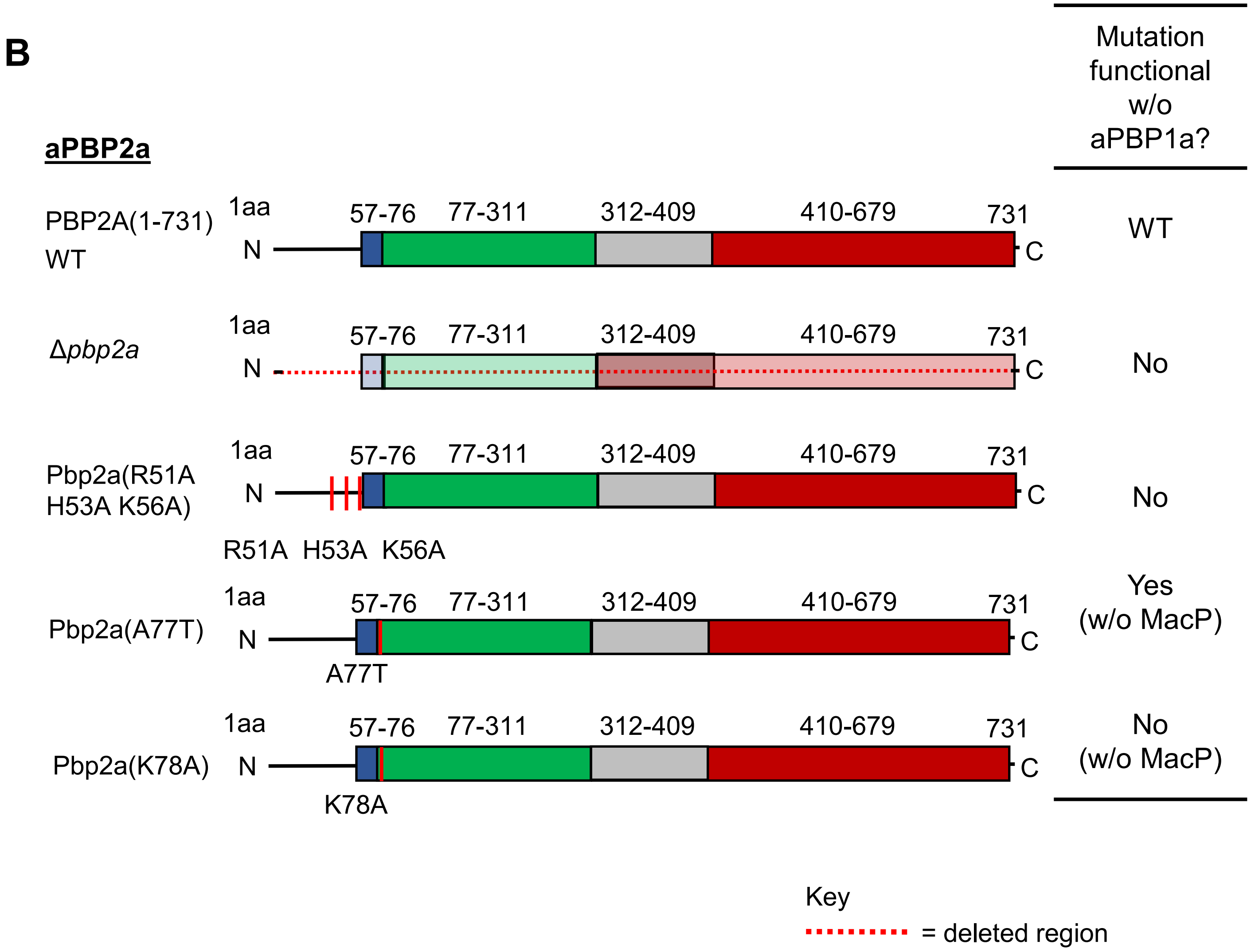


**FIGURE S4 (continued)**

**A**

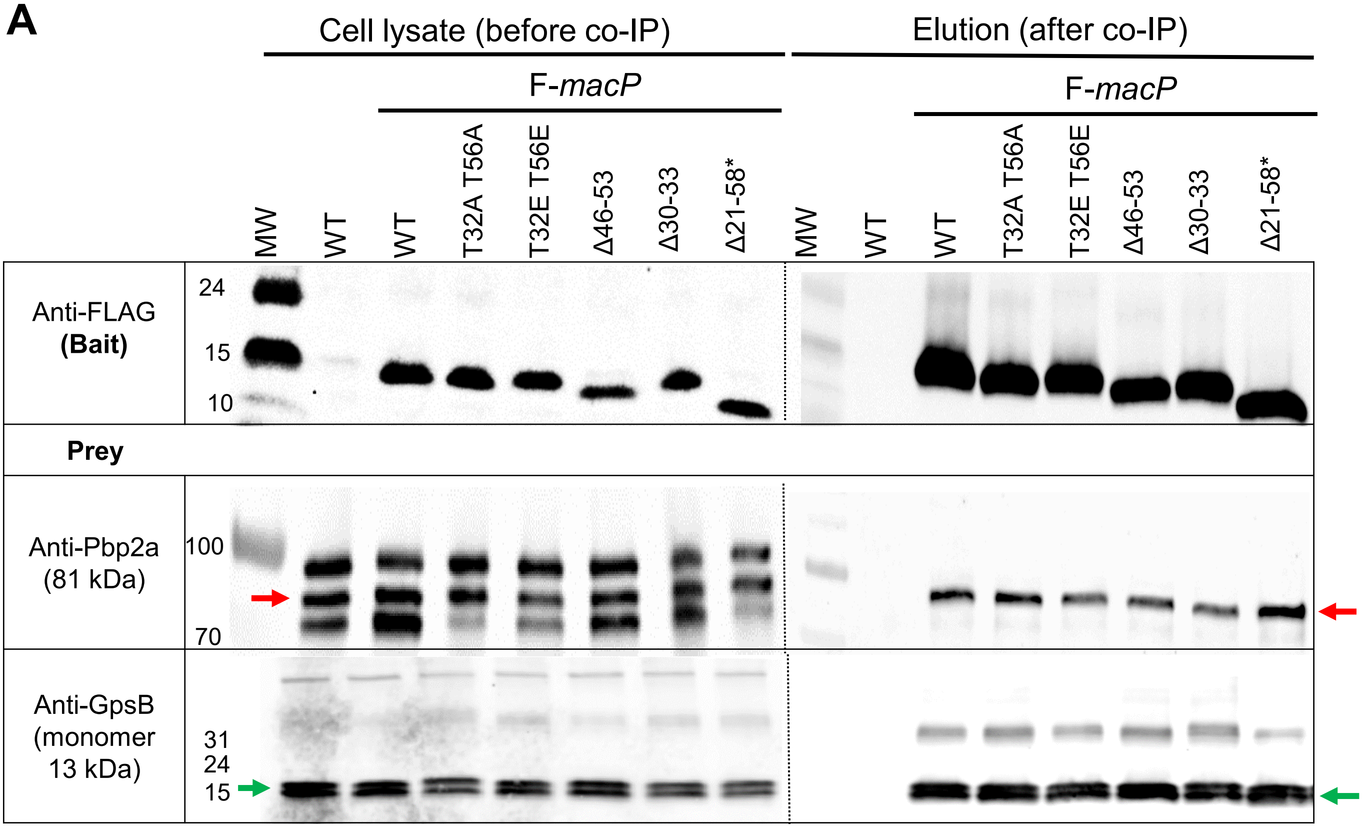


* 2X sample volume was used to obtain the about the same amount of F-MacP

**B**

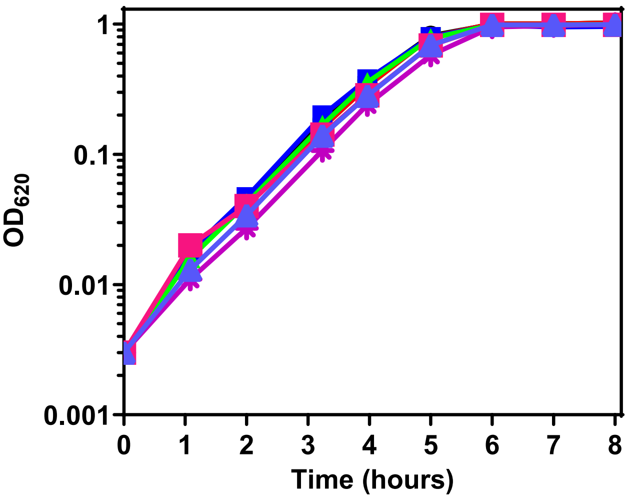


| | Mean ± SEM | |
|---|---|---|
| | **Doubling time (min)** | **Growth yield ($OD_{620}$)** |
| *macP*⁺ WT (1-104) | 41.9 ± 2.0 | 1.00 ± 0.02 |
| Δ*macP* | 31.4 ± 1.9 | 1.01 ± 0.01 |
| *macP*(**Δ5-8**) | 40.6 ± 3.0 | 0.98 ± 0.01 |
| *macP*(**Δ21-58**) | 43.8 ± 1.1 | 1.05 ± 0.03 |
| *macP*(**Δ30-33**) | 37.6 ± 0.2 | 0.90 ± 0.02 |
| *macP*(**Δ46-53**) | 39.6 ± 0.7 | 0.95 ± 0.03 |
| *macP*(**1-77**) | 43.0 ± 1.0 | 1.03 ± 0.02 |
| *macP* (**1-87**) | 41.2 ± 1.0 | 1.00 ± 0.02 |

**Figure S5 (continued)**

C

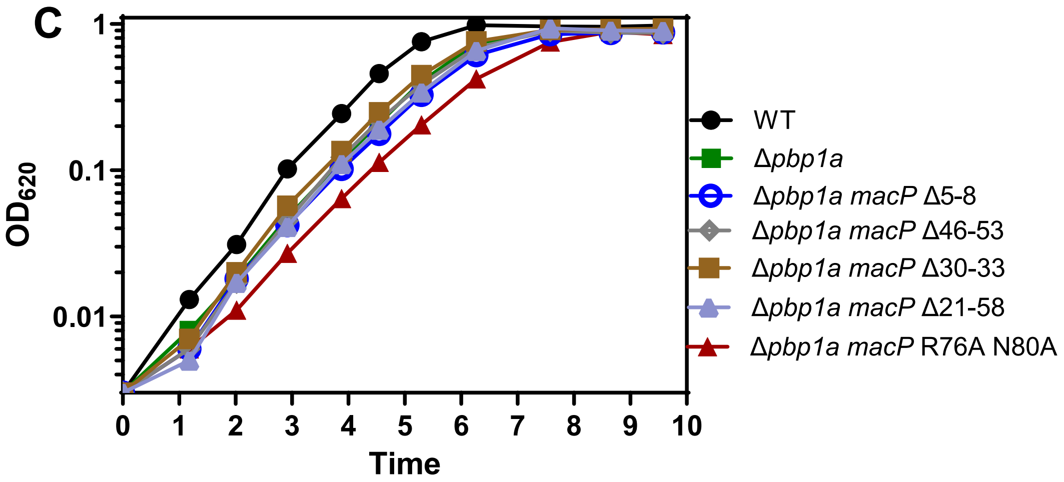


**Mean ± SEM**

| DT (min) | Growth yield ($OD_{620}$) |
|---|---|
| 33 ± 1 | 0.99 ± 0.01 |
| 43 ± 3 | 0.93 ± 0.03 |
| 46 ± 2 | 0.91 ± 0.03 |
| 43 ± 2 | 0.92 ± 0.02 |
| 43 ± 2 | 0.92 ± 0.01 |
| 46 ± 1 | 0.92 ± 0.02 |
| 52 ± 1 | 0.87 ± 0.03 |

D

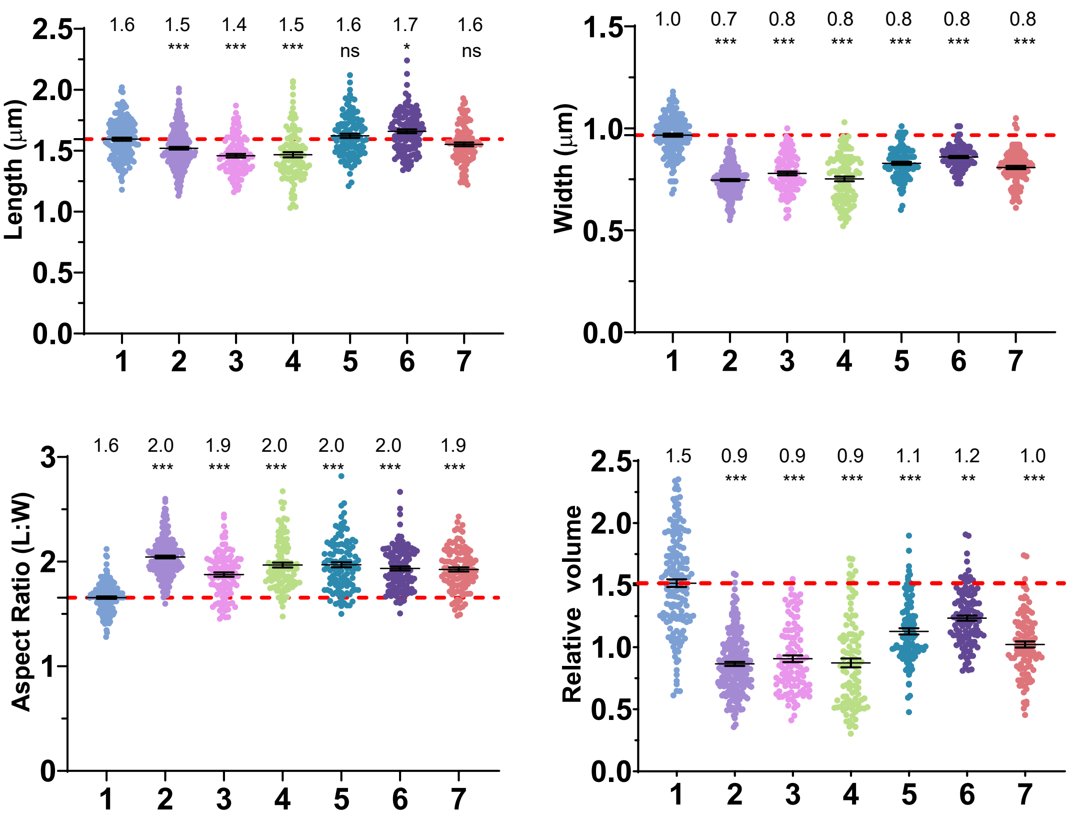


| | Genotype |
|---|---|
| 1 | WT |
| 2 | *Δpbp1a* |
| 3 | *Δpbp1a macP*(Δ5-8) |
| 4 | *Δpbp1a macP*(Δ46-53) |
| 5 | *Δpbp1a macP*(Δ30-33) |
| 6 | *Δpbp1a macP*(Δ21-58) |
| 7 | *Δpbp1a macP*(R76A N80A) |

**FIGURE S5 (continued)**

co-IP- > SDS PAGE -> Silver staining -> MS

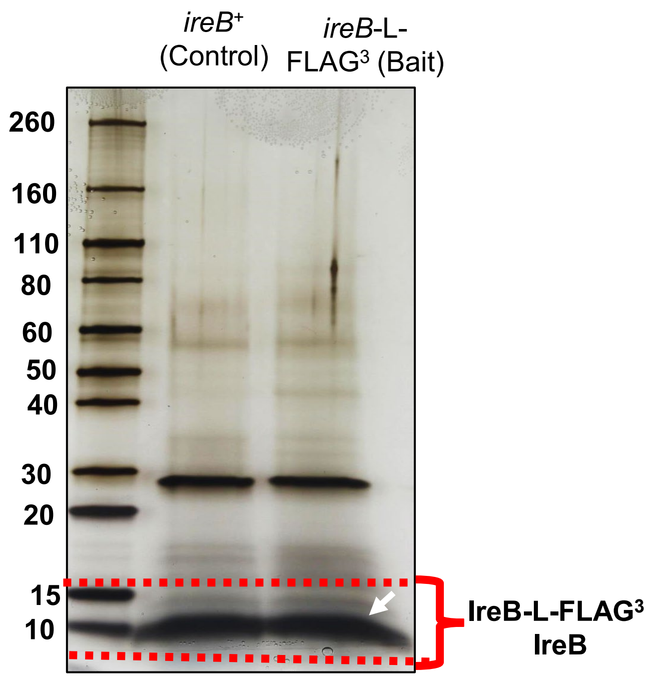


| Mass spectrometry result | | |
|---|---|---|
| Protein | MW (kDa) | Num. Unique Peptides |
| MacP | 13.0 | 13 |

**FIGURE S6**

**A**

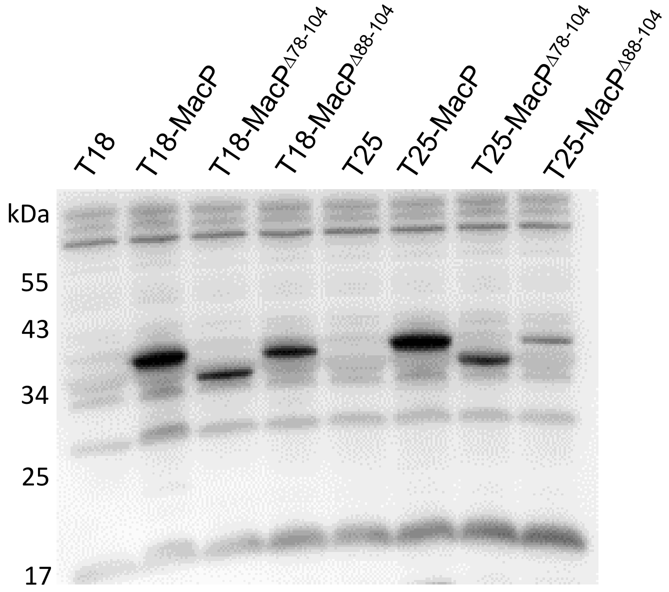


**T18/T25-MacP$^{\Delta 78\text{-}104}$ = T18/T25-MacP 77(stop)**

**T18/T25-MacP$^{\Delta 88\text{-}104}$ = T18/T25-MacP 87(stop)**

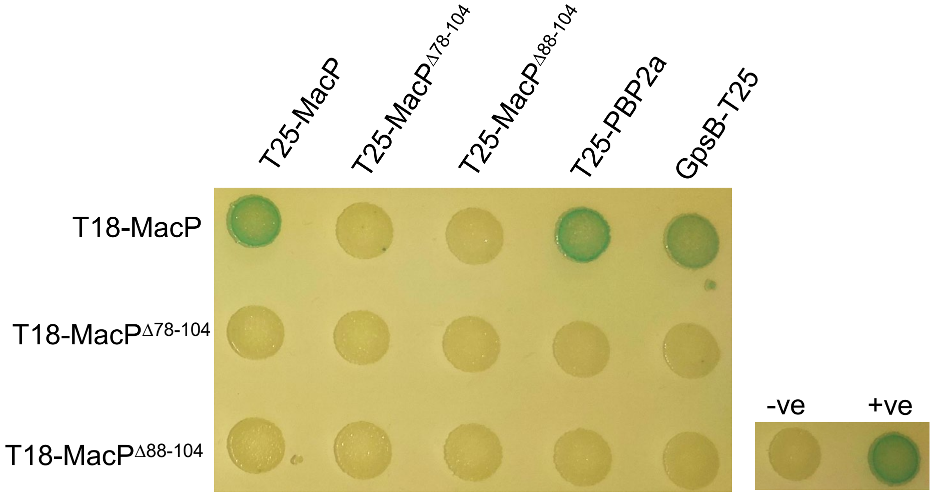


**FIGURE S7 (continued)**

**B**

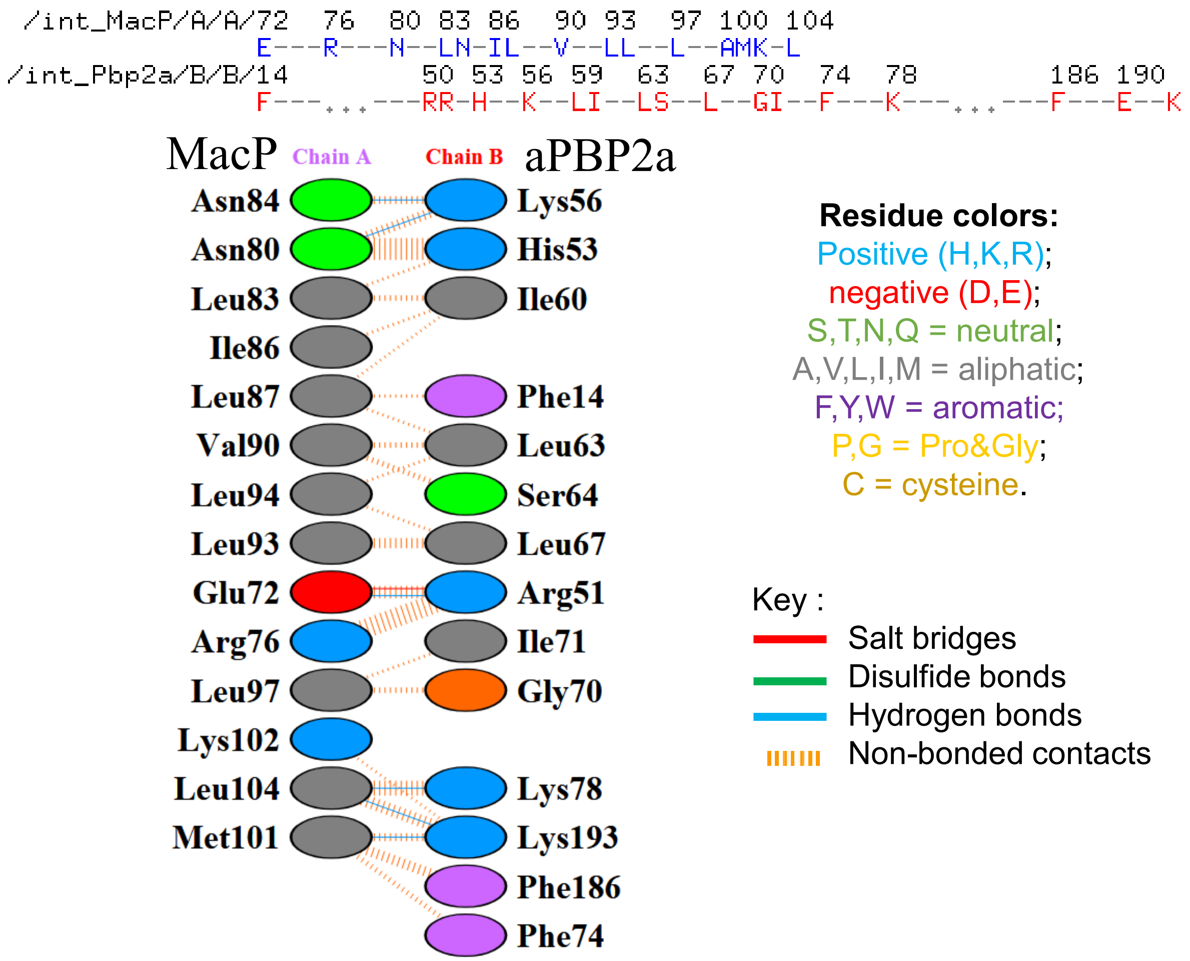


**C**

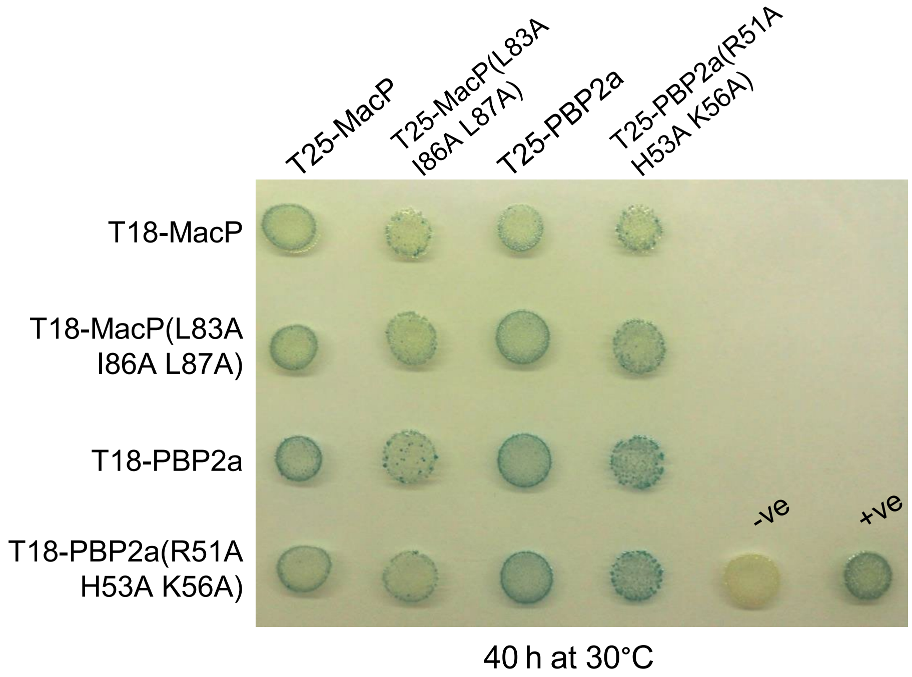


**FIGURE S7 (continued)**

**D**

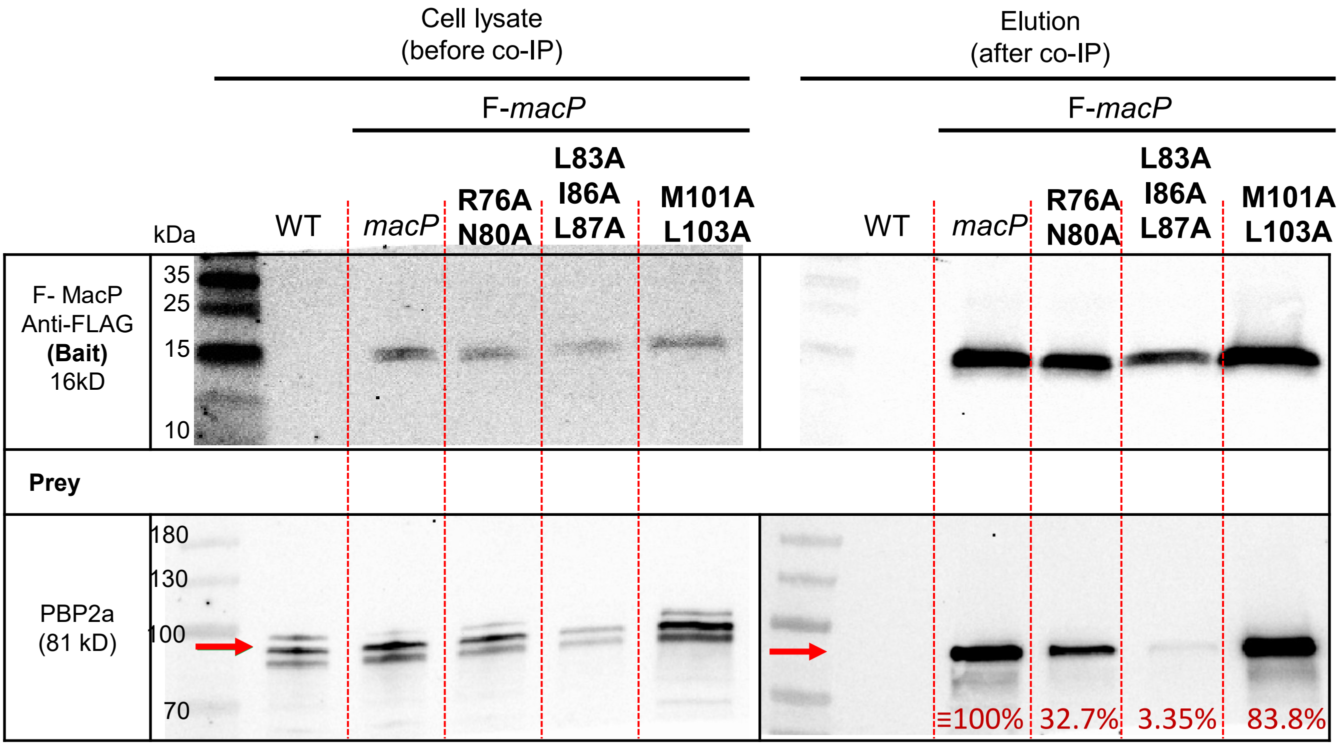


**E**

**Relative amounts of aPBP2a bound to F-MacP variants**

| Genotype | Mean ± SEM |
|---|---|
| | PBP2a |
| WT (IU1824) | 0% |
| F-*macP* (IU17032) | ≡100% |
| F-*macP*(**R76A N80A**) (IU19524) | 20.0 ± 12.7% |
| F-*macP*(**L83A I86A L87A**) (IU19528) | 5.02 ±1.7% |
| F-*macP*(**M101A L103A**) (IU19529) | 67.3 ±16.5% |

**FIGURE S7 (continued)**

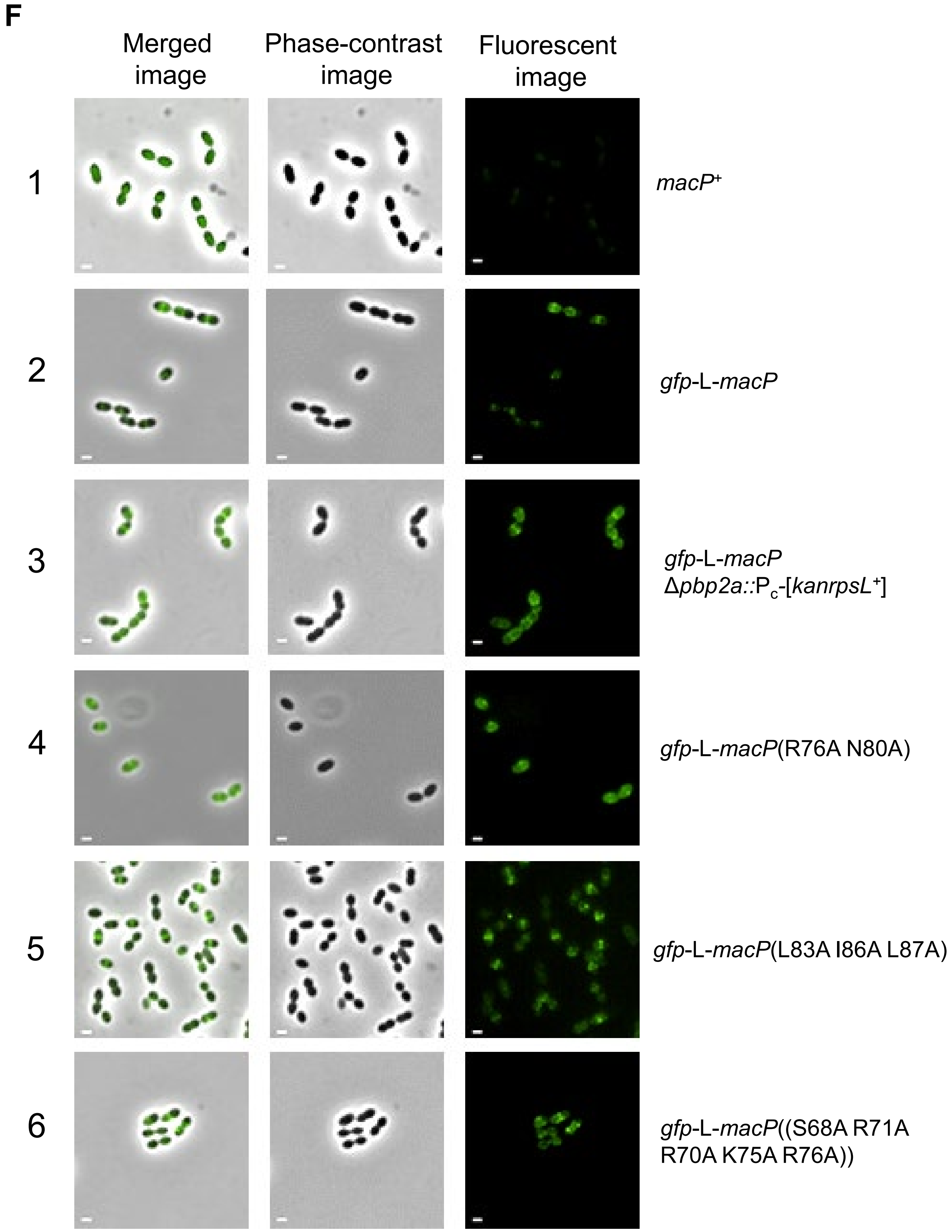


**FIGURE S7 (continued)**

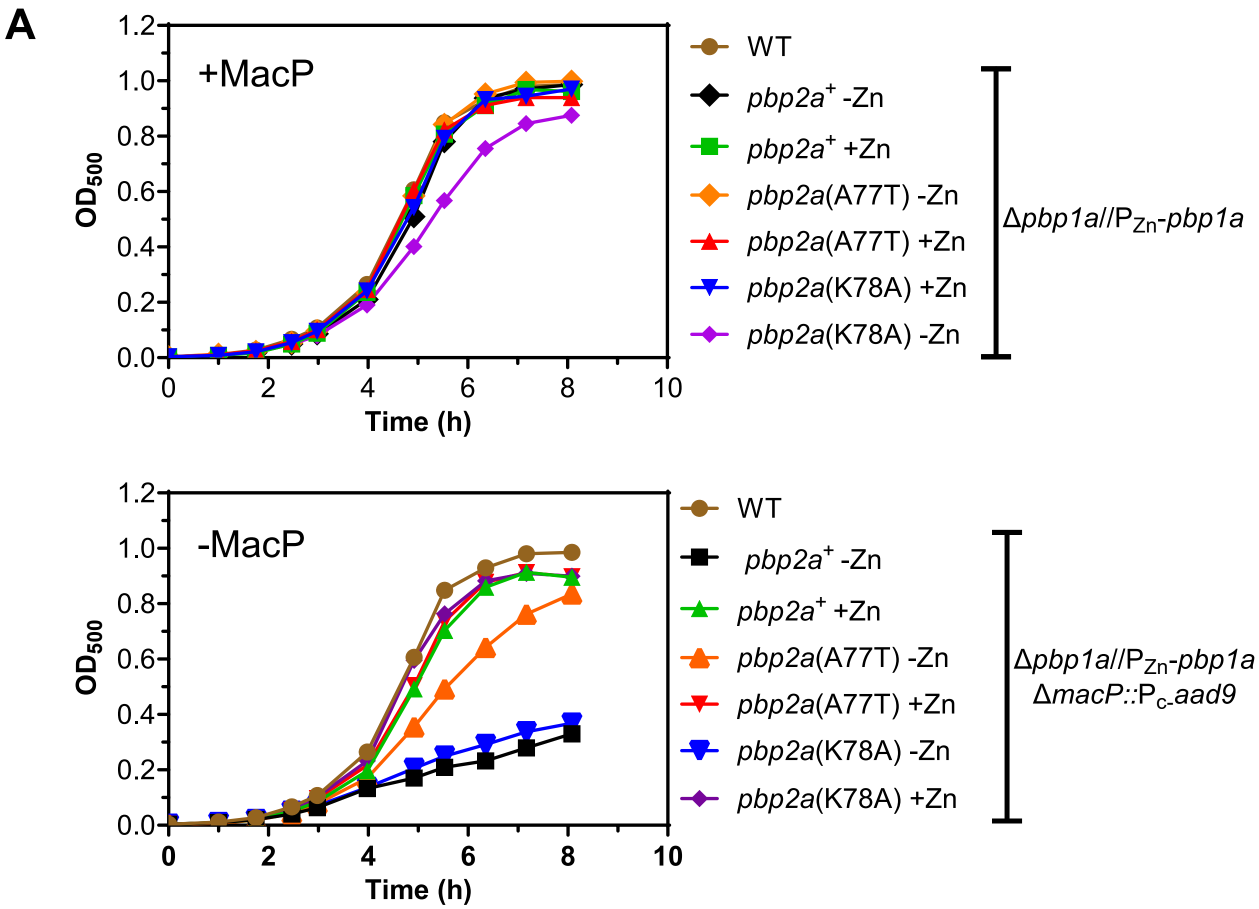


**FIGURE S8 (continued)**

**B**

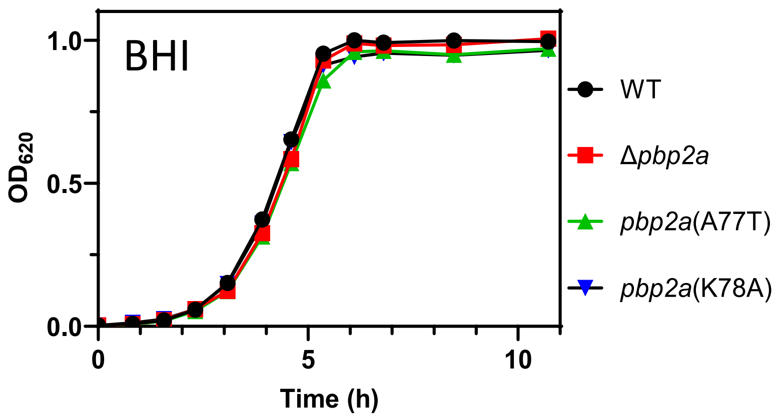


**Mean ± SEM**

| Doubling time (min) | Growth yield ($OD_{620}$) |
|---|---|
| 37.0 ± 0.2 | 0.99 ± 0.002 |
| 38.5 ± 1.3 | 0.99 ± 0.005 |
| 37.8 ± 0.6 | 0.96 ± 0.004 |
| 37.1 ± 1.1 | 0.96 ± 0.005 |

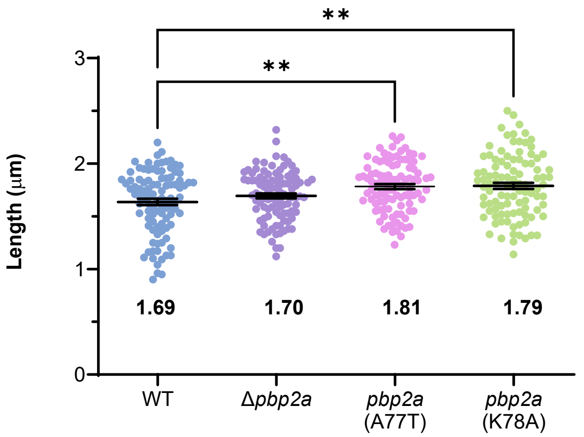


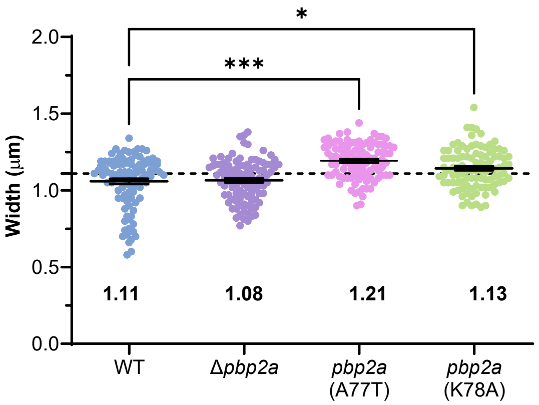


**C**

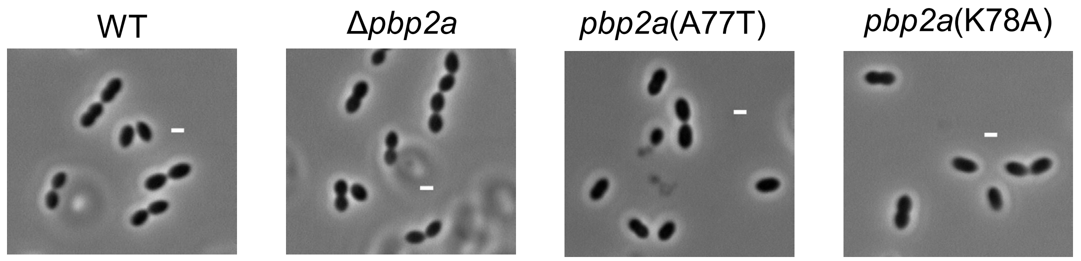


**D**

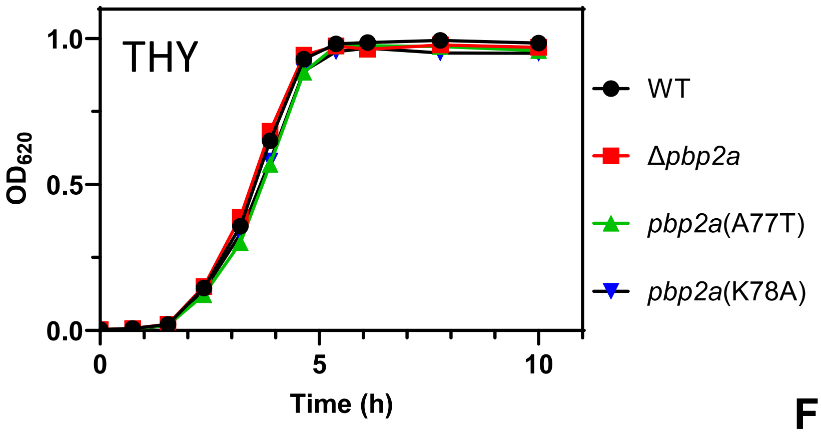


**Mean ± SEM**

| Doubling time (min) | Growth yield ($OD_{620}$) |
|---|---|
| 44.4 ± 0.2 | 0.98 ± 0.004 |
| 44.5 ± 0.6 | 0.97 ± 0.003 |
| 47.0 ± 4.7 | 0.97 ± 0.005 |
| 45.6 ± 0.5 | 0.95 ± 0.006 |

**FIGURE S8 (continued)**

**A**

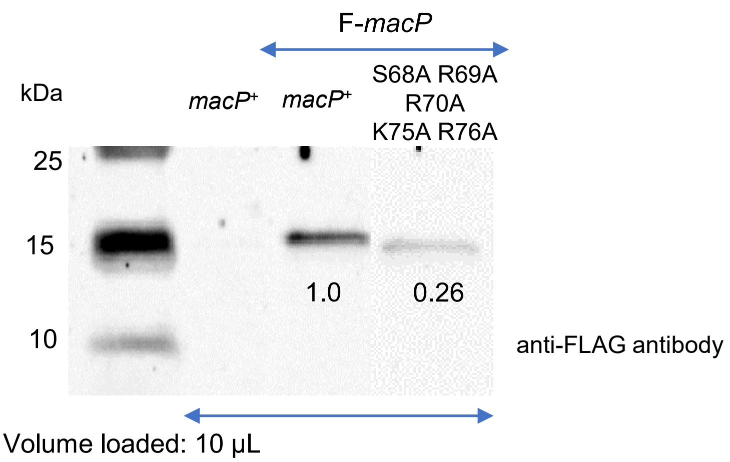


**B**

| Genotype (strain no) | Relative protein amount<br>**Mean ± SEM** |
|---|---|
| *macP*+ (IU1824) | 0 |
| F-*macP* (IU17032) | ≡1.00 |
| F-*macP*(**S68A R69A R70A K75A R76A**) (IU21376 and IU21377) | 0.34 ± 0.04 |

**C**

**Number of colonies 20h after transformation**

| | **Amplicon** | | |
|---|---|---|---|
| Recipient Strain Genotype | Δ*pbp1a*::P$_c$-*erm* | Δ*pbp2a*::P$_c$-*erm* | Δ*bgaA*::P$_c$-*erm* |
| WT (IU1824) | > 500 | > 500 | > 500 |
| *macP*(S68A R69A R70A K75A R76A) (IU21139, IU21140) | > 500 Small/tiny 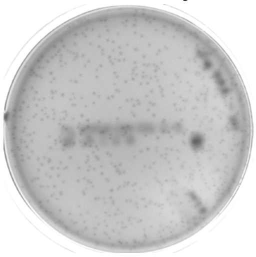 | > 500 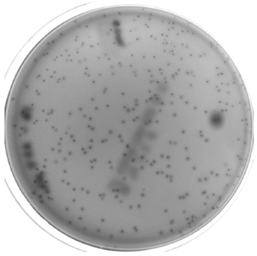 | > 500 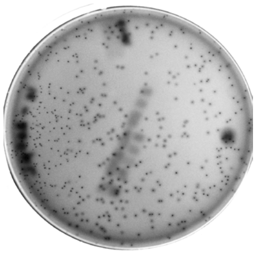 |

*Normalized to 1mL

**FIGURE S9 (continued)**

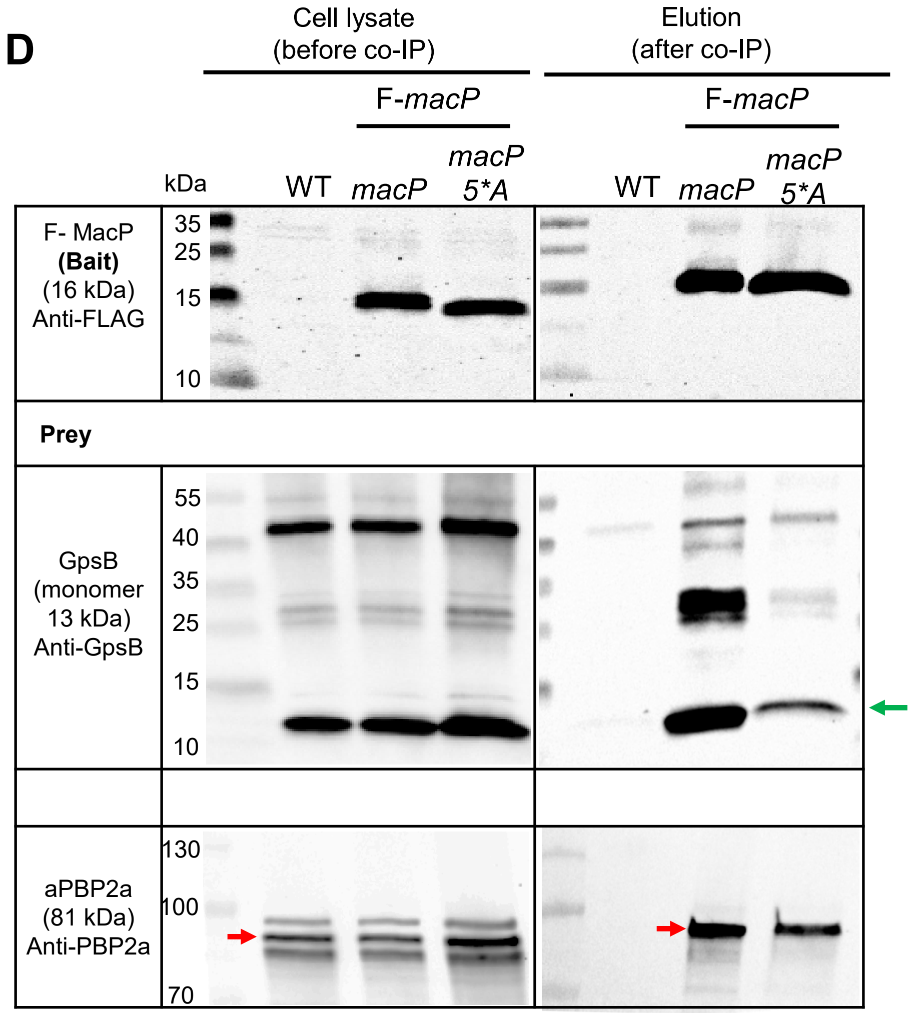


*macP* (5*A) (IU21376) = *macP*(S68A R69A R70A K75A R76A).

(Right panel is repeated from **Figure 8D** for comparison).

The position of GpsB or aPBP2a is marked by a green or red arrow, respectively.

**FIGURE S9 (continued)**

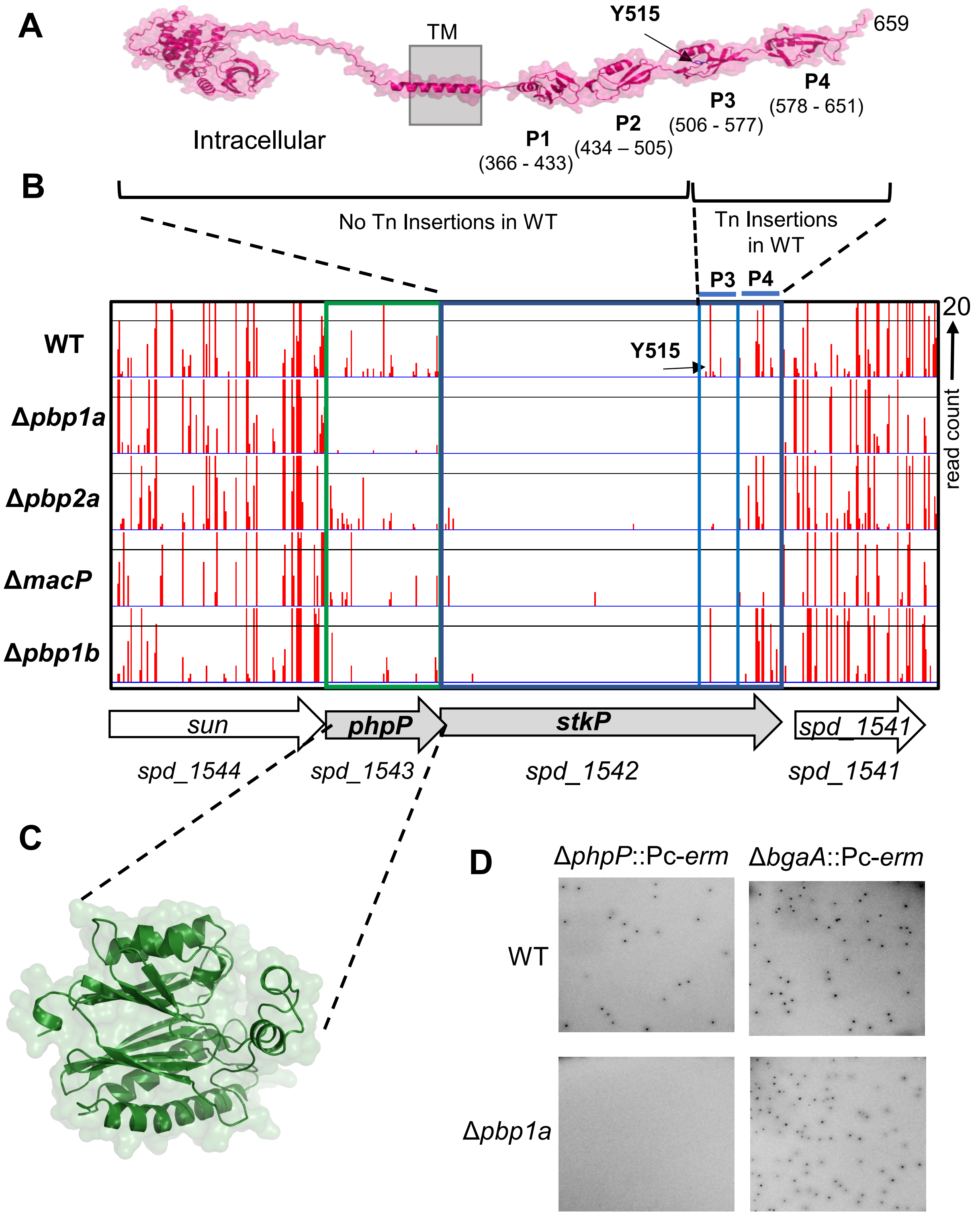


**FIGURE S10 (continued)**

**E**

| | # of colonies and appearance of transformants after 22 h of incubation (# of replicates) | | |
|---|---|---|---|
| | Amplicons | | |
| **Recipient strain** | Δ*pbp1a::*P$_c$-[*kanrpsL*$^+$][a] | Δ*macP* ::P$_c$-*aad9*[b] | Δ*bgaA* ::P$_c$-*aad9*[b] |
| **WT (IU1824)** | >500 normal | >500 normal | >500 normal |
| ***murZ*(D280Y) Δ*stkP* (IU16885)** | >500 small | >500 normal | >500 normal |
| ***murZ*(D280Y) Δ*stkP* Δ*pbp1a* (IU19908)** | N/A | 0 | >500 tiny |

[a]Selection with kanamycin

[b]Selection with spectinomycin

**FIGURE S10 (continued)**

**A**

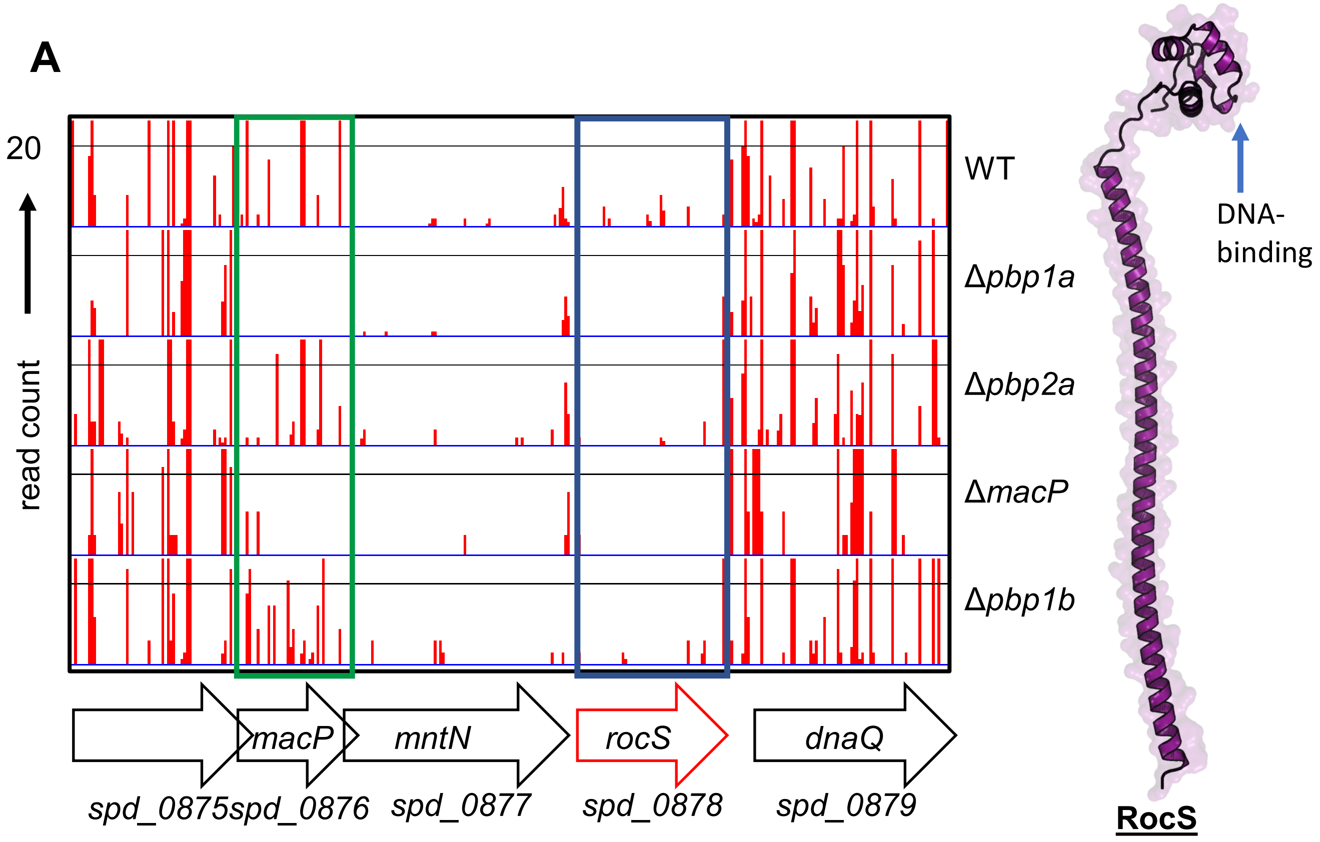


**B**

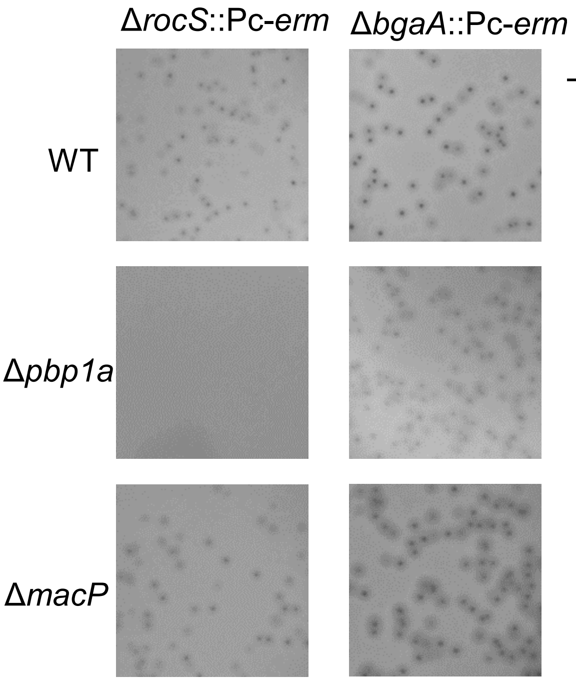


| # of colonies and appearance of transformants after 22 h of incubation (# of replicates) | | | |
|---|---|---|---|
| **Recipient strain** | Amplicons | | |
| | ΔrocS | ΔbgaA | Δpbp1b |
| **WT (IU1824)** | >500 small (2) | >500 normal (>3) | >500 normal (2) |
| ***Δpbp1a* (IU18579)** | 0 (2) | >500 normal (4) | >500 small (2) |
| ***Δpbp2a* (IU13256)** | >500 small (2) | >500 normal (2) | >500 normal (2) |
| ***ΔmacP* (IU14699)** | >500 small (2) | >500 normal (4) | >500 small (1) |
| ***Δpbp1b* (IU14697)** | >500 small (2) | >500 normal (1) | >500 normal (1) |

**FIGURE S11**
